# A portable dynamic loop module enables phosphotriesterase function in cysteine-dependent hydrolases

**DOI:** 10.1101/2025.10.30.685675

**Authors:** J. David Schnettler, Eleanor Campbell, Rashid Khashiev, Oskar James Klein, Florian Hollfelder

**Author notes:** Correspondence to: Florian Hollfelder. Bioprocess Laboratory, Department of Biosystems Science and Engineering, ETH Zurich, Klingelbergstrasse 48, 4056 Basel, Switzerland. Australian Synchrotron, 800 Blackburn Rd, Clayton VIC 3168, Australia. Department of Chemical and Biological Engineering, University of Sheffield, Mappin Street, S1 3JD, Sheffield, United Kingdom. Manchester Institute of Biotechnology, University of Manchester, 131 Princess Street, M1 7DN, Manchester, United Kingdom.

## Abstract

A large disparity has emerged between the availability of massive metagenomic DNA sequence data and the actual functional annotation of enzyme catalysts, their mechanistic classification and insight into catalytic stragies. Exploration by sequence homology is intrinsically conservative, but ultrahigh-throughput functional metagenomics offers the chance to ‘jump’ into unknown sequence space to provide novel catalysts without precedent. Having identified a novel metal-free phosphotriesterase with an active site cysteine-containing triad, here, we launch an exploration of sequence-structure-function relationships of a range of related proteins from the dienelactone hydrolase (DLH) family, revealing 10 new phosphotriesterases. Four crystal structures provide clues to mechanism, suggesting – based on phylogenetic and structural analysis – that phosphotriesterase activity is mediated by loops surrounding the active site with activity changes over 4 orders of magnitude correlated to loop flexibility. These insights allow protein engineering by loop grafting across family members, resulting in 8-fold increased phosphotriesterase activity in a human enzyme. We thus demonstrate that identification of a single ‘pioneer’ enzyme can provide starting points for activity engineering strategy yielding new reagents for bioremediation or treatment of organophosphate poisoning.

## Introduction

The biosphere is replete with functional molecules, yet the overwhelming majority remains undiscovered. Our current knowledge of such biomolecules, particularly of functional proteins, is strongly biased by their history of discovery and limited by the experimental effort expended on understanding their structure and function. The enormous combinatorial diversity of protein sequences has led to the concept of a ‘vast’ sequence space (1). Indeed, metagenomic databases such as MGnify which (as of April 2024) contains over 2.4 billion open reading frames encoding diverse proteins with specific functions (2), but only a few tens of thousand proteins are experimentally characterised. This is why assigning function to novel sequences and elucidating their underlying catalytic strategies and mechanisms remains a major challenge. While sequence-based homology searches are commonly used, they are limited to our – often sparse – existing knowledge (i.e., merely assign the function of the closest experimentally characterised homologue) but fail to discover unsuspected functions with yet unknown underlying mechanisms. Further, this approach is blind for the hidden ‘promiscuous potential’ of the proteome (3, 4). Therefore, to uncover novel functions, we need additional experimental annotation in less-characterised regions of sequence space.

The need for new functional proteins is exemplified by the degradation of man-made compounds without previous prevalence on Earth, such as phosphotriester pesticides. This class of compounds has entered the environment only recently through human pollution, starting in the late 1940s. Within a relatively brief timeframe (compared to an evolutionary scale), enzymes capable of hydrolysing these toxic compounds have rapidly emerged in less than a century. All of them have evolved from native hydrolases with a promiscuous phosphotriesterase side activity, even though an identical catalytic feature – a catalytic metal cofactor – has been shown to be repurposed in unrelated protein superfamilies: the amidohydrolase (5), the pita-bread (6), the beta-propeller (7, 8), the metallo-beta-lactamase (9), and the cyclase (10) superfamilies.

Given that the vastness of sequence space is still untested for phosphotriesterase function, it is conceivable that there are alternative starting points for engineering enzymes against organophosphate poisoning that may also reveal novel mechanistic features. By carrying out an ultrahigh-throughput function screen of metagenomic DNA, we previously provided evidence for this hypothesis. We isolated a hitherto unknown phosphotriesterase, P91, from the alpha/beta-hydrolase superfamily, where – until then – no such activity had been detected (11). P91 broke new mechanistic ground, because it does not – in contrast to all other triesterases – require a metal cofactor. Instead, the active site harbours a catalytic triad with a cysteine nucleophile that forms a covalent intermediate, but also enables multiple turnover. This stands in contrast to the physiological target of phosphotriesters, the enzyme acetylcholinesterase, where a serine-containing catalytic triad is inactivated by suicide inhibition when reacting with a phosphotriester. We subsequently evolved P91 to reach activities similar to those of metal-containing phosphotriesterases (12), establishing it as a potentially useful alternative for bioremediation.

The discovery of P91 and its evolution to a proficient enzyme with a second order rate acceleration of 10^14^-fold raises questions about its wider context and occurrence, prompting an examination of whether this activity is primary (native) or secondary (promiscuous). In addition, this novel bridgehead in sequence space calls for a classification that correctly identifies and annotates additional phosphotriesterases. By sequence similarity annotation, P91 belongs to the dienelactone hydrolase (DLH) family proteins (Pfam: PF01738) within the alpha/beta hydrolase superfamily (Pfam: CL0028, Figure **1a**). However, P91’s annotation as dienelactone hydrolase is merely based on the functional and structural characterisation of a single enzyme, the dienelactone hydrolase from *Pseudomonas knackmussii (Pk*DLH) (13). Only three further enzymes from the family of DLH-like proteins have been functionally characterised (apart from P91): one has been shown to act on the same dienelactone substrate as *Pk*DLH (14) but two others act on different substrates, a carboxyester (15) and an organic acid anhydride (16), providing evidence that the current annotation is not unambiguously predictive and does not account for the phosphotriesterase activity of family member P91.

**Figure 1:**
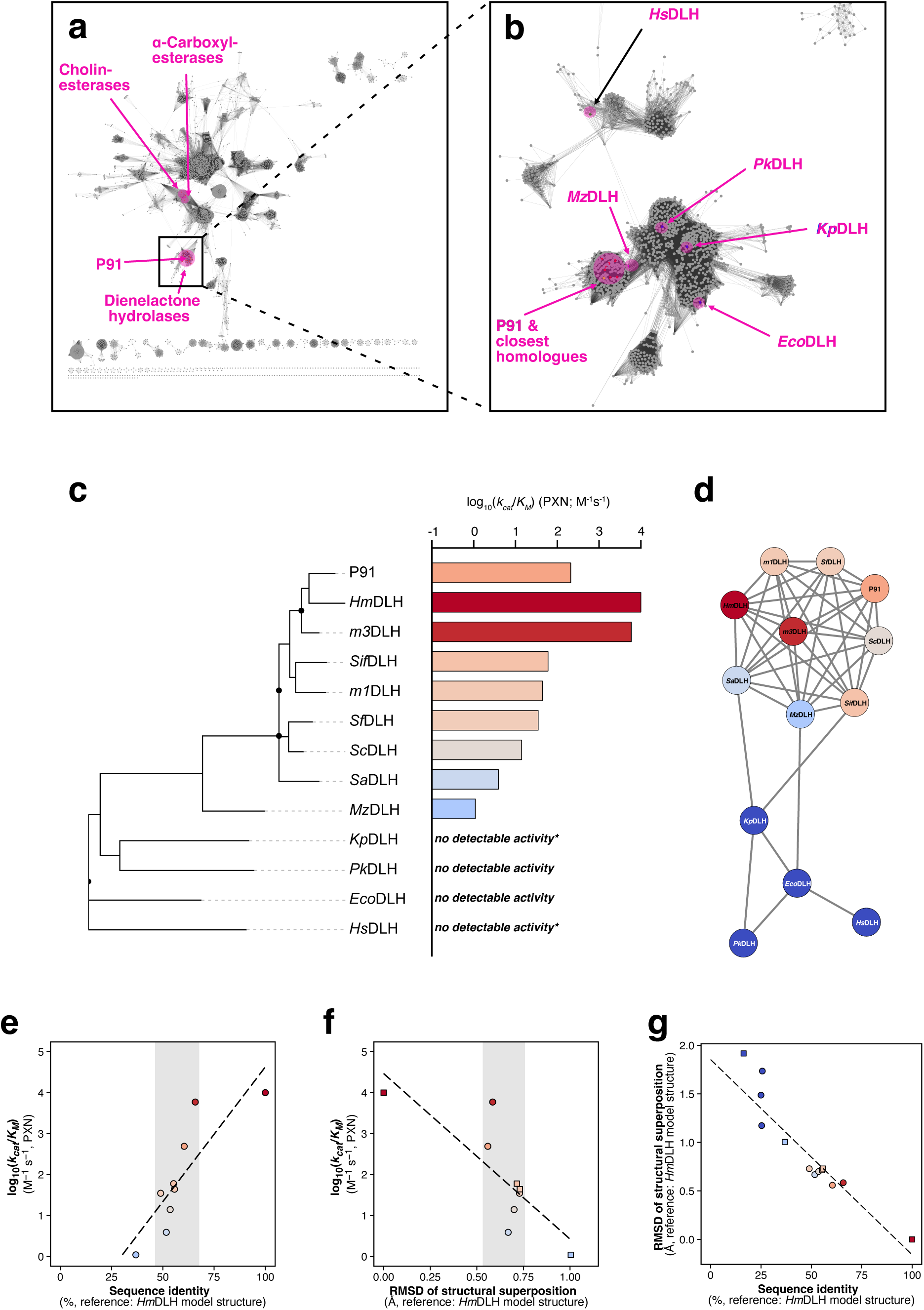
Phosphotriesterase activity is widespread in the DLH protein family. **(a)** Sequence similarity network (SSN) of the alpha/beta hydrolase superfamily (Pfam: CL0028) containing 5062 sequences. Nodes with an alignment score ≥ 12 are connected by an edge, corresponding to a median sequence identity of ≈ 27 %. **(b)** A separate SSN of the dienelactone hydrolase (DLH) protein family (Pfam: PF01738), containing 1992 sequences. Nodes with an alignment score ≥ 25 are connected by an edge, corresponding to a median sequence identity of ≈ 32 %. In this SSN, separate small clusters and unconnected nodes (singletons) were removed for clarity. All enzymes characterised in this study are highlighted: P91 and its closest homologues, an enzyme identified by sequence similarity of a neighbouring gene fragment (*Mz*DLH), DLH family proteins with a publicly available structure (*Pk*DLH, *Eco*DLH, *Kp*DLH), and the only homologue of P91 in the human genome (*Hs*DLH). **(c)** Phylogenetic tree and catalytic efficiencies for phosphotriester hydrolysis (log_10_(*k_cat_*/*K_M_*)) of all tested DLH family proteins. Bars are coloured according to *k_cat_/K_M_* for paraoxon hydrolysis in a colour gradient from blue (no detectable activity) to red (high activity).. For four enzymes (*Kp*DLH, *Pk*DLH, *Eco*DLH, *Hs*DLH), no activity towards paraoxon could be detected. However, *Kp*DLH and *Hs*DLH (marked with an asterisk) showed activity towards the phosphotriester FDDEP. **(d)** Sequence similarity network of all tested DLH family proteins. Nodes with an alignment score ≥ 12 are connected by an edge. Nodes are coloured according to their phosphotriesterase activity (log_10_(*k_cat_*/*K_M_*)) for paraoxon) from blue (no detectable activity) to red (high activity). **(e–g)** Sequence-structure-function map of promiscuous phosphotriesterase activity in the DLH protein family. Sequence identity and structural similarity of all studied enzymes correlate with levels of promiscuous phosphotriesterase activity, suggesting a sequence-structure-function trajectory (dotted line). However, within a narrow range of sequence identity and structural similarity to highly active phosphotriesterases, promiscuous phosphotriesterase activity spans over five orders of magnitude (grey areas). Plots show the relationship of **(e)** catalytic efficiency towards the phosphotriester paraoxon over sequency identity, **(f)** catalytic efficiency over root mean square deviation (RMSD) of structural superpositions, and **(g)** sequence identity over RMSD of structural superpositions. In all comparisons, the enzyme with the highest phosphotriesterase activity, *Hm*DLH, was used as the point of reference. Where no experimental structure is available, structural models created with AlphaFold2/ColabFold (18, 19) were used (square markers).

Here we experimentally explore the distribution of this recently discovered phosphotriesterase activity within the DLH family. We expand the annotation for this promiscuous activity by identifying 10 new phosphotriesterases, four of which were also crystallised and structurally characterised. These experiments provide a framework for tracking the sequence-structure-function relationships in the DLH family, illuminate the regions in sequence space where promiscuous phosphotriesterase activity occurs and identify a portable dynamic feature that enhances catalytic activity.

## Results

### Latent phosphotriesterase activity is widespread among DLH family enzymes

P91’s newly identified promiscuous phosphotriesterase activity raised the question whether phosphate triester hydrolysis is a constant side activity of related enzymes. P91 had been assigned by sequence homology as a member of the dienelactone hydrolase (DLH) protein family, a subclass of the alpha/beta hydrolase superfamily (Pfam clan CL0028) (17). Promiscuous activities are difficult to predict, even when similar active-site architectures persist in a protein family. We thus set out to test experimentally for phosphotriester hydrolysis, in order to add annotations for this function to DLHs, and map where this recently discovered activity can be found across members of the DLH family to derive structure-activity relationships.

To define a test panel of candidate proteins that spans across the sequence space of the DLH protein family (**Figure 1b**), we explored increasingly distant homologues, to reach as little as 27% sequence identity to P91. Seven family members with high homology to P91 (65–56%) were derived from the databases GenBank (*Hm*DLH, *Sa*DLH, *Sc*DLH, *Sf*DLH, *Sif*DLH), marDB (*m3*DLH) and MGnify (*m1*DLH) (**Table S1**). These sequences originate from aquatic gammaproteobacteria (genera *Haliea*, *Solimonas*, and *Sinimarinibacterium*) or are metagenomic sequences of aquatic origin. A homologue from *Marinobacter zhejiangensis* (*Mz*DLH) has lower sequence identity to P91 (27 %) but has a similar adjacent gene in its genomic context, possibly implying a similar native function as P91 (**Figure S2**). In addition, we sought to test all DLH family proteins of which a crystal structure is available and thus expressed and purified P91 homologues from *Pseudomonas knackmussii* (*Pk*DLH), *Escherichia coli* (*Eco*DLH), and *Klebsiella pneumoniae* (*Kp*DLH). Furthermore, we also included P91’s homologue from the human genome (*Hs*DLH, previously referred to as CMBL (15)) into the panel (**Table S1**).

We tested the activity of these 12 enzymes towards the phosphotriester substrate paraoxon (2). Most (8 out of 12) of the DLH family members showed detectable activity against this substrate. Of the four enzymes without detectable activity, two (*Kp*DLH and *Hs*DLH) showed activity towards the slightly more reactive phosphotriester substrate FDDEP (leaving group p*K_a_* 6.4 for fluorescein in contrast to p*K_a_* 7.1 for 4-nitrophenylate) and only 2 out of 12 entirely lacked detectable promiscuous activity (*Pk*DLH and *Eco*DLH). Reaction progress curves (**Figure S3**) followed a single exponential product increase, with no visible burst kinetics on a timescale of minutes to hours. We used these time courses to derive Michaelis-Menten parameters for each enzyme (**Figures 1c, S4, and S5**, **Table S2**). While previously no phosphotriesterase activity had been assigned in the alpha/beta hydrolase superfamily (except for P91), this annotation now extends to eight members of the DLH subfamily.

Mapping phosphotriester activity to sequence identity and structural similarity (**Figure 1e–g**) shows a decrease in activity with decreasing homology to P91 (**Figure 1c)**. However, this sequence-activity correlation shows some scatter: in a close cluster of homologues large differences in activity (≈ 2600-fold) between highly homologous enzymes can be observed (e.g., *Sa*DLH with *k_cat_*/*K_M_* = 3.9 M^-1^s^-1^ and *Hm*DLH with *k_cat_*/*K_M_* = 10000 M^-1^s^-1^, grey areas in **Figure 1e, f** and **Figure 1d**), suggesting that phosphotriesterase activity is determined by local rather than global enzyme properties (e.g., overall fold). We therefore initially hypothesised that latent phosphotriesterase activity is determined by specific activity-conferring mutations.

### Crystal structures reveal a shared topology and active-site features but no determinants of phosphotriesterase activity

We obtained crystal structures for *m3*DLH, *Sa*DLH, *Sc*DLH, *Sf*DLH and reveal that they share the same structural topology as the canonical dienelactone hydrolase *Pk*DLH, a core beta-sheet consisting of eight strands surrounded by several alpha-helices (**Figure 2a**). Analogous to cysteine proteases (such as papain) they share a catalytic Cys-His-Asp triad, albeit in a mirrored orientation, implying a catalytic mechanism for phosphotriester hydrolysis that involves a covalent intermediate (**Figure 2b**), as previously shown in detail for P91(11, 12). In P91, the active-site cysteine is present in two alternative conformations, one pointing towards the active site, and one pointing inwards, possibly to be protected from oxidation(11).

**Figure 2:**
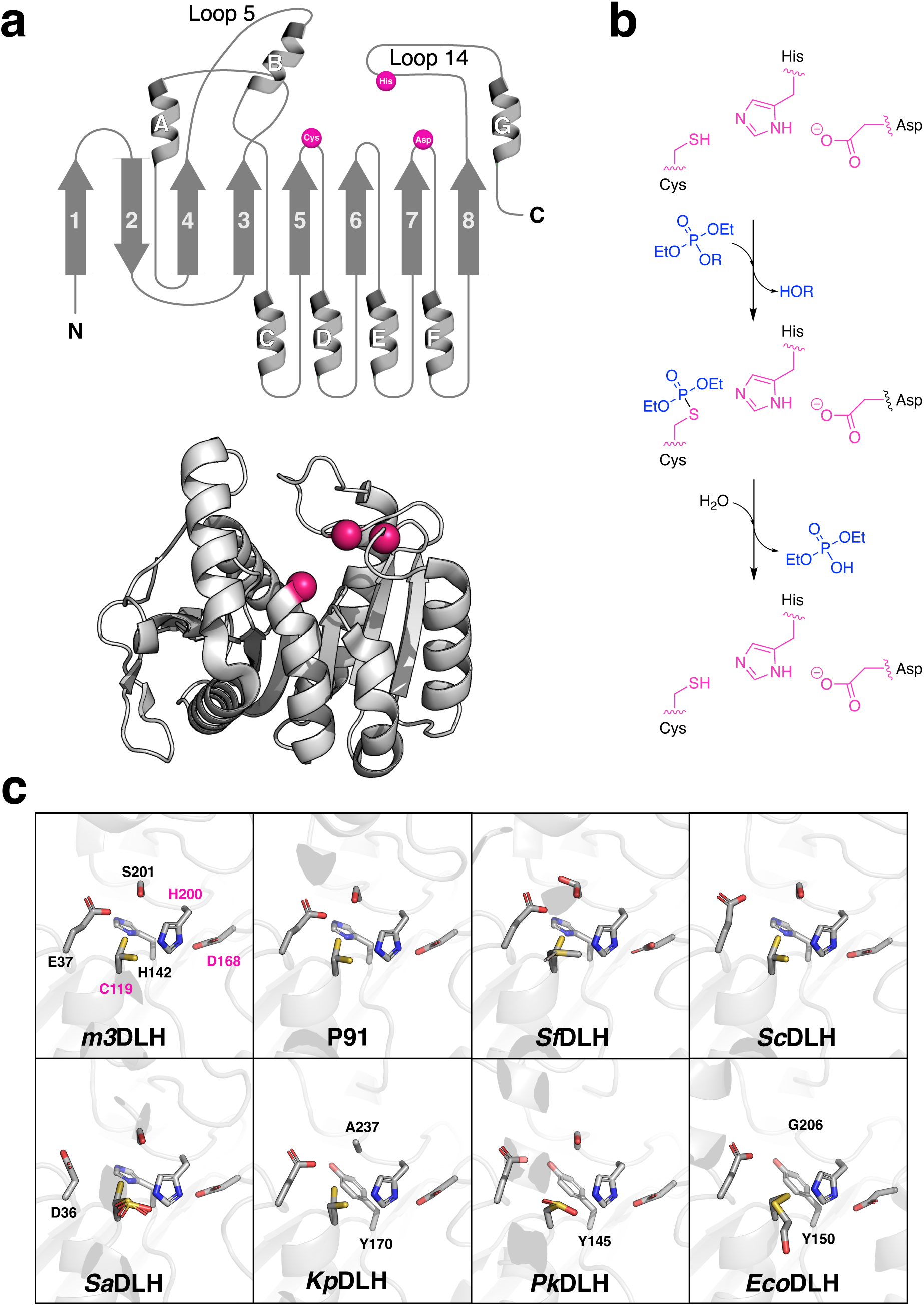
DLH family proteins share a common structural topology, a Cys-His-Asp catalytic triad, and a phosphotriesterase mechanism involving a covalent intermediate. **(a)** All proteins studied in this work share the same structural topology which is archetypical for the alpha/beta hydrolase superfamily: A core beta-sheet consisting of eight strands is surrounded by several alpha-helices. The catalytic Cys-His-Asp triad (magenta spheres) is located in topologically conserved positions, exemplified here with the structure of *Pk*DLH. The connecting loops between secondary structure elements are variable in length, with loops 5 and 14 partly covering the active site. **(b)** DLH family proteins hydrolyse phosphotriesters (blue) via a catalytic mechanism analogous to cysteine proteases, involving the formation of a covalent intermediate between the active-site cysteine (magenta) and the substrate. **(c)** Catalytic triad and ‘extended hexad’ of all DLH family with a known crystal structure ordered by decreasing activity. In P91, the active-site cysteine is present in two conformations: One pointing towards the active site, and one pointing inwards (stabilised by three further residues E37, H142, and S201 in the case of *m3*DLH). We also observe these two conformations in crystal structures of *m3*DLH, *Sf*DLH, *Sc*DLH, *Sa*DLH, and *Kp*DLH. Further, we observe covalent modification or oxidation of the active site cysteine in structures of *Sf*DLH, *Sa*DLH, *Pk*DLH, and *Eco*DLH. The catalytic triad is highlighted in magenta. Residues are labelled for *m3*DLH and in other enzymes only where they deviate from the consensus.

To resolve the determinants of phosphotriesterase activity, we compared the principal active site features, including the three residues that stabilise the inwards-pointing conformation of the active-site cysteine, which we termed the ‘extended hexad’ (**Figure 2c**). The extended hexad is identical in all homologues with high phosphotriesterase activity (P91, *Hm*DLH, *m3*DLH, *Sif*DLH, *Sf*DLH, *Sc*DLH) but deviates in homologues with very low (*Sa*DLH, *Mz*DLH, *Hs*DLH, *Kp*DLH) or no detectable activity (*Pk*DLH, *Eco*DLH), possibly indicating an involvement in catalysis. However, mutating the hexad towards the sequence found in active phosphotriesterases did not increase activity in inactive phsophotriesterases.

To extend the range of active-site residues we surveyed the surroundings of the active site and selected 20 residues that were identical in all active homologues and differed in inactive homologues. We then devised a combinatorial library based on the inactive enzyme *Pk*DLH in which each of these residues could either be wild-type or according to the active-enzyme consensus. However, complete screening of this million-membered library in microfluidic droplets did not identify any enzyme with detectable phosphotriesterase activity (see **Supplementary Information, section 1.4**).

### Lid loop flexibility correlates with promiscuous phosphotriesterase activity

After the previously successful approach (12) of assigning phosphotriesterase function to active-site residue arrangements failed, we focussed on the differences in structural topology. Comparison of all available structures revealed that the main differences lie in the length and conformational diversity of two loops that shape the access to the active site, termed loop 5 and loop 14 (**Figure 3a, Figure S15**). Length and normalised B-factors (B’) of these loops loosely correlate with phosphotriesterase activity (**Figure S16**). Analysis of the B’-factor profile of active homologues revealed that B-factor values show local maxima in the regions of loop 14 and the transition of helix B and loop 5, which form the ‘lid’ domain shaping the entrance of the active site (**Figure 3b**). In inactive homologues, however, this pattern is completely absent (**Figure 3c**). Averaging the normalised B-factor for each of these sections and for every protein reveals a clear correlation between phosphotriesterase activity level and lid domain flexibility (**Figure S17**). This correlation of conformational diversity in loops 5 and 14 with promiscuous phosphotriesterase activity is further supported by ensemble refinement (**Figure 3**).

**Figure 3:**
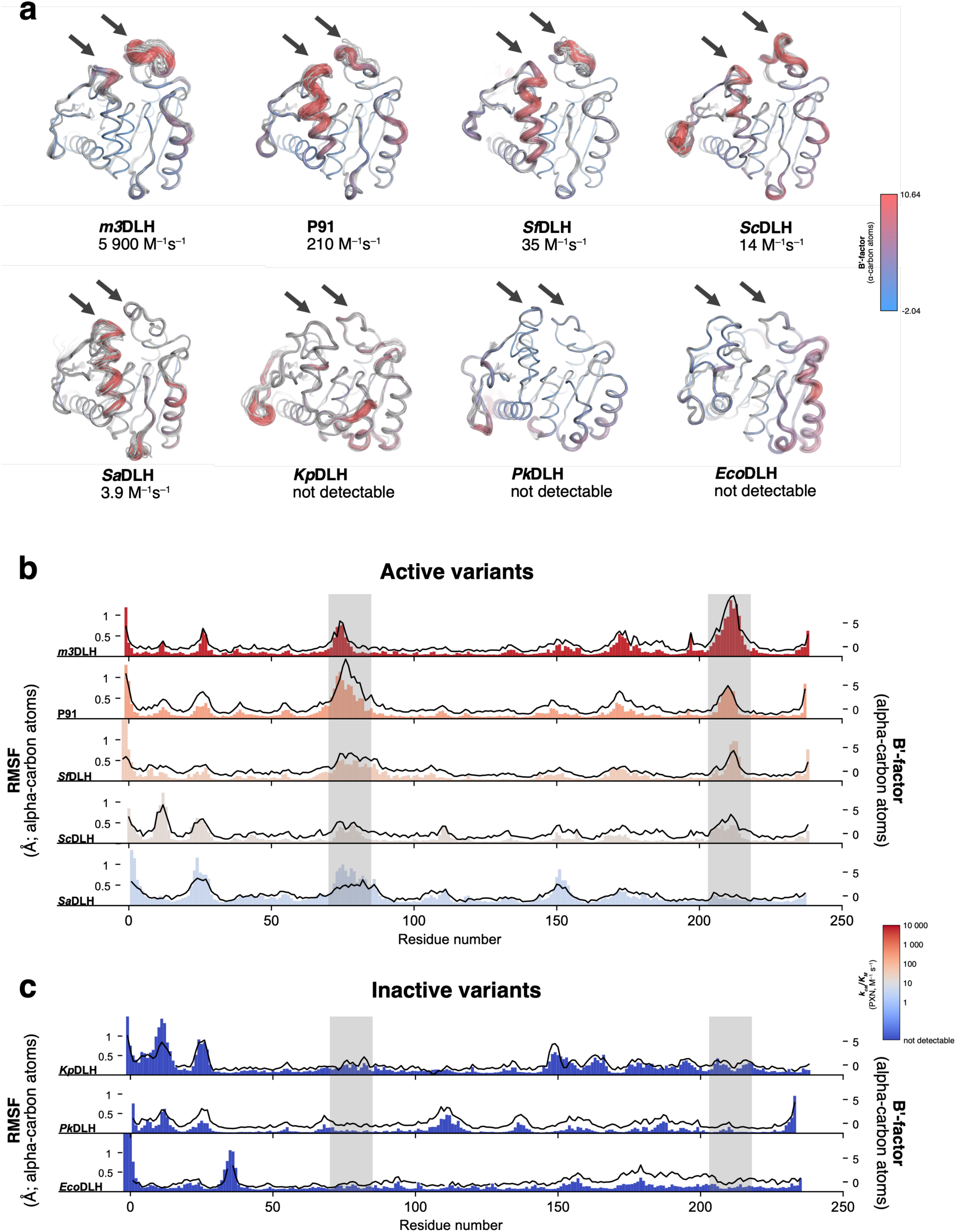
The flexibility of active-site-surrounding loops correlates with phosphotriesterase activity. **(a)** Ensemble refinement and B-factor analysis reveal that loops 5 and 14 have higher mobility in homologues with phosphotriesterase activity. For each enzyme, 20 poses are shown as ensemble in grey. Normalised B-factors (B’) are shown in putty representation in a colour gradient. Loop 5 and loop 14 are highlighted with black arrows. **(b–c)** Comparison of root mean square fluctuations (RMSF) of overlayed structural ensembles and B’-factors of active (b) and inactive (c) homologues. RMSFs are shown as coloured bars, B’-factors as black line. The structural features covering the active site, the C-terminal end of helix B and the adjacent stretch of loop 5, as well as loop 14, are highlighted in grey boxes. Structures used for analysis (PDB IDs): *m3*DLH, 7JKA; P91, 4ZI5; *Sf*DLH, 8VB3; *Sc*DLH, 7JIZ; *Sa*DLH, 9N2W; *Kp*DLH, 3F67; *Pk*DLH, 4U2B; *Eco*DLH, 4ZV9.

To explore whether flexibility predictions would recapitulate our experimental observations, we used AlphaFold 3 structural models to derive proxys of conformational diversity (root mean square fluctuation of 50 overlayed independent models as well as average pLDDT). Both metrics successfully identified the increased flexibility of active site-surrounding loops in enzymes with promiscuous phosphotriesterase activity, even though they failed to provide quantitative discrimination between different activity levels (**Figure S19**).

### Loop grafting increases phosphotriesterase activity

To experimentally resolve the role of the loops we designed a small library of 80 variants that includes the exchange of entire loop elements from a more active enzyme into a less active homologue. As a testing scaffold we chose the DLH family protein *Hs*DLH (also known as CMBL (15)). *Hs*DLH is the only human DLH-family protein, a cytosolic protein of unknown function that had previously been shown to be involved in the pre-activation of the drug olmesartan by (presumably promiscuous) ester hydrolysis (15). Notably, *Hs*DLH, which is even more distant to P91 in sequence space than the non-active homologues *Pk*DLH and *Eco*DLH, displays very weak activity towards the fluorogenic phosphotriester FDDEP (*k_cat_* ≈ 1.6 × 10^-4^ s^-1^*, k_cat_/K_M_* ≈ 4.8 M^-1^s^-1^) and therefore represents, together with the well-known metal-dependent paraoxonase PON1 (7), the second reported human phosphotriesterase.

To construct a small combinatorial library amenable to microtiter plate screening, we chose four loop segments to be grafted into *Hs*DLH (**Figure 4a** and **Table S8**; for details on the choice of loop elements see **Note 1.6 in the Supplementary Methods**). The grafted loop segments were sourced from the homologues with highest phosphotriesterase activity: *Hm*DLH, *m3*DLH, P91-WT, and two evolved P91 variants (from a previous directed evolution experiment (12)) and cloned into *Hs*DLH, generating a small combinatorial library with a theoretical diversity of 80 variants. We expressed the library in *E. coli* and screened lysates of 264 clones in microtiter plates (3.3-fold oversampling of the library diversity) for turnover of the phosphotriester FDDEP. We purified and kinetically characterised the most active variant *Hs*DLH-1E11, revealing a 7.8-fold increase in *k_cat_*/*K_M_*, mainly conferred by increase in *k_cat_* (**Figure 4b**). Interestingly, in this variant the entire loop 14 (Fragment F4, **Figure 4c**) is replaced by the corresponding loop from the highly active phosphotriesterase P91-R2 that was evolved for turnover of FDDEP (12). In addition, the acidic residue stabilising the catalytic nucleophile in the inwards-pointing position is exchanged from an aspartate to a glutamate (D50E). The identification of a loop grafted-*Hs*DLH with significantly increased activity confirms the hypothesis that phosphotriesterase activity is co-determined by architecture of the active-site surrounding loops.

**Figure 4:**
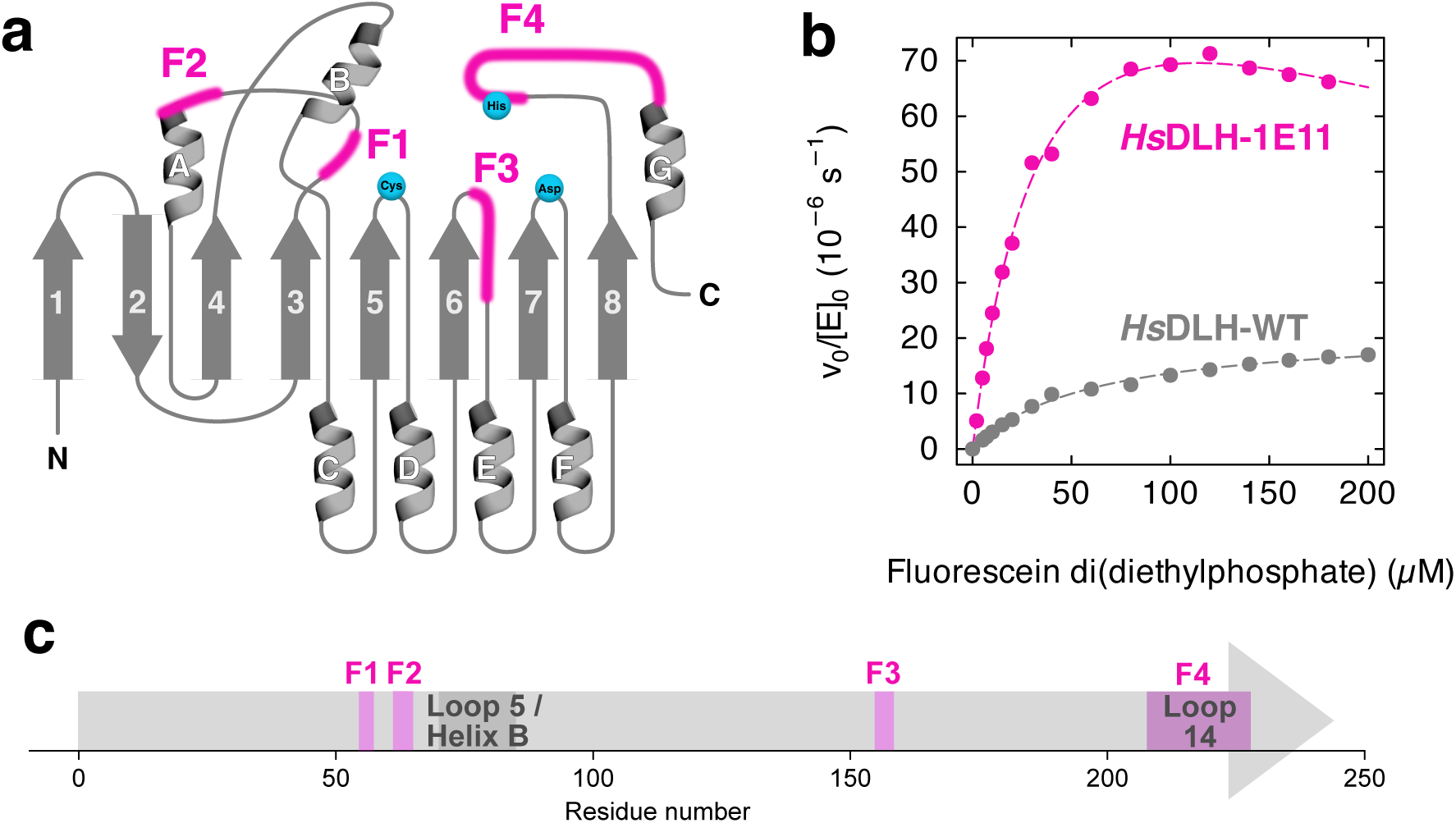
Loop grafting increases the phosphotriesterase activity of *Hs*DLH. **(a)** Schematic representation of the fragments (magenta) grafted into the small combinatorial *Hs*DLH library. Fragment 1 (**F1**) contains the acid residue stabilising the inwards-pointing conformation of the cysteine nucleophile, as well as a residue contributing to the oxyanion hole. Fragment 2 (**F2**) encompasses the transition between helix B and loop 5, which is in contact with loop 14 in both structural models of *Hs*DLH. Fragment 3 (**F3**) forms the bottom of the active site cavity where the attacking water or hydroxide molecule might be positioned. Fragment 4 (**F4**) encompasses the entire loop 14, which partly covers the active site cavity and whose flexibility correlates with phosphotriesterase activity across different P91 homologues. **(b)** Michaelis-Menten kinetics of wild-type and chimeric *Hs*DLH: The variant *Hs*DLH-1E11, which was isolated from the combinatorial library and contains fragment 1 (F1) from P91-WT and fragment 4 (F4, the entire loop 14) from P91-R2, has a ≈ 8-fold increased catalytic efficiency towards the phosphotriester substrate FDDEP as compared to the wild-type. Note that both wild-type and variant 1E11 carry an N-terminal His_6_-Twin-Strep-SUMO tag used to ensure soluble expression. This tag reduces catalytic efficiiency overall but was required for solubility of variant 1E11. **(c)** Schematic topology of fragments that were recombined in the library.

### Phosphotriesterase activities are truly promiscuous and not the result of specific evolutionary adaptation

The two closest homologues to P91, *m3*DLH and *Hm*DLH, even surpass the catalytic efficiency of P91 by one to two orders of magnitude, with *k_cat_/K_M_* of 5900 and 10000 M^1^s^-1^, respectively. These activity levels approach the median activity levels of naturally evolved, secondary metabolism enzymes (median *k_cat_/K_M_* of 7 × 10^4^ M^-1^s^-1^, median *k_cat_* of 2.5 s^-1^) (20). To address the question whether these relatively high activities result from natural selection rather than accidental promiscuity, we analysed the promiscuity patterns and their co-evolution in two series of experiments:

#### (i) Is transition-state adaptation reflected in promiscuity profiles?

First, we constructed promiscuity profiles for all enzymes with three substrates representing three different transition states: carboxyester (tetrahedron), lactone (distorted tetrahedron), and phosphotriester (trigonal-bipyramid). Evolutionary pressure induces an adaptation to a specific transition state geometry, as previously observed for an *in-vitro* evolved variant of P91 (12), so signs of specific adaptation to either the tetrahedral transition state of ester/lactone hydrolysis versus the trigonal bipyramidal transition state of phosphotriester hydrolysis would be expected. Specifically, a positive correlation between esterase and lactonase activities would reflect the similarity of their transition states and would stand in contrast to a negative trade-off between phosphotriesterase and esterase/lactonase that would indicate adaptation of the enzymes to the phosphotriesterase transition state.

For all enzymes, carboxyesterase activity was the highest and phosphotriesterase activity the lowest measured catalytic activity. Esterase and lactonase activities show a loose correlation (*R^2^* = 0.457, **Figure 5a**), indicating adaptation of these enzymes to the (similar) transition states of both reactions. However, no correlation or trade-off between phosphotriesterase activity and esterase or lactonase activity was discernible (*R^2^* = 0.004 and 0.006, respectively, **Figure 5a**). This indicates that the observed phosphotriesterase activity differences between close homologues are not governed by differences in adaptation to a specific transition state geometry but must be ascribed to other factors, such as leaving group preference or binding complementarity.

**Figure 5:**
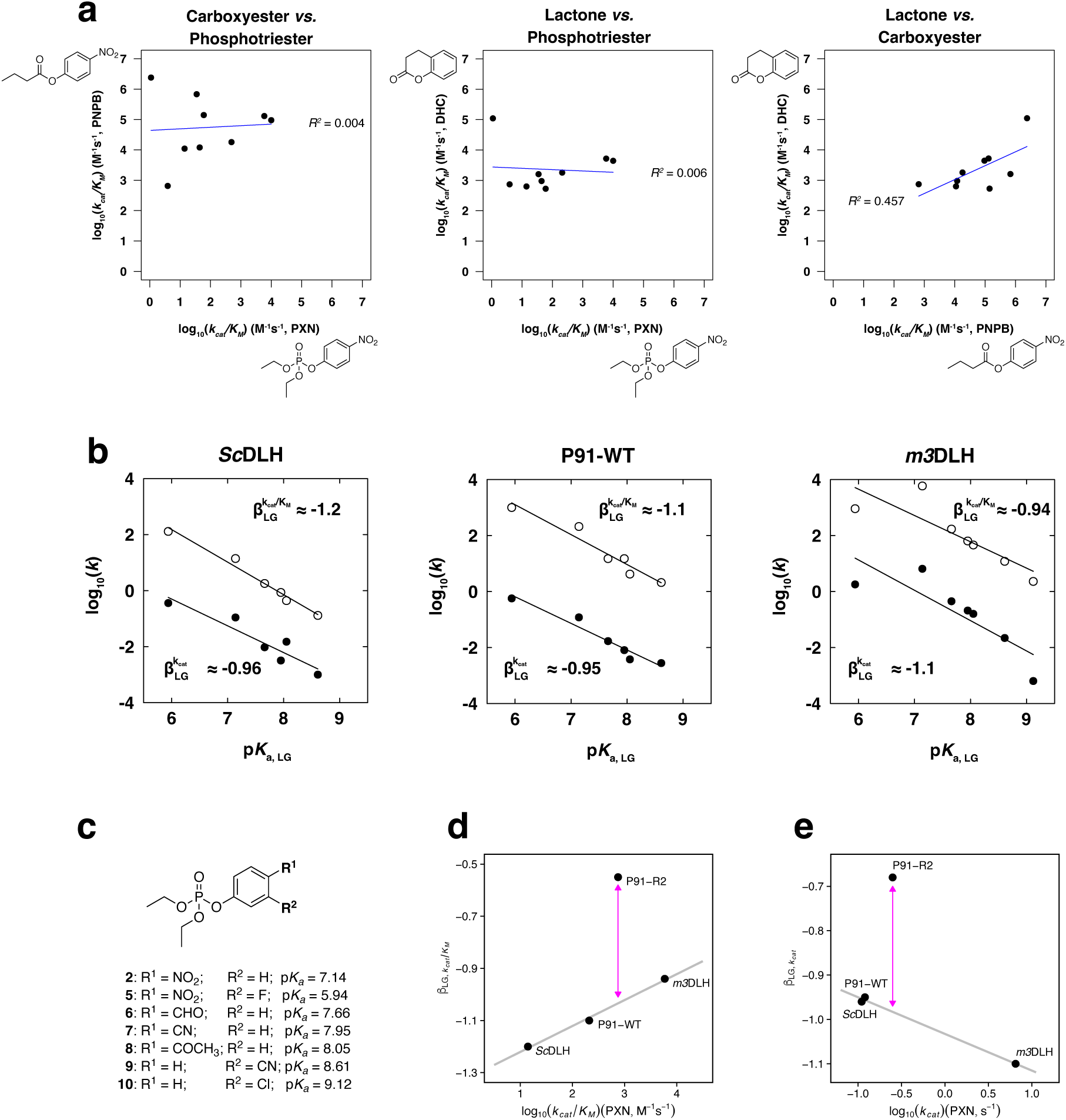
Promiscuity profiles and Brønsted analysis show that the phosphotriesterase activity is truly promiscuous. **(a)** Promiscuity profiles of the studied DLH family proteins displaying paraoxonase activity. Steady-state kinetics were measured for phosphotriester **2** (paraoxon-ethyl, PXN), carboxyester **4** (*p*-nitrophenyl butyrate, PNPB), and lactone **11** (dihydrocoumarin, DHC). The blue line indicates a linear fit. For all enzymes, carboxyesterase is the highest and phosphotriesterase the lowest activity. Phosphotriesterase activity follows no distinct correlation or trade-off with carboxylesterase or lactonase activity (*R^2^*= 0.004 and 0.006, respectively). In contrast, lactonase and carboxylesterase activities are correlated (*R^2^* = 0.457), indicating that the observed activity differences are not governed by differences in adaptation to a specific transition state geometry but might be due to other factors, such as leaving group preference or binding. **(b)** Brønsted analysis shows no differences in leaving group stabilisation for phosphotriesterase across non-evolved P91 homologues. The Brønsted plots show the linear free-energy relationship between the rate of hydrolysis of paraoxon derivatives and the p*K_a_* of the leaving group for *Sc*DLH, P91-WT, and *m3*DLH (in order of their phosphotriesterase activity), measured with phosphotriester substrates **2** and **5**–**10**. Filled dots: *k_cat_* in s^-1^. Open circles: *k_cat_*/*K_M_* in M^-1^s^-1^. (a) Brønsted plot for *Sc*DLH: *k_cat_*/*K_M_*: β_LG_ = –1.2, *R^2^* = 0.98; *k_cat_*: β_LG_ = –0.96, *R^2^* = 0.87. (b) Brønsted plot for P91-WT: *k_cat_/K_M_*: β_LG_ = –1.1, *R^2^* = 0.93; *k_cat_*: β_LG_ = –0.95, *R^2^* = 0.94. (c) Brønsted plot for *m3*DLH: *k_cat_*/*K_M_*: β_LG_ = –0.94, *R^2^* = 0.72; kcat: β_LG_ = –1.1, *R^2^* = 0.72. As the slope of the linear fits (β_LG_) is very similar for both kinetic parameters (*k_cat_* and *k_cat_*/*K_M_*) intermediate formation (*k_2_*) must be the rate-limiting step. All β_LG_ values are very similar between the different enzymes, despite the fact that they differ in phosphotriesterase activity by 400-fold. Hence, leaving group stabilisation does not contribute to the observed differences in phosphotriesterase activity. **(c)** Overview of substrates used to construct the linear free-energy relationships. **(d)** and **(e)**: β_LG_ values are almost constant across the three enzymes (β_LG_ ≈ -1), even though they differ in phosphotriesterase activity by two orders of magnitude. This finding stands in stark contrast to the ‘jump’ (purple arrows) in β_LG_ that P91 displayed upon directed evolution for a phosphotriester substrate (12). Although the difference in activity between *Sc*DLH and *m3*DLH is as large as between P91 and its evolved variant P91-R2 (≈ 400-fold), this is not reflected in their leaving-group stabilisation.

#### (ii) Transition-state adaptation analysed by free energy relationships

We further adressed the adaptation to a phosphotriester transition state by Brønsted analysis (21). To this end, we chose three enzymes with a wide range of paraoxonase activities: *Sc*DLH (low, *k_cat_*/*K_M_* = 14 M^-1^s^-1^), P91 (intermediate, *k_cat_*/*K_M_* = 210 M^-1^s^-1^), and *m3*DLH (high, *k_cat_*/*K_M_* = 5900 M^-1^s^-1^). We measured steady-state kinetics with a collection of paraoxon analogues with differing leaving group p*K_a_* values (substrates 2 and 5–10, **Figure 5c**) to establish a linear free-energy relationship for those enzymes (**Figure 5b**).

In the resulting Brønsted plots for all three enzymes, the slope of the linear fits (β_LG_) is very similar for both kinetic parameters (*k_cat_* and *k_cat_*/*K_M_*, **Figure 5b**), consistent with a scenario in which intermediate formation is the rate-limiting step, as previously shown for P91. However, all β_LG_ values are almost constant across the three enzymes (β_LG_ ≈ -1), even though they differ in phosphotriesterase activity by two orders of magnitude (**Figure 5d, e**). This finding stands in stark contrast to the marked increase in β_LG_ that P91 displayed upon directed evolution for a phosphotriester substrate (12). In other words: although the difference in activity between *Sc*DLH and *m3*DLH is as large as between P91 and its evolved variant P91-R2 (≈ 400-fold)), this is not reflected in their apparent leaving-group stabilisation. Hence, we conclude that differential leaving group stabilisation does not contribute to the observed differences in phosphotriesterase activity between *Sc*DLH, P91, and *m3*DLH.

The lack of efficient leaving group charge offset is consistent with the enzymes’ scattered promiscuity profiles and indicates that their phosphotriesterase activities are – independent of magnitude – not the result of adaptive evolution to a specific organophosphate transition state but instead fortuitous and accidental. Conversely, the observation of promiscuity (depending on the substrate with varying efficiency) indicates that these enzymes have a considerable, yet untapped potential for becoming efficient phosphotriesterases by adaptive evolution, just as demonstrated for P91 by active site-directed evolution of P91 (12).

## Discussion

The advent of large genomic and metagenomic sequence databases (e.g. MGnify with > 2 billion ORFs (2)) provides an untapped source of new enzymes and snapshots of their evolution. Making use of this treasure trove, however, requires many more functional assignments to link sequence information to one or more activities, comprising both dominant main as well as evolutionary relevant (22) promiscuous side activities. Currently sequence and even experimental structures are poor predictors of function across the large diversity in such databases, as homology-based assignments fail to be predictive over large sequence distances. AlphaFold-derived structure models also do not directly inform on function and are unable to reflect protein dynamics (e.g., brought about by loop movements (23)) that are intrinsic elements of mechanism), and cannot quantify the effects of transition state interactions with with individual amino acids. Experimental assignments of ‘pioneer’ activities in new sequence space are thus necessary to provide a ground truth that, with increaseing coverage of functional annotation, may ultimately make activity prediction possible and provide insight into hitherto unexplored catalytic strategies. For example, the discovery of such a ‘pioneer enzyme’, the metal-free phosphotriesterase P91, revealed a hydrolysis mechanism involving covalent catalysis (11) in the absence of metal cofactors and was found in a sequence neighbourhood where no phosphotriesterases had previously been known (11). Neither sequence similarity to existing phosphotriesterases nor phylogeny would thus have predicted phosphotriesterase function. We now expand the ‘pioneer’ bridgehead P91 across the wider dienelactone hydrolase (DLH) family, in which it lies. The 10 enzymes experimentally explored here (with homology decreasing to 27%) have near identical active sites (with a Cys-His-Asp catalytic triad and highly conserved active-site architectures), but differ by four orders of magnitude in catalytic rates. The focus on active-site residues alone is thus an insufficient marker of function in this family.

We establish a practical route to discovery of function by showing that expansion of the initial pioneer hit into a family provided a set of candidate enzymes, suitable for delineating catalytic features by comparison. A comparison of structure and activity suggested that latent catalytic potential in the active site triad is not simply governed by static active-site geometry, but by a portable dynamic feature: the flexibility of loops surrounding the active site providing conformational dynamics. The observation that catalytic efficiency correlates with loop length and dynamism provides a concrete experimental illustration of an emerging ‘sequence-dynamics-function’ view of enzymes. Conformational dynamics are increasingly recognised as an integral part of catalysis and fundamental to enzyme promiscuity and evolvability (24–27). Catalytic ensembles (rather than single enzymes) are increasingly used to define mechanism (28, 29). In this view, enzymes populate ensembles of states on a rugged energy landscape and catalysis proceeds through those ensembles rather than through a single static structure. This view is underpinned by recent studies showing that transitions in catalytic activity are accompanied by altered loop dynamics (30, 31). Enzyme families in which loops act as ligand-gated or rate-limiting features include TIM-barrel isomerases, tryptophan synthases, tyrosine phosphatases, and beta-lactamases (reviewed in ref. (27) and provide a basis for dynamic control of reactivity and binding. Further, loop grafting has been explicitly used to alter activity or specificity while leaving the core active site in place (32–38). Riziotis *et al*. (39) compared structures around conserved active site residues but found no systematic relationship between conformational variability and catalytic function. In contrast, our data show that flexible structural features away from the active site can become the decisive determinants of activity in DLH family enzymes. The ≈ 8-fold increase in catalytic efficiency obtained by transplantation of lid-loop regions from a highly active homologue into the weakly active human enzyme *Hs*DLH provides direct experimental validation that loop flexibility in this scaffold is not merely a correlative descriptor but a causal, transferable and, in principle, engineerable module for tuning phosphotriesterase function.

The DLH family possesses a mechanistic advantage for this function, rooted in its cysteine-containing triad. Based on our previous work (12), we propose that the cysteine nucleophile is pre-disposed to this activity, enabling rapid dephosphorylation and thereby shifting the rate-limiting step from chemical turnover to the initial formation of the covalent intermediate, i.e., to a step that depends on substrate access and orientation – the very step that the flexible loops modulate. It has been postulated that evolution of phosphotriesterase function in canonical serine hydrolases must navigate a binding versus turnover trade-off (40). Our evidence in the DLH-family is consistent with the idea that enzymes can bypass the constraints of this trade-off by mechanistic decoupling, promoting fast intermediate formation through flexible loops while also undergoing rapid dephosphorylation due to their cysteine nucleophile and the increased reactivity of the intermediate, making it an evolvable scaffold for multiple-turnover phosphotriester hydrolysis.

The broad observation of phosphotriesterase promiscuity raises a fundamental evolutionary question: do the high-activity marine homologues, such as *Hm*DLH, represent a “parallel universe” of latent promiscuity, or have we detected an activity that evolved in response to a (yet unidentified) source of anthropogenic organophosphates (10)? Our mechanistic data support the first interpretation: we observed no trade-off between phosphotriesterase activity and the native esterase or lactonase functions. Furthermore, a flat Brønsted relationship (β_LG_) across all homologues despite large rate differences argues against specific transition-state optimisation for phosphotriester hydrolysis. These are hallmarks of accidental, raw promiscuity, i.e., latent activity that does not reflect microbial adaptation to a naturally occurring substrate, or, alternatively anthropogenic organophosphates as a phosphate source, as recently suggested (10). However, these promiscuous enzymes are of course well positioned to respond to an environmental challenge and specialise for higher efficiency under selection.

At the same time, the ecological context of some of the studied promiscuous enzymes might indicate (weak) selection. A previous study (41) identified a promiscuous atrazine-degrading enzyme in the same organism, *Mangrovimicrobium sediminis* (42), from which we isolated *Hm*DLH. The coexistence, in a nutrient-limited mangrove sediment bacterium, of two distinct enzymes that can scavenge nitrogen (atrazine) and phosphorus (paraoxon-like triesters) from xenobiotic compounds potentially is consistent with a broader resource-acquisition strategy: low-level promiscuous activities can become selectively advantageous when nitrogen and phosphate are limiting, even if the selection pressure is insufficient to drive full catalytic optimisation.

Our findings have direct implications for enzyme engineering and discovery. The characterisation of *Hs*DLH as a second human phosphotriesterase (alongside PON1) (7, 43) establishes a new starting point for developing human-compatible organophosphate bioscavengers. As a human cytosolic enzyme, this scaffold avoids the immunogenicity of bacterial enzymes or the complex stability and expression issues of plasma-derived PON1. Crucially, our work provides the engineering blueprint: optimisation of *Hs*DLH via grafting the dynamic loop identified here presents a strategy based on a recognised, portable modular feature for developing a therapeutic to treat pesticide poisoning (44) or for protection against chemical warfare agents (45). More broadly, the functional transition across the DLH family from non-reactive to proficient phosphotriesterases hinges on this portable dynamic feature that had not been recognised before by sequence analysis or static structure comparisons. Our initial functional metagenomics screening thus revealed P91 as a pioneer enzyme that neither sequence-based annotation nor structural modelling (as long as, e.g., AI/ML approaches do not predict function) would have anticipated (11), as a suitable starting point for directed evolution (12). The present work shows that conformational – rather than purely chemical – innovation can drive the emergence of new enzymatic functions, and it identifies loop flexibility as the quantitative link to function, a relationship that computational prediction of structural variability (e.g., deriving per-residue RMSF values by treating arrays of AlphaFold model structures like conformers, akin to similar recent approaches (46)) may reproduce qualitatively, but not functionally assign or anticipate. By identifying and transferring this dynamic module, we demonstrate a practical example for conformationally guided enzyme engineering, consistent with the general view that conformational plasticity underlies both catalytic promiscuity and evolvability (24–27). Together, these results provide both analytical evidence (correlations across an enzyme family) and guidance for engineering (loop transplantation) that integrates dynamic features of proteins instead of static structures that guide most current enzyme engineering efforts. Taken together, this workflow comprises (i) ultrahigh-throughput library screening to cover wide diversity, (ii) detailed characterisation of a single hit and (iii) expansion into the periphery of the hit to draw up sequence-structure-activity relationship. It thus charts a way to discover novel function and mechanism in the vastness of sequence space that complements historical one-by-one characterisation of enzymes or purely computational design or exploration (47). Such ground truth data that sheds light on unexplored functions, e.g., as input for AI/ML approaches (48), is necessary especially for rare activities (including accidental, promiscuous ones), to enable better future functional exploration of the enormous amounts of functionally uncharacterised sequences currently resting as ‘biological dark matter’ (e.e. unclassified DNA of unknown function) in databases.

## Methods

### Mapping sequence-structure relationships

Sequence identity was determined from pairwise alignments using EMBOSS Needle (49) at default parameters (Matrix = BLOSUM62, GAP OPEN = 10, GAP EXTEND = 0.5, END GAP = false, END GAP OPEN = 10, END GAP EXTEND = 0.5). Where no experimental structure was available, structures were modelled using AlphaFold2/ColabFold (18, 19). Root mean square deviation (RMSD) of structural superpositions was determined using the ‘*super’* function in the PyMOL software (Version 2.5.2).

### Sequence similarity networks

All sequence similarity networks (SSNs) were generated using the EFI-EST web resource (https://efi.igb.illinois.edu/efi-est/) (50, 51). Edge calculation for the SSNs was done by all-by-all BLAST alignment of the input sequences using an *E*-value of 10^-5^ and excluding UniProt-defined fragments. For **Figure 1a,b** the number of nodes in the networks was reduced by using a fractioning parameter of N = 1000 (a, alpha/beta hydrolase superfamily) and N = 40 (b, DLH protein family). This setting selects every N^th^ sequence in the family, while keeping all sequences with Swiss-Prot annotations. Sequences were additionally trimmed to the domain boundaries containing the predefined input Pfam family. SSNs were visualised using Cytoscape 3.4.0 in the yFiles Organic Layout (52).

### Multiple sequence alignments and phylogenetic trees

Multiple sequence alignments were done with Clustal Omega at default settings (https://www.ebi.ac.uk/Tools/msa/clustalo/) (53). For the generation of the phylogenetic tree shown in **Figure 1c**, a multiple sequence alignment was generated using T-coffee Expresso (http://tcoffee.crg.cat/apps/tcoffee/do:expresso) (54), which takes structural information into account. This multiple sequence alignment was used as input for PhyML 3.0 (http://www.atgc-montpellier.fr/phyml/) (55). To choose a substitution model, automatic model selection with Smart Model Selection (SMS) was used with the AIC criterion (56). Tree search was initiated with BioNJ and tree improvement was done using nearest-neighbour interchanges (NNI). Bootstrapping was performed with N = 1000.

### Protein expression and purification

Plasmids isolated from single colonies were used to transform BL21(DE3) cells. Expression cultures were then inoculated by a similar ‘plating’ method as previously described (57, 58). In brief, a dense lawn of freshly transformed BL21(DE3) cells was directly scraped into 500 mL TB medium containing 100 μg/mL carbenicillin in a 2-L baffled erlenmeyer flask. The cells were grown for ≈ 60 min at 37 °C and 200 rpm shaking before being induced with anhydrotetracyclin (final concentration 200 ng/mL). Protein was then expressed at 20 °C/200 rpm for 18 to 20 h. Cells were harvested by centrifugation at 4 000 rcf for 10 min, the supernatant was discarded, and the dry pellet was stored at -80 °C.

Proteins were purified by affinity chromatography. All constructs were initially cloned with an N-terminal StrepII-tag (MASWSHPQFEKGA) which allows single-step affinity capture at very high purities on Strep-tactin resin. For high-yield purification, as required for transient-state kinetics, the StrepII-tag of *m3*DLH and *Sc*DLH was exchanged for an N-terminal His_6_-tag (`MHHHHHHGGS`) for purification by Ni-NTA affinity chromatography. Due to low soluble expression of *Hs*DLH, an N-terminal His-TwinStrep-SUMO-tag was cloned into the construct for higher solubility. *Hs*DLH with this tag was also purified by Ni-NTA affinity chromatography. The sequence of the His-TwinStrep-SUMO-tag is:

MGSSHHHHHHSSGLVPRGSHMASWSHPQFEKGGGSGGGSGGSAWSHPQFEKMSDSEVNQEAKPEVKPEVKPETHINLKVSDGSSEIFFKIKKTTPLRRLMEAFAKRQGKEMDSLRFLYDGIRIQADQTPEDLDMEDNDIIEAHREQIGGS

### Strep-Tactin affinity chromatography

For purification of StrepII-tagged proteins, the cell pellet was resuspended in lysis buffer (50 mM HEPES-NaOH, 150 mM NaCl, pH 8.0, 1 mM TCEP, 0.5–1 mg/mL lysozyme, 0.1 % Triton X-100, 0.01 % (= 25 units/mL) benzonase nuclease) and rolled for 30–60 min at room temperature (≈ 23 °C). Afterwards, the lysate was cleared by centrifugation at 20 000 rcf/4 °C for 20 min and the soluble fraction was loaded onto an equilibrated Strep-Tactin gravity flow column (Strep-Tactin Agarose Resin, IBA Life Sciences, Germany). The protein on the column was washed with 5 × 1 column volume of wash buffer (50 mM HEPES-NaOH, 150 mM NaCl, pH 8.0, 1 mM TCEP) and eluted with 6–8 × 0.5 column volumes of elution buffer (50 mM HEPES-NaOH, 150 mM NaCl, pH 8.0, 1 mM TCEP, 2.5 mM *d*-desthiobiotin).

### Ni-NTA affinity chromatography

For purification of His_6_-tagged proteins, the cell pellet was resuspended in lysis buffer (50 mM HEPES-NaOH, 150 mM NaCl, pH 8.0, 1 mM TCEP, 20 mM imidazole, 0.5–1 mg/mL lysozyme, 0.1 % Triton X-100, 0.01 % (= 25 units/mL) benzonase nuclease) and rolled for 30–60 min at room temperature (≈ 23 °C). Afterwards, the lysate was cleared by centrifugation at 20 000 rcf/4 °C for 20 min and the soluble fraction was loaded onto an equilibrated Ni-NTA gravity flow column (Super Ni-NTA Agarose Resin, Neo Biotech, France). The protein on the column was washed with 5 × 2.5 column volumes of wash buffer (50 mM HEPES-NaOH, 150 mM NaCl, pH 8.0, 1 mM TCEP, 20 mM imidazole) and eluted with 5 × 0.5 column volumes of elution buffer (50 mM HEPES-NaOH, 150 mM NaCl, pH 8.0, 1 mM TCEP, 250 mM imidazole).

### Buffer exchange and protein quantification

Following affinity chromatography, the eluate was concentrated with a spin concentrator (Vivaspin 10 000 kDa MWCO, Sartorius, Germany) and subsequently exchanged into eluent-free assay buffer (50 mM HEPES-NaOH, 150 mM NaCl, pH 8.0, 1 mM TCEP) using PD 10 desalting columns (Cytiva, USA). Enzyme purity was controlled by SDS-PAGE and concentrations were determined by measurement of absorption at 280 nm on a NanoDrop 2000 Spectrophotometer (Thermo Fisher Scientific, USA), using an extinction coefficient calculated with the ProtParam web tool (https://web.expasy.org/protparam) (59).

### Michaelis-Menten kinetics

We measured steady-state kinetics with purified StrepII-tagged protein, keeping enzyme concentrations at least 10–100-fold lower than the lowest substrate concentration. We chose substrate concentrations to span ≈ 10-fold below and above *K_M_*, as far as not limited by substrate solubility. We determined optimal starting enzyme concentrations E_0_ and substrate concentration ranges for each variant and substrate combination by empirical sampling. We pre-dissolved substrates in DMSO. As we found that the DMSO content influences the catalytic parameters (while being required as a co-solvent for substrate stocks), we kept the series of substrate concentrations in stocks of 200-fold the final concentration in order to ensure constant DMSO concentration (0.5 %) across all substrate concentrations. Upon measurement, we diluted aliquots of these substrate stocks 1:100 in assay buffer (50 mM HEPES-NaOH, 150 mM NaCl, pH 8.0, 1 mM TCEP), of which we subsequently mixed 100 µL with 100 µL of 2-fold concentrated enzyme solution (= 2×E_0_) in microtiter plate wells. We monitored the progress of the reaction by absorbance (for wavelengths and extinction coefficients for each substrate leaving group see **Table S3**) or fluorescence (at an excitation wavelength of 480 nm and an emission wavelength of 520 nm for fluorescein) in a spectrophotometric microplate reader (Tecan Infinite 200PRO, Tecan, Switzerland) at 25 °C. For absorbance measurements at wavelengths < 320 nm we used a 96-well plate made of quartz. We determined the absorbance maxima and extinction coefficients for the leaving group of each substrate by an absorbance wavelength scan followed by a calibration curve (see Table S4). We extracted the initial rates by linear fit of the first measurements (at < 10 % progress of the reaction) for each substrate concentration and normalized them with an extinction coefficient determined from a calibration curve. To determine the Michaelis−Menten parameters *k_cat_* and *K_M_*, we fitted the initial rates to the Michaelis−Menten equation:

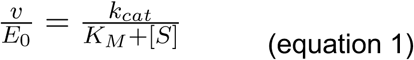

### Protein crystallisation and structure determination

Purified variants were concentrated to 10–20 mg/mL in 50 mM HEPES-NaOH, 150 mM NaCl, pH 8.0. Screens were optimised by sitting-drop vapour diffusion at 19 °C. Protein solutions were dispensed using a Mosquito Protein Crystallisation System (sptlabtech Ltd.; UK), with final drop volumes of 300 nL (1:2 ratio protein to crystallisation buffer). Crystals of *m3*DLH grew in a solution of 0.1 M sodium citrate (pH 5.6), 30 % (w/v) PEG 4K, and 0.2 M ammonium acetate. Crystals of *Sa*DLH grew in a solution of 0.1 M HEPES (pH 7.5) and 4.3 M NaCl. Crystals of *Sc*DLH grew in a solution of 0.1 M Tris (pH 7.0), 20 % (w/v) PEG 3K, and 0.2 M calcium acetate. Crystals of SfDLH grew in a solution 0.1 M Bicine (pH 9.0) and 65 % (v/v) 2-methyl-2,4-pentanediol (MPD).

All crystal samples were snap frozen in liquid nitrogen with no additional cryoprotection, and diffraction data were collected at Diamond Light Source (beamlines i03, i04, and i24) at 0.987 Å, 80 K. Diffraction data were indexed and integrated using XDS(60) and merged using AIMLESS as implemented within the CCP4 program suite(61). Data resolution cut-offs were chosen using half dataset correlation coefficients, as described previously(62). Structures were solved by molecular replacement (MOLREP)(63) using the structure of a putative dienelactone hydrolase from *Klebsiella pneumoniae* (*Kp*DLH, PDB ID: 3F67) as the input model. Refinement was performed using REFMAC(64) and phenix.refine(65), and manual rebuilding was done in Coot(66).

### Microtiter plate screening

To quantify the lysate activity of loop-grafted *Hs*DLH variants and for secondary screening of the combinatorial *Pk*DLH library, individual colonies were picked and grown in 96-deep-well plates in 500 μL Luria-Bertani (LB) medium with 100 μL/mL carbenicillin at 37 _°_C/1050 rpm for ≈ 14 h. Subsequently, 10 μL of overnight cultures were used to inoculate 490 μL of medium for expression cultures in 96-well deep-well plates which were grown at 37 °C/1050 rpm for ≈ 2 h until OD_600_ ≈ 0.5. Expression was then induced with anhydrotetracycline (final concentration 200 ng/mL; IBA Life Sciences, Germany) and carried out for 14 h at 20 °C and 1050 rpm shaking. Cells were pelleted by centrifugation at 3320 rcf for 60 min, the supernatant was then discarded, and cells were lysed with 100 μL lysis buffer (50 mM HEPES-NaOH, 150 mM NaCl, pH 8.0, 60 kU/mL rLysozyme, 1× BugBuster) for 20 min at 20 _°_C and 1050 rpm shaking. Cell lysates were diluted 1:20 or 1:400 in assay buffer (50 mM HEPES-NaOH, 150 mM NaCl, pH 8.0). For the reaction, 190 μL of the phosphotriester substrate FDDEP in assay buffer (200 μM) were added to 10 μL aliquots of the diluted lysate in microtiter plates and the formation of fluorescein was recorded in a plate reader (Infinite M200, Tecan, Switzerland) for 45 min at an excitation wavelength of 480 nm and an emission wavelength of 520 nm.

### Ensemble refinement

Ensemble refinements were carried out using the Phenix software suite (Version 1.21.2) (67). First, we removed anomalies from the reflections file using the inbuilt reflection file editor. Then, we added hydrogens to the structure file using the ‘ReadySet’ function. Finally, we used both files as input for the ‘phenix.ensemble_refinement’ function.

### B-factor normalisation

For comparison of B-factors across crystal structures, we normalised them using the BANΔIT toolkit (https://bandit.uni-mainz.de/) (68). We normalised the B factors of all atoms across all residues using the MAD_E_ method (IBM z-score) with a MAD threshold of 3.5.

### Prediction of conformational diversity and local confidence with AlphaFold

To assess the predicted conformational diversity and local confidence of the eight enzymes with available experimental structures, we generated 50 independent AlphaFold 3 (69) models for each enzyme. We then superimposed these models and calculated the root mean square fluctuation (RMSF) for all α-carbon atoms as a measure of predicted conformational diversity, analogous to the RMSF derived from ensemble refinements of experimental structures. Additionally, we calculated the mean pLDDT score from AlphaFold across all 50 structure models for each α-carbon. To align with the RMSF interpretation (where higher values indicate lower confidence/higher conformational diversity), the pLDDT scores were inverted (100−pLDDT) and plotted, in analogy to the normalised B-factors from the experimental structures.

### Associated content

Additional experimental procedures, figures, tables, and sequences are available in the Supplementary Information.

## Data availability

Crystal structures of *m3*DLH, *Sc*DLH, *Sf*DLH, and *Sa*DLH are deposited on the Protein Data Bank PDB (PDB IDs: 7JKA, 7JIZ, 8VB3, 9N2W).

## Acknowledgements

J.D.S. was supported by a Gates Cambridge Scholarship. We thank H. Adrian Bunzel for feedback on the manuscript. This work was supported by the Biotechnology and Biological Sciences Research Council (BB/W000504/1), the European Research Council (695669) and the Horizon Europe programmes *BlueRemediomics* (101082304) and *BlueTools* (10058118) (implemented by Innovate UK).

## Author contributions

**J. David Schnettler**: Funding acquisition, Conceptualization, Investigation, Writing - Original Draft, Writing - Review & Editing; **Eleanor Campbell**: Investigation, Data curation; **Rashid Khashiev**: Investigation; **Oskar James Klein**: Investigation; **Florian Hollfelder**: Funding acquisition, Conceptualization, Writing - Original Draft, Writing - Review & Editing

