## Supplementary Information for "A portable dynamic loop module enables phosphotriesterase function in cysteine-dependent hydrolases"

### 1. Supplementary Methods

#### 1.1 Materials

All chemicals were purchased from Sigma-Aldrich and all biological reagents from New England Biolabs, unless otherwise stated. Primers were purchased from Merck and genes for cloning were ordered from GeneArt/ThermoFisher Scientific. The plasmid pASK-IBA5+ with the StrepII-P91 insert was provided by Pierre-Yves Colin, who modified it from the commercial plasmid pASK-IBA5+ (IBA Life Science). The solubility tag His-TwinStrep-SUMO was excised from the commercial pSOL SUMO plasmid (Lucigen). All plasmid construct sequences are deposited in **Section 5** (Sequences) below.

#### 1.2 Substrates

An overview of all substrates is shown in **Figure S1**. The fluorogenic phosphotriester model substrate fluorescein di(diethylphosphate) (FDDEP, substrate **1**) was synthesised by Mark F. Mohamed as previously described (1). The paraoxon derivatives used for Brønsted analysis (substrates **5–10**) were synthesised by Oskar James Klein as previously described (2).

#### 1.3 Assignment of putative signature residues

In all four previously structurally characterised DLH family proteins (P91*, Pk*DLH, *Kp*DLH, and *Eco*DLH), the active-site cysteine of the triad is present in two conformations in the crystal structure, one pointing inwards and one pointing outwards (**Figure 2c**, **Figure S11b,f, g,** and **h**). Structural and computational evidence has been presented to demonstrate that in *Pk*DLH this dual conformation is part of a mechanism of substrate-induced activation, hypothesised to protect the oxidation-prone cysteine when not interacting with the substrate (3). Binding of the dienelactone substrate to residue R206 in the lid loop (loop 15) which partly covers the active site, breaks a chain of hydrogen bond bridges, resulting in the active-site cysteine C123 to swing from the inwards-pointing ('inactive') into the outwards-pointing ('active') conformation (4, 5). In P91 however, the residue corresponding to R206, D203, carries the opposite charge and is not part of a similar chain of hydrogen bonds reaching the catalytic cysteine C118. This renders substrate-induced activation, at least mediated by the same chain of interactions as in *Pk*DLH, unlikely in P91. The inwards-pointing conformation of C118 in P91 is stabilised by three neighbouring residues: E37, S200, and H141 (**Figure S11b**). Mutating E37 or H141 into an alanine does not fully abolish phosphotriesterase activity but makes P91 more susceptible to losing activity upon treatment with alkylating reagents, which proposes a role of these residues in protecting C118 from oxidation (6). The residues stabilising the inwards-pointing conformation of the catalytic cysteine are not completely conserved across the different studied homologues. Following the hypothesis that the subtle positioning of the two alternative conformations and the activation of the thiolate by these three residues could be a key determining feature of phosphotriesterase activity, these residues were systematically compared between all characterised homologues (**Figure S11**). In P91 and all other identified highly active enzymes, the alternative conformation of the cysteine nucleophile is stabilised by a Glu/Ser/His arrangement. However, enzymes displaying none or only marginal activities deviate from this consensus. Notably, *Sa*DLH is the only enzyme from the close sequence cluster around P91, which does not conform to this consensus, having an aspartate instead of a glutamate in the corresponding position. At the same time, the turnover rate of *Sa*DLH falls off by one order of magnitude as compared to the very similar, closely P91-related homologues (**Table S1**).

#### 1.4 A combinatorial library derived from the active-site patterns of homologues in the DLH family does not lead to improved triesterases

**Library design and construction**

Our objective was to extract and test active-site determinants from DLH family homologues to probe whether specific residue arrangements confer phosphotriesterase activity. As droplet microfluidics enables full oversampling of defined combinatorial libraries, both the presence and absence of active variants become informative, allowing hypotheses to be tested by protein engineering. We therefore designed a binary combinatorial library to examine whether grafting residues from active homologues into an inactive scaffold could generate phosphotriesterase activity. As the archetypical but inactive DLH family member with known structure and native function, we selected *Pk*DLH as the starting scaffold. Despite low sequence identity, *Pk*DLH and P91 are structurally highly similar (RMSD ≈ 1.5 Å), with most first- and second-shell residues positionally conserved. The main structural differences lie in helix B and loops 5 and 14 (**Figure S15**). We selected candidate positions by comparing the active-site environments of *Pk*DLH, P91, and its active homologues. We then inferred homologous residues in proteins lacking structural data from SWISS-MODEL structural predictions (7). We chose residues according to three criteria:

i. Proximity to the catalytic triad (within 12 Å of the active-site cysteine);
ii. Interaction with the extended hexad (residues contacting the catalytic triad or its auxiliary stabilising residues in P91); and
iii. Differential consensus (identical or chemically similar in active homologues but distinct in *Pk*DLH).

Applying these criteria yielded 20 candidate specificity-determining positions (**Figure S12**, **Tabe S7**). A variant carrying all 20 substitutions simultaneously proved insoluble even with a solubility-enhancing N-terminal SUMO tag, and no activity could be detected. A combinatorial approach was therefore required. Complete randomisation of 20 residues ($20^{20} \approx 10^{26}$ variants) exceeds feasible screening capacities ($10^{6}$–$10^{7}$ droplets). We thus restricted variation to a binary choice at each site (wild-type or consensus residue), resulting in a screenable diversity of $\approx 10^{6}$ variants.

To assemble this binary library in the plasmid pASK-IBA5+, we divided the *Pk*DLH gene into seven fragments generated by PCR using mutagenic primers (**Figure S13**). Fragments were combined by BsaI-mediated Golden Gate assembly. As one-pot ligation was inefficient we ligated the fragments in subsets, followed by limited-cycle PCR (8 cycles) for amplification of the full-length assembly and size selection on an agarose gel. The final ligation into the pASK-IBA5+ backbone yielded a library of > $3 \times 10^{7}$ transformants with < 0.5 % background. Sequencing of random clones confirmed > 80 % full-length assemblies without duplication or deletion.

##### Microfluidic droplet screening

We screened the library for activity towards the fluorogenic phosphotriester 1 (FDDEP) as previously described for P91 (2). We transformed plasmids into *E. coli* BL21(DE3) to maximise expression, then encapsulated the cells into picolitre droplets containing lysis buffer and substrate. We analysed and sorted the droplets by fluorescence at a throughput of ≈ 0.5–1 kHz.

Because *Pk*DLH displays no detectable baseline activity, we increased assay sensitivity by raising the substrate concentration from 3 to 50 µM FDDEP and extending the incubation time to three days. We screened approximately six million droplets and collected the top 2.5 % with the highest fluorescence. We repeated the enrichment twice (three total rounds), each time using the previously sorted fraction as input.

After droplet sorting, we picked and screened ≈ 500 clones in microtiter plates. In parallel, we analysed one 96-well plate each of the unsorted and enriched libraries to monitor changes in the distribution of activity.

##### Analysis of screening outcome

Secondary screening in microtiter plates revealed no substantial enrichment after three rounds (**Figure S14**). A modest increase in apparent activity (≈ 4-fold over *Pk*DLH WT lysate) disappeared when normalised against empty-vector controls, which consistently showed higher background hydrolysis. The elevated baseline in the empty vector likely stems from transcriptional read-through in pASK-IBA5+, enhancing expression of backbone genes (TEM-116 beta-lactamase and Tet repressor), one of which may exhibit weak promiscuous phosphotriesterase activity.

The slight enrichment observed nonetheless justified sequencing of the 15 most fluorescent clones. Nine clones contained frameshifts producing short peptides; their enrichment mirrors similar artefacts reported in functional metagenomic screens (8), where non-enzymatic peptides may act as transcriptional modulators of weak host background activities. The six remaining clones encoded full-length proteins bearing 8–11 mutations, but none exhibited measurable phosphotriesterase activity above chemical background. We thus concluded that no variant with detectable phosphotriesterase activity was present in the library, leading us to the conclusion that latent phosphotriesterase activity in DLH-family enzymes is not solely determined by a specific active-site residue arrangement.

**Alternative explanations for the absence of screening hits**

Given that the assembled DNA library (≈ 25-fold coverage) and screened droplet pool (≈ 6-fold oversampling) exceeded the theoretical diversity, it is unlikely that an active combination was missed.

A more plausible cause is scaffold incompatibility: some residue combinations may be non-tolerated in PkDLH without compensatory stability-restoring mutations. Stability constraints often limit evolvability (9–11).

Finally, the assay sensitivity may have been limiting. With a 2 pL droplet volume and a detection limit of ≈ 2.5 nM fluorescein (12, 13), ≈ 3000 turnovers per droplet are required for detection. Under comparable conditions, promiscuous phosphotriesterases with $k_{cat}/K_M \approx 50$ $M^{-1}$ $s^{-1}$ are detectable (13). As *Pk*DLH lacks any starting activity, the minimal activity increase needed to surpass this threshold remains uncertain.

#### 1.5 Calculation of library oversampling factors

Library oversampling factors were calculated according to the following equation(14):

$$F = 1 - e^{\frac{-L}{V}} \qquad \text{(equation 1)}$$

where *L* is number of samples, library size or screening effort; *V* is the total number of possible variants $X^n$ (where *X* denotes the number of codons and *n* the number of saturated residues); and *F* is the fractional library completeness, e.g., 0.95 for 95 %.

#### 1.6 Construction of the loop-grafting library

We serendipitously identified a P91 variant bearing an 18-amino-acid insertion in the helix adjacent to loop 5, flanking the active site. Unexpectedly, this large insertion did not disrupt folding or activity but instead doubled the paraoxonase activity (**Figure S18**). Encouraged by P91's tolerance to extensive loop modifications, we adopted an alternative strategy to the previous single-residue-focused 'active-site arrangement grafting', targeting entire loop or loop-adjacent fragments instead.

We selected four fragments (**Figure 4a**) based on sequence alignment and structural comparison between active homologues and the human DLH (*Hs*DLH):

(i) **Fragment F1**: includes the acidic residue that stabilises the inwards-pointing conformation of the cysteine nucleophile (D50 in *Hs*DLH; E37 in P91) and an adjacent residue contributing to the oxyanion hole (I51 in *Hs*DLH; A38 in P91).

(ii) **Fragment F2**: comprises the transition between helix B and loop 6, a region predicted to interact with loop 14 based on structural models of *Hs*DLH).

(iii) **Fragment F3**: contains a segment of loop 10 that forms the base of the active-site cavity, where the attacking water or hydroxide is positioned.

(iv) **Fragment F4**: spans the entire loop 14, which partly covers the active site and whose length and flexibility correlate with phosphotriesterase activity across P91 homologues (**Figure 3, Figure S16**).

We cloned all fragments, together with their corresponding wild-type *Hs*DLH sequences, into the *Hs*DLH scaffold by cassette mutagenesis and Golden Gate assembly. The resulting combinatorial library was expressed in *E. coli* with an N-terminal SUMO solubility tag, which was retained during expression and kinetic characterisation. To ensure comprehensive library sampling, we oversampled the theoretical diversity by a factor of 3.3, providing a ≈ 96 % probability that all sequence combinations were tested. We screened the library in microtitre plates at a substrate concentration of 200 μM FDDEP.

### 2. Supplementary Figures

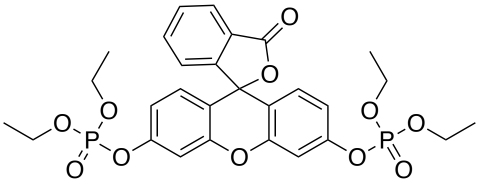

**1**

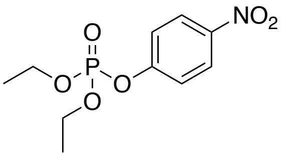

**2**

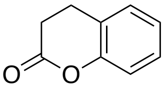

**3**

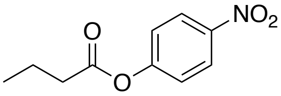

**4**

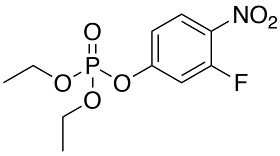

**5**

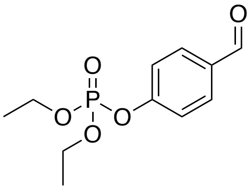

**6**

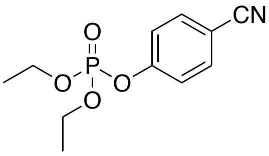

**7**

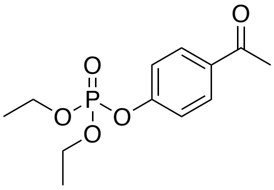

**8**

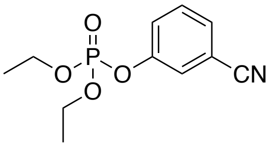

**9**

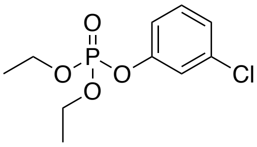

**10**

**Figure S1: Structures of the substrates used in this study. 1**: Fluorescein di(diethylphosphate) (FDDEP); **2:** Paraoxon-ethyl (PXN); **3:** Dihydrocoumarin; **4:** *p*-Nitrophenyl butyrate; **5**: 3-Fluoro-4-nitrophenyl diethylphosphate; **6**: 4-Formylphenyl diethylphosphate, **7**: 4-Cyanophenyl diethylphosphate, **8**: 4-Acetylphenyl diethylphosphate; **9**: 3-Cyanophenyl diethylphosphate; **10**: 3-Chlorophenyl diethylphosphate

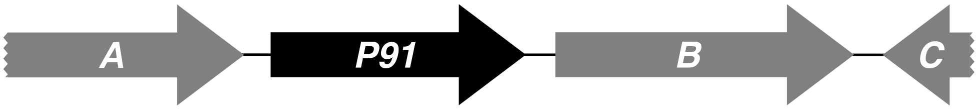

**Figure S2: Genomic context of P91.** The DNA fragment on which P91 was identified by a functional metagenomic screening(13) contains three further potential genes. Genes A and B can be annotated as Major Facilitator Superfamily (MFS) transporters, whereas C matches the C-terminal fragment of a NADP-specific glutamate dehydrogenase. The sequence similarity of gene fragment C was the basis for the identification of *Mz*DLH.

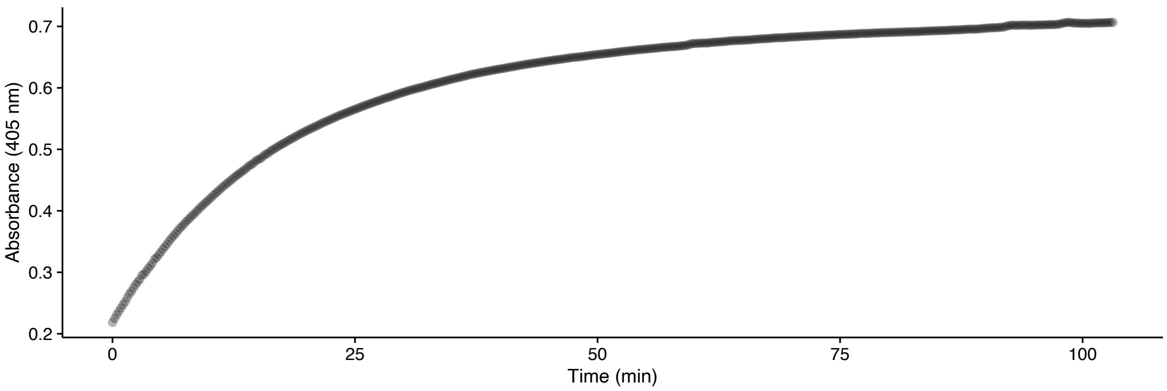

**Figure S3: Example progress curve of paraoxon-ethyl hydrolysis by m3DLH,** measured in 50 mM HEPES-NaOH, 150 mM NaCl, pH 8.0 at 25 °C at a substrate concentration of 1 mM and an enzyme concentration of 10 nM.

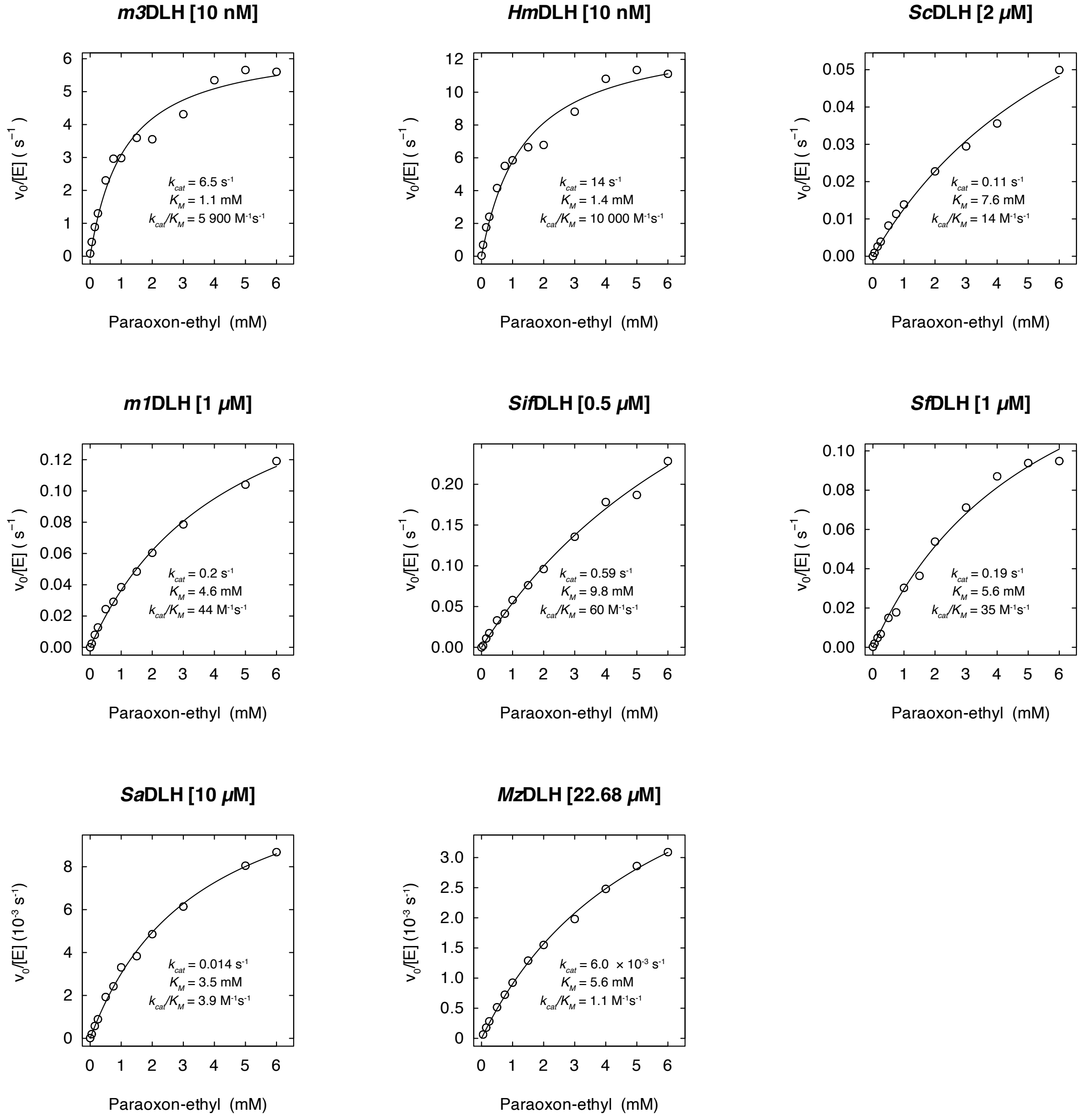

**Figure S4: Michaelis-Menten plots for steady-state kinetics of DLH family proteins with the phosphotriester paraoxon-ethyl (2)**, measured in 50 mM HEPES-NaOH, 150 mM NaCl, pH 8.0 at 25 °C.

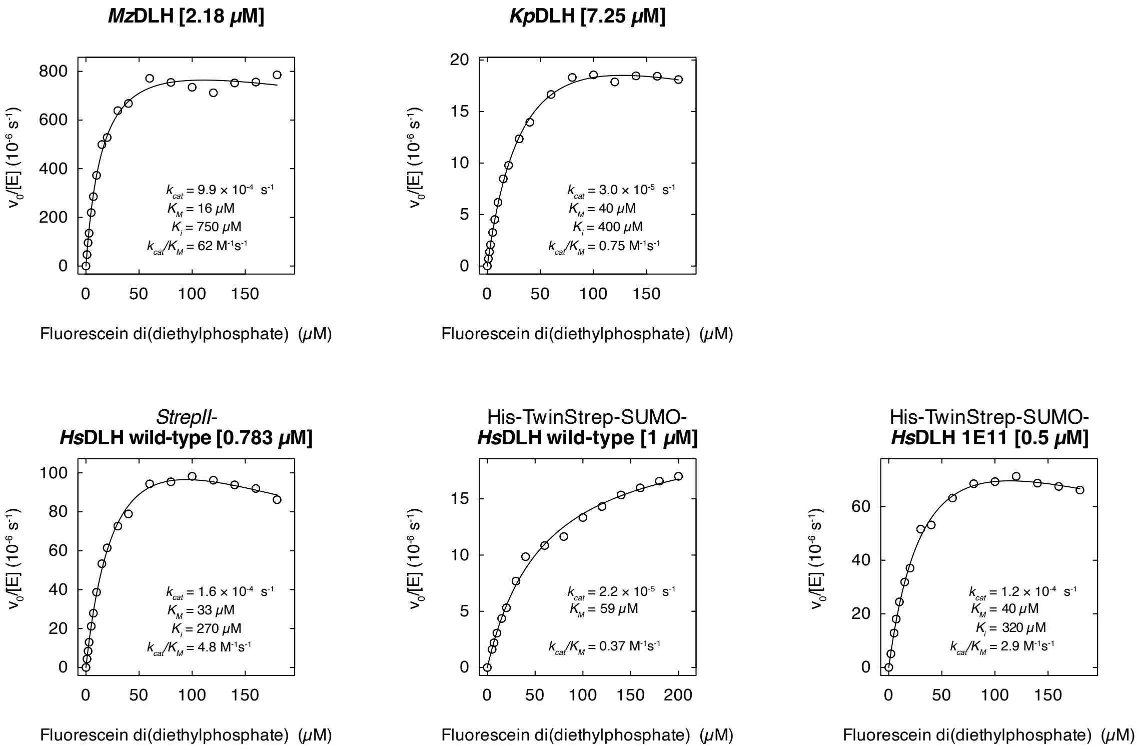

**Figure S5: Michaelis-Menten plots for steady-state kinetics of DLH family proteins with the phosphotriester fluorescein di(diethylphosphate)** (FDDEP, **1**), measured in 50 mM HEPES-NaOH, 150 mM NaCl, pH 8.0 at 25 °C.

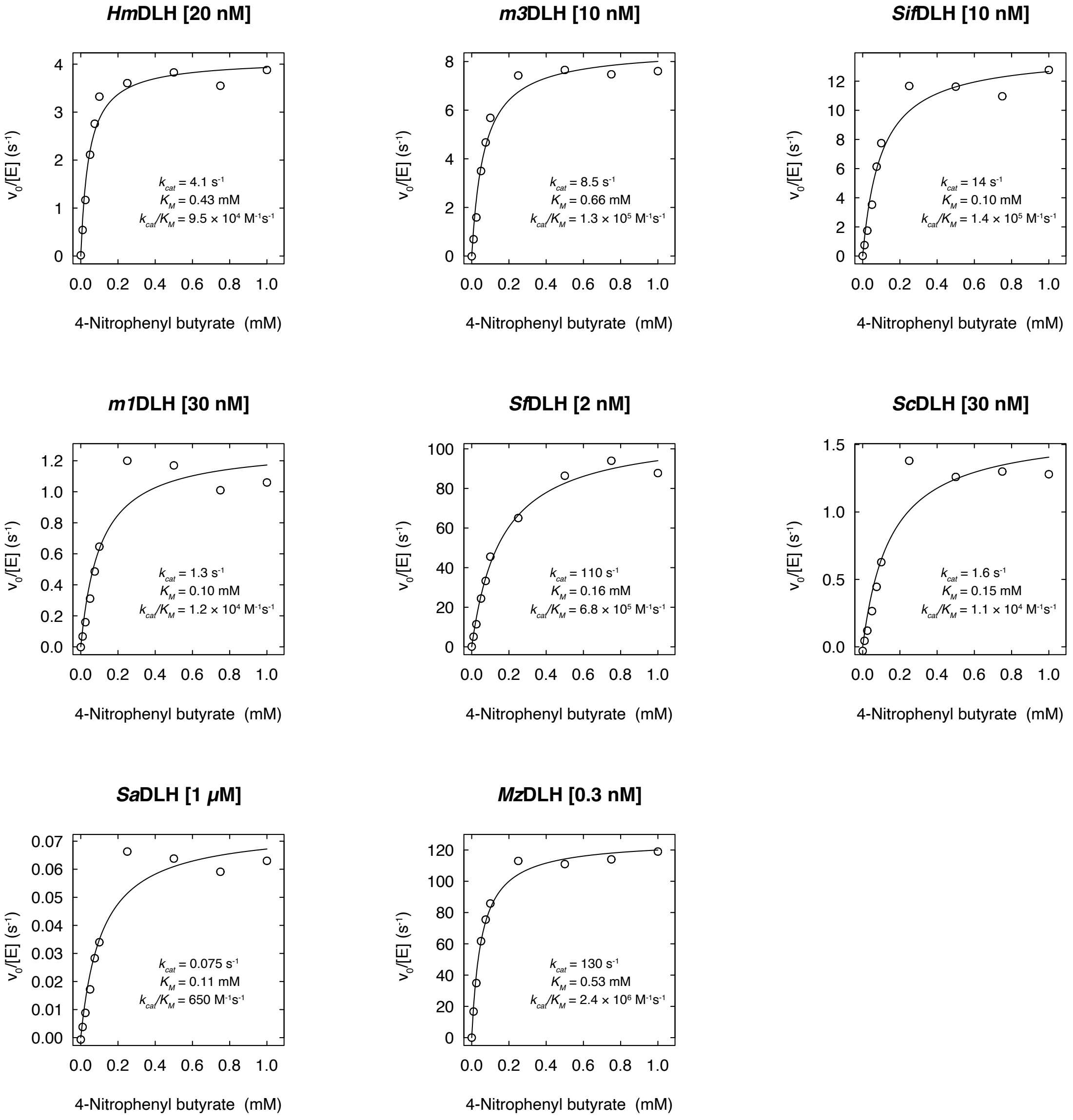

**Figure S6: Michaelis-Menten plots for steady-state kinetics of DLH family proteins with the carboxyester p-nitrophenyl butyrate (4)**, measured in 50 mM HEPES-NaOH, 150 mM NaCl, pH 8.0 at 25 °C.

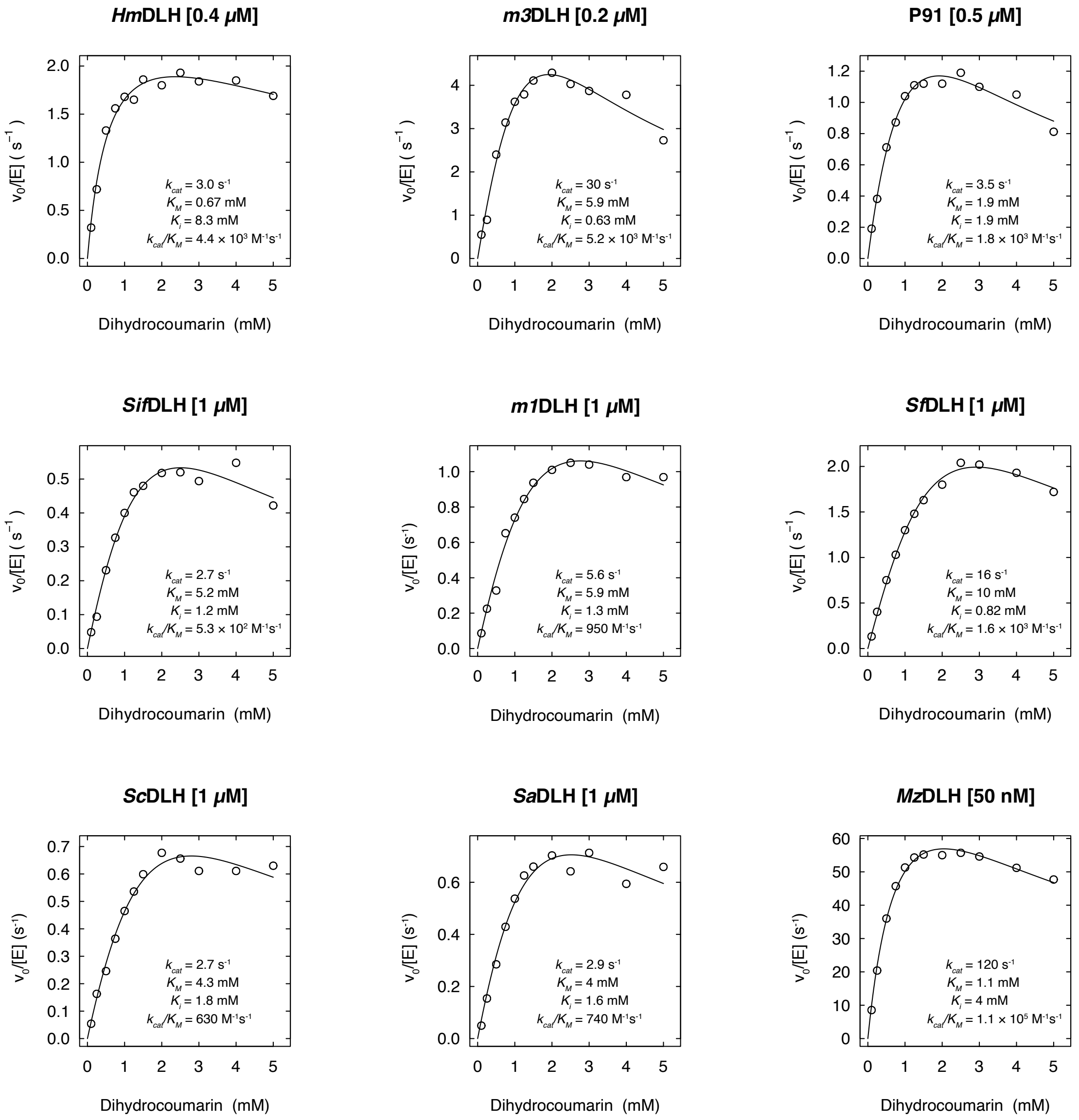

**Figure S7: Michaelis-Menten plots for steady-state kinetics of DLH family proteins with the lactone dihydrocoumarin (3)**, measured in 50 mM HEPES-NaOH, 150 mM NaCl, pH 8.0 at 25 °C.

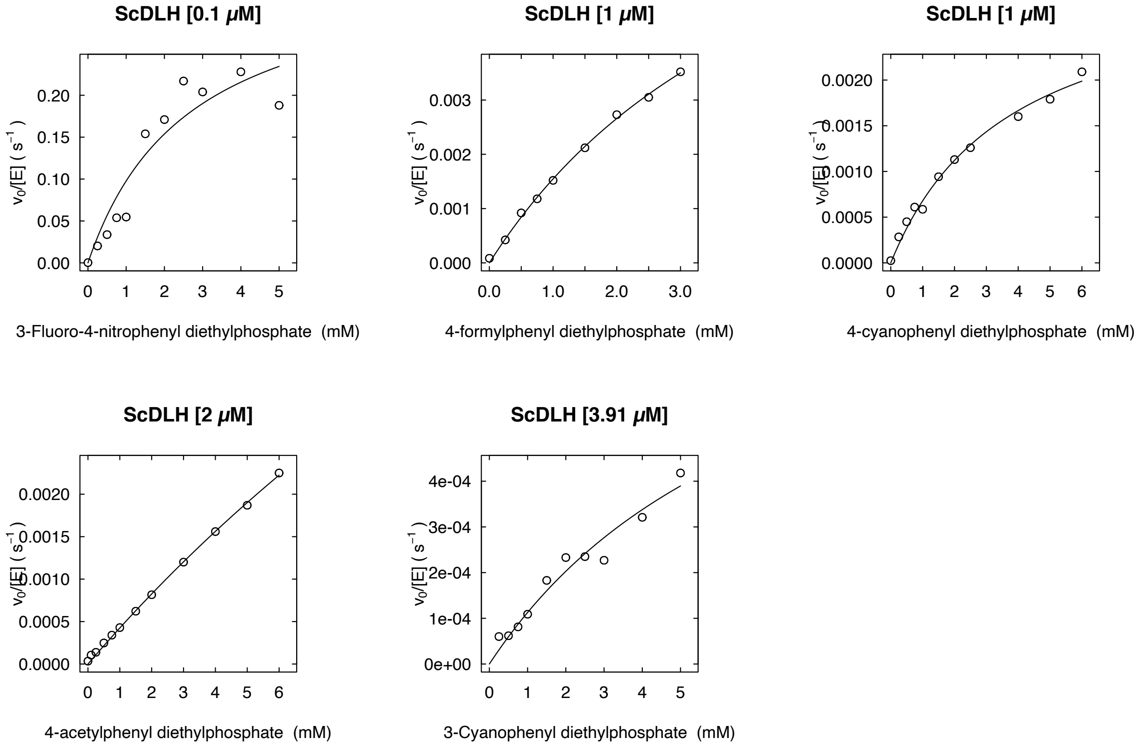

**Figure S8: Michaelis-Menten plots for steady-state kinetics of ScDLH with paraoxon derivatives 5–9 for Brønsted analysis**, measured in 50 mM HEPES-NaOH, 150 mM NaCl, pH 8.0 at 25 °C.

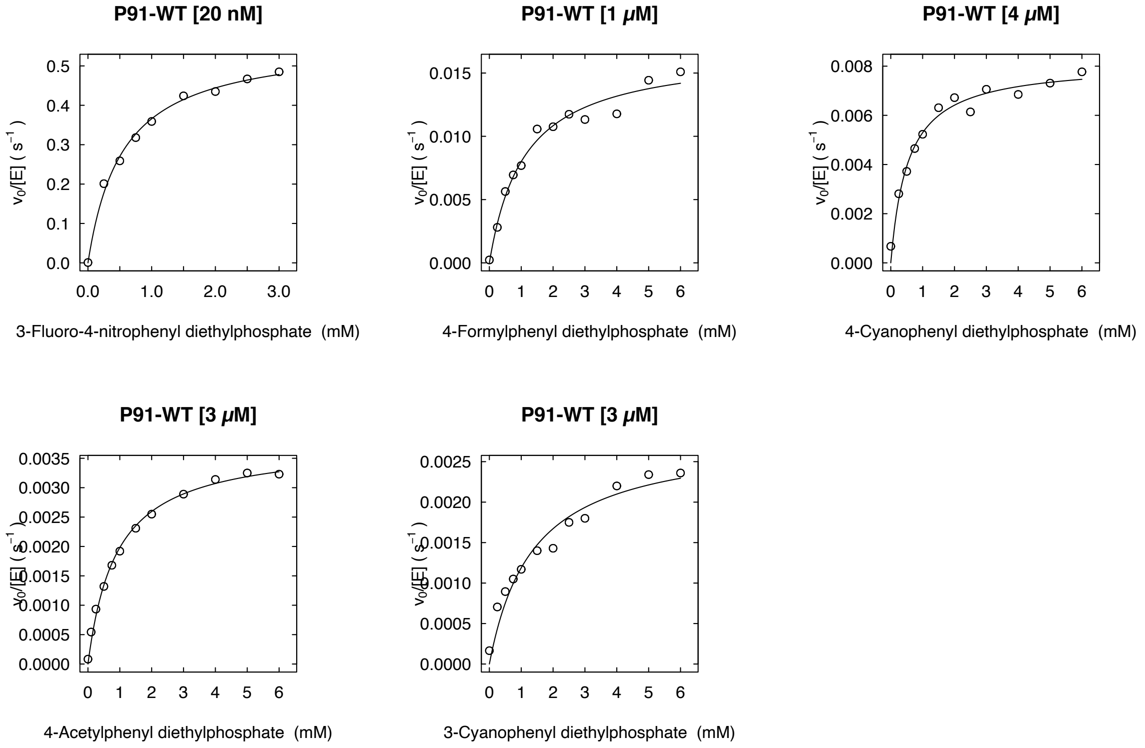

**Figure S9: Michaelis-Menten plots for steady-state kinetics of wild-type P91 with paraoxon derivatives 5–9 for Brønsted analysis**, measured in 50 mM HEPES-NaOH, 150 mM NaCl, pH 8.0 at 25 °C.

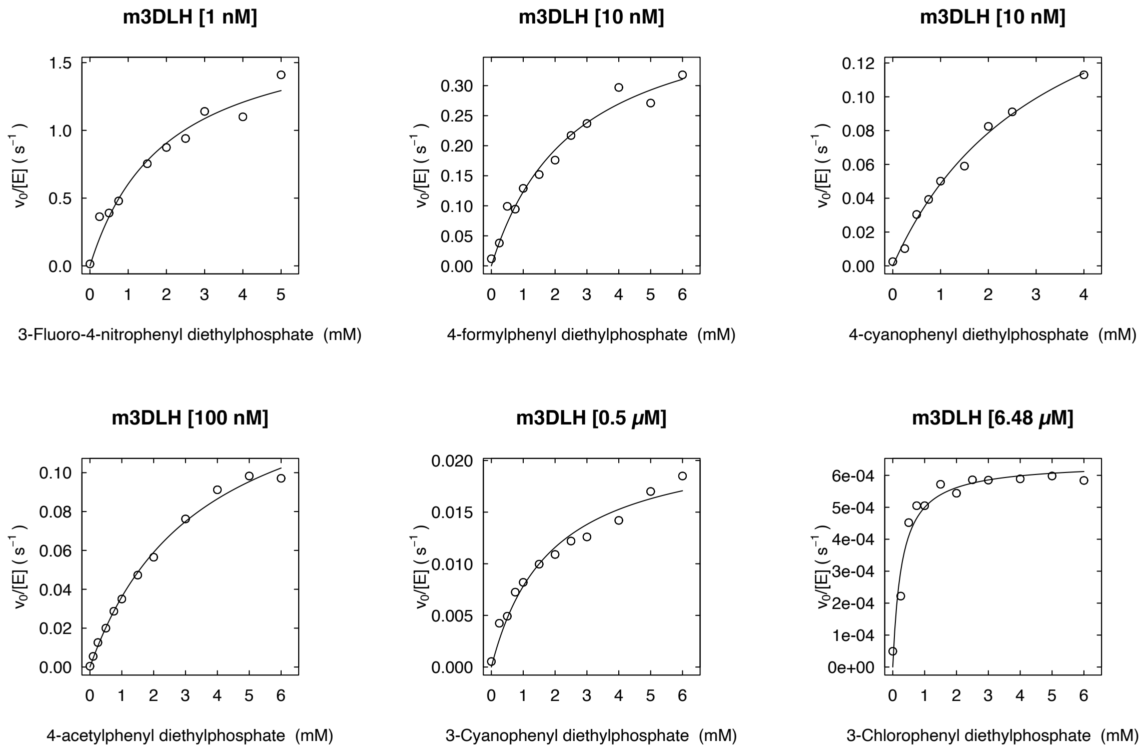

**Figure S10: Michaelis-Menten plots for steady-state kinetics of m3DLH with paraoxon derivatives 5–10 for Brønsted analysis**, measured in 50 mM HEPES-NaOH, 150 mM NaCl, pH 8.0 at 25 °C.

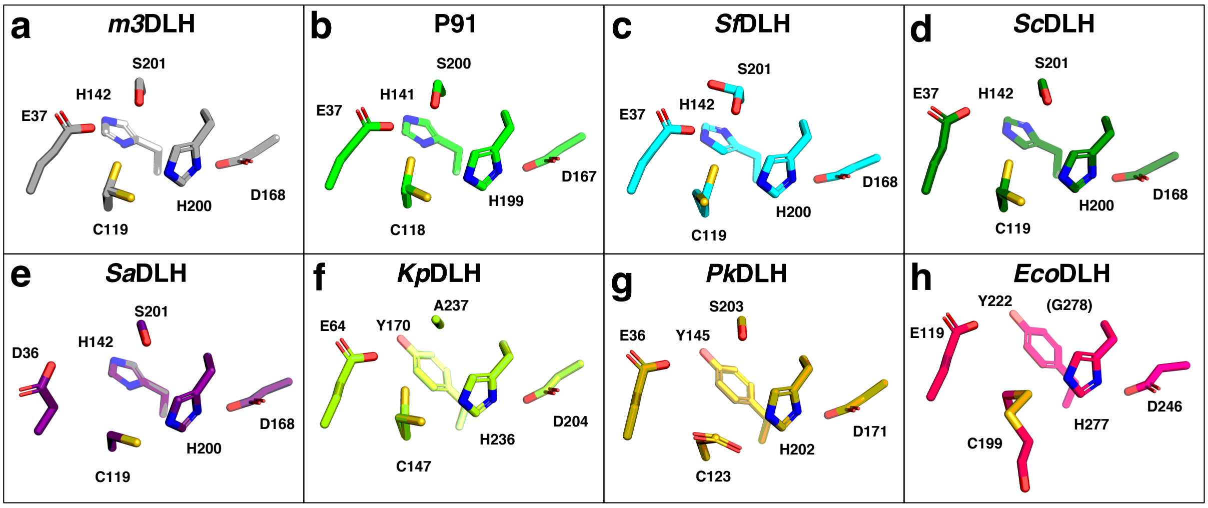

**i**

| | $k_{cat}/K_M$ ($M^{-1}s^{-1}$, PXN) | E37 | C118 | H141 | D167 | H199 | S200 |
|---|---|---|---|---|---|---|---|
| ***Hm*DLH** | ***10 000*** | **E** | **C** | **H** | **D** | **H** | **S** |
| ***m3*DLH** | ***5 900*** | **E** | **C** | **H** | **D** | **H** | **S** |
| **P91** | **210** | **E** | **C** | **H** | **D** | **H** | **S** |
| ***Sif*DLH** | ***60*** | **E** | **C** | **H** | **D** | **H** | **S** |
| ***m1*DLH** | ***44*** | **E** | **C** | **H** | **D** | **H** | **S** |
| ***Sf*DLH** | ***35*** | **E** | **C** | **H** | **D** | **H** | **S** |
| ***Sc*DLH** | ***14*** | **E** | **C** | **H** | **D** | **H** | **S** |
| ***Sa*DLH** | ***3.9*** | **D** | **C** | **H** | **D** | **H** | **S** |
| ***Mz*DLH** | ***1.1*** | **E** | **C** | **H** | **D** | **H** | **S** |
| ***Kp*DLH** | ***n.d.**** | **E** | **C** | **Y** | **D** | **H** | **A** |
| ***Hs*DLH** | ***n.d.**** | **D** | **C** | **Y** | **D** | **H** | **G** |
| ***Pk*DLH** | ***n.d.*** | **E** | **C** | **Y** | **D** | **H** | **S** |
| ***Eco*DLH** | ***n.d.*** | **E** | **C** | **Y** | **D** | **H** | **G** |

**Figure S11: Structural alignment of the active sites of all characterised DLH family proteins**, arranged in decreasing phosphotriesterase activity from top left **(a)** to bottom right **(h)**. All structural arrangements are shown in identical orientations, based on structural superposition of the publicly available (P91, *Kp*DLH, *Pk*DLH, *Eco*DLH) and the newly determined structures of DLH family proteins (*m3*DLH, *Sf*DLH, *Sc*DLH, *Sa*DLH). The tree residues of the Cys–His–Asp triad are shown as well as the three residues which are presumably stabilising the alternative conformation of the triad cysteine, together referred to as the 'extended hexad'. Note that in **(e)**, **(g)**, and **(h)** the two alternative cysteine conformations are not reflected in the structure but are visible in the original electron density. The cysteine C123 in *Pk*DLH is oxidised and the apparent covalent mercaptoethanol adduct of C199 in *Eco*DLH is a false interpretation of the electron density which presumably represents two alternative cysteine conformations. The table in panel **(i)** shows the paraoxonase activities and the residues of the 'extended hexad' for all characterised DLH family proteins, including enzymes for which there is no structure available. Enzymes with promiscuous phosphotriesterase activity are highlighted in green, while enzymes with marginal activity on only one of the two tested phosphotriester substrates are highlighted in faint green. Residues mismatching the corresponding residues in P91 are highlighted in red. *n.d.*, no detectable phosphotriesterase activity; *n.d.**, no detectable activity on paraoxon, but marginal activity on FDDEP.

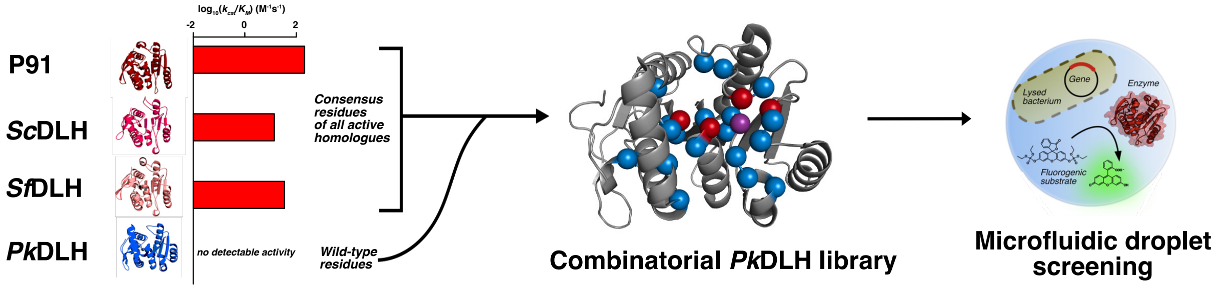

**Figure S12: Rationale and library design for the functional conversion of PkDLH into a phosphotriesterase**. Taking advantage of the high structural similarity despite low sequence similarity, *Pk*DLH and homologues with high promiscuous phosphotriesterase activity were structurally superimposed (only P91, *Sc*DLH, and *Sf*DLH shown) and all residues in the active site that are interacting with the catalytic triad or other key residues (red spheres in the structure) were matched across structures. Positions for which there was a consensus across all active homologues, while being different in *Pk*DLH, were then included into a combinatorial *Pk*DLH library. In this library, the 20 selected residues (blue spheres in the structure) can either have the wild-type side chain or the consensus side chain from the active homologues, resulting in a theoretical diversity of ≈ 1 × $10^6$ variants. This large 'binary' library was then screened for phosphotriesterase activity in microfluidic droplets.

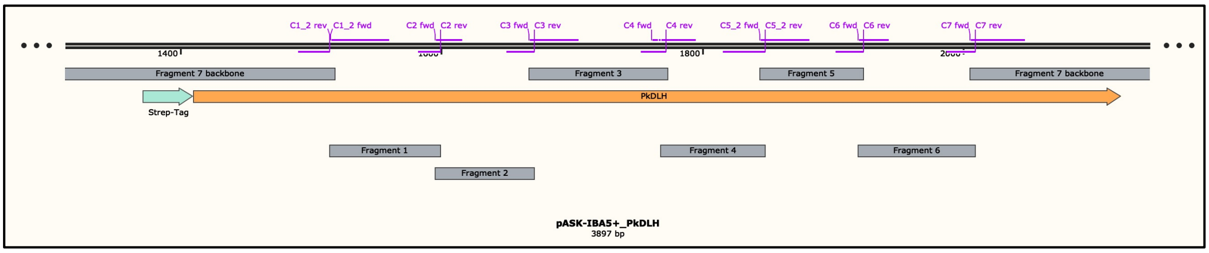

**Figure S13: Library cloning strategy for the assembly of the binary PkDLH library**. The *Pk*DLH gene (orange) was divided into 7 fragments (grey) which were created by PCR, thus introducing the mutations at the primer annealing sites (purple). The short fragments were then assembled separately and, after limited-cycle PCR amplification, ligated into the large backbone fragment.

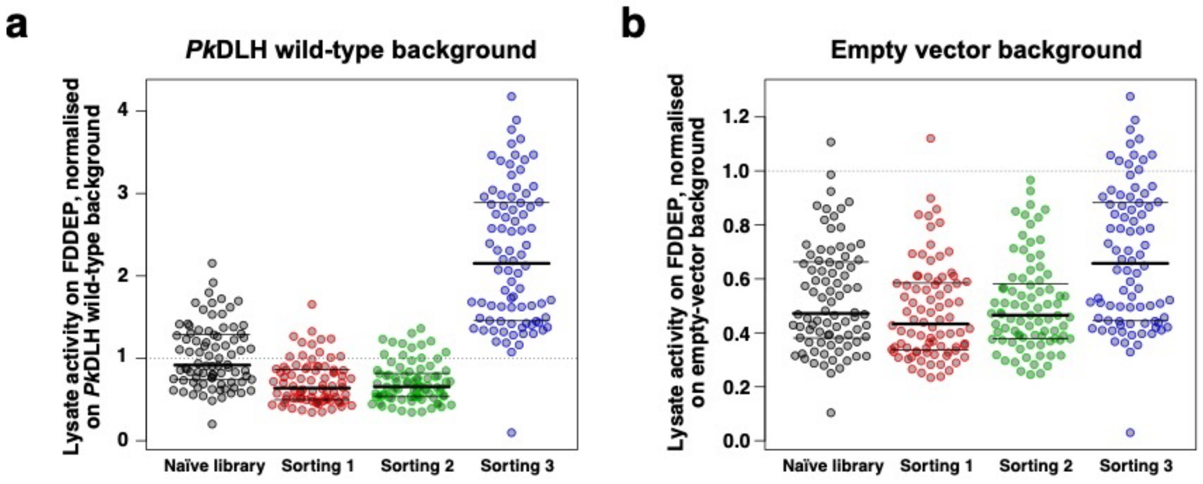

**Figure S14: Enrichment analysis of the binary *Pk*DLH library across droplet screening rounds reveals no strong enrichment for clones with increased phosphotriesterase activity**. To quantify the enrichment of active clones through droplet screening, 84 clones were randomly selected, before screening and after each droplet screening round, and their activity towards the phosphotriester substrate **1** (FDDEP) was measured in a microtiter plate-based lysate assay. The black horizontal lines in each beeswarm plot indicate the first quartile, the median, and the third quartile of the data. (a) Normalisation on lysate of bacteria expressing the wild type *Pk*DLH enzyme as a negative control. (b) The same data, normalised on lysate of bacteria with the empty vector as a negative control.

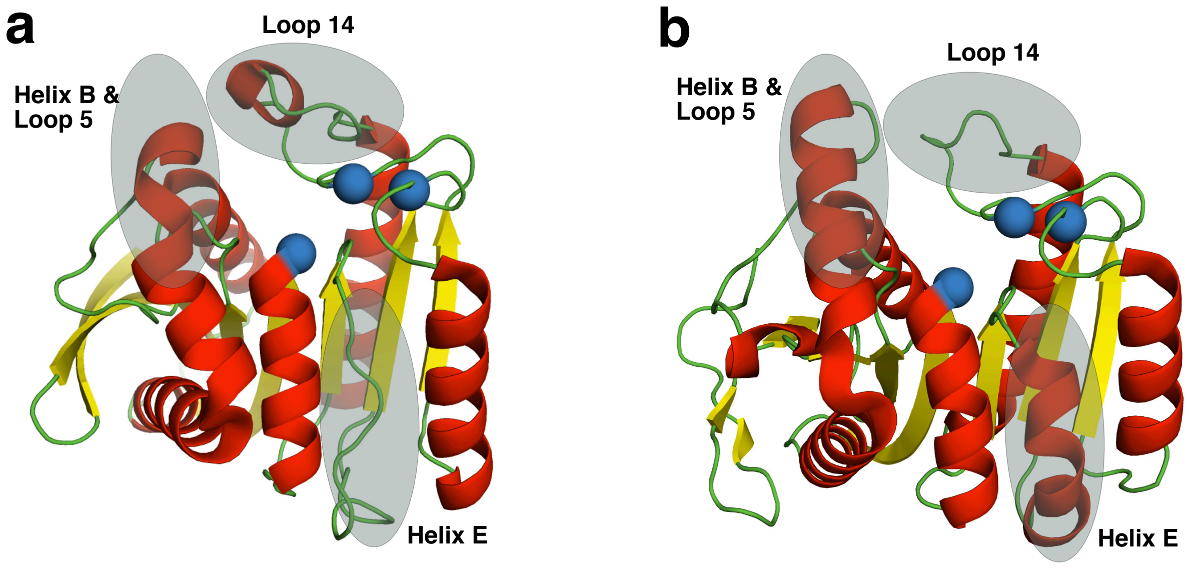

**Figure S15: Structural comparison of P91 and PkDLH reveals systematic differences**. **(a)** Schematic structure of P91. In P91, helix B is short, bends over the active site and disorders early into loop 5. In contrast, loop 14 is long and has a short helical element. The canonical helix E is missing and present as a disordered loop. **(b)** Schematic structure of *Pk*DLH. In *Pk*DLH, helix B is long and straight, whereas loop 14 is short and the canonical helix E is present. As loop 14 and helix B/loop 5 form the 'lid' domain above the active site, these differences result in a deeper, more closed active site cavity in P91 and a shallower, more open cavity in *Pk*DLH.

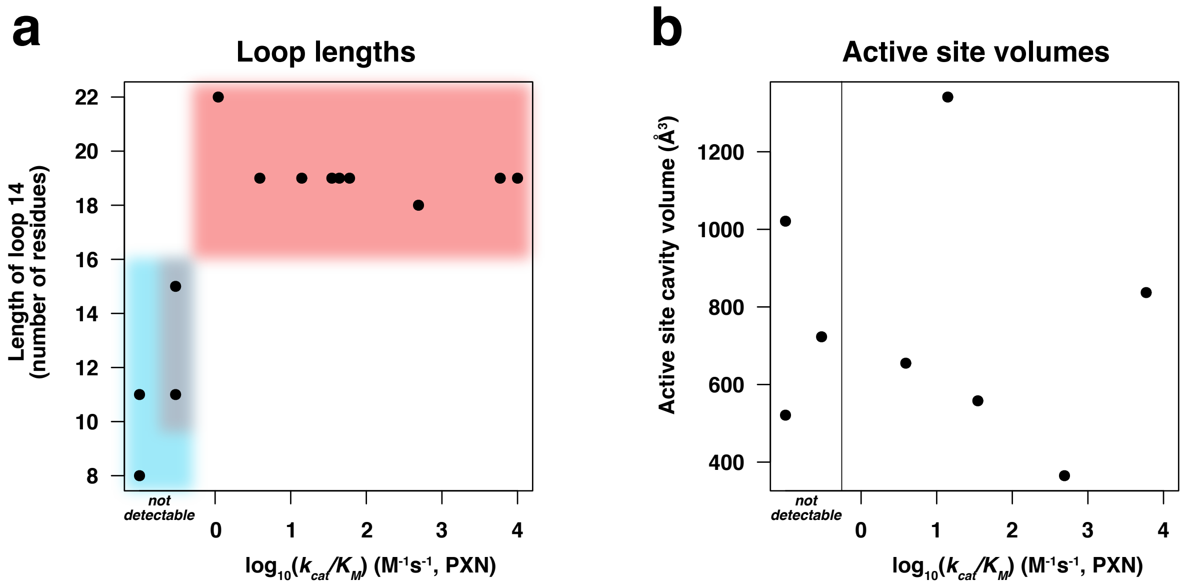

**Figure S16: Correlation of loop lengths and active site volumes with phosphotriesterase activity. (a)** The length of loop 14 correlates with phosphotriesterase activity.The number of residues in loop 14, as measured from the catalytic triad histidine to helix G, was plotted against the phosphotriesterase activity ($\log_{10}(k_{cat}/K_M)$) towards paraoxon, PXN) for every P91 homologue. While all active homologues have long loops of 18 residues or more (red sector), the inactive homologues have shorter loops (blue sector). The variants displaying marginal activity on the ore reactive phosphotriester FDDEP (*Hs*DLH, *Kp*DLH) lie in-between in loop length (grey sector). **(b)** Active site volumes do not correlate with phosphotriesterase activity. Active site volumes were measured for all P91 homologues of which as structure is available and plotted against their paraoxonase activity ($\log_{10}(k_{cat}/K_M)$).

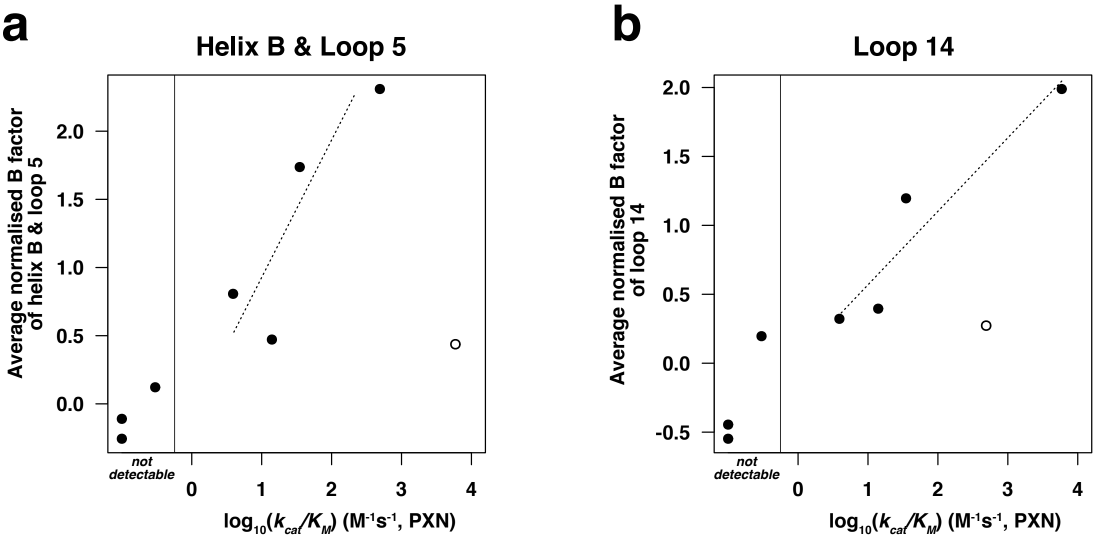

**Figure S17: Average B-factors of cavity-forming loop regions correlate with phosphotriesterase activity**. Normalised B-factors (B') of α-carbon atoms in all DLH family proteins with an available crystal structure were averaged over the two regions that partly cover the active site cavity. Two outliers (*m3*DLH in (a) and P91 in (b)) are highlighted in empty circles. Note that the value for the marginally active variant *Kp*DLH (no activity on paraoxon but low activity on FDDEP) was placed right of the non-active values in the 'not detectable' box to indicate its tendency in activity despite a measurable value for paraoxonase activity. **(a)** Average normalised B-factors of the left cavity-forming domain, formed by the transition of helix B and loop 5, correlate well with phosphotriesterase activity ($R^2$ = 0.76). **(b)** Average normalised B-factors of the left cavity-forming domain, formed by loop 14, also correlate well with phosphotriesterase activity ($R^2$ = 0.90). Both correlations indicate that the flexibility of the 'lid domain' formed by those two loops is a determinant of promiscuous phosphotriesterase activity.

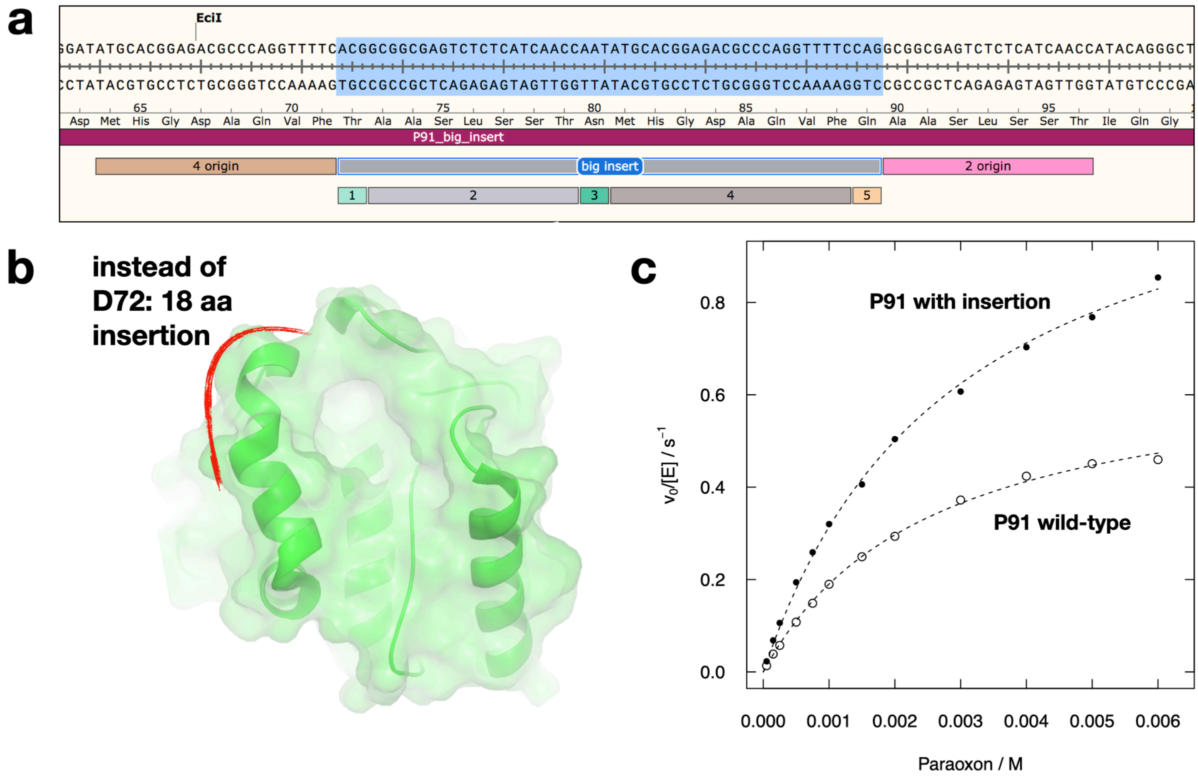

**Figure S18: A P91 variant with a 18-residue loop insertion in the cavity-forming domain is stable and displays increased phosphotriesterase activity** compared to the wild-type enzyme. (a) The insertion consists of a duplication of two adjacent sequence motifs, resulting in a 18-residue insertion of new sequence after F71. (b) Frontal view of P91 with the active site cavity in the centre. The insertion lies at the junction between helix B and loop 5 (marked in red) which partyl covers the active site entrance. (c) The insertion variant has a ≈ 2-fold higher $k_{cat}$ for paraoxon than the wild type.

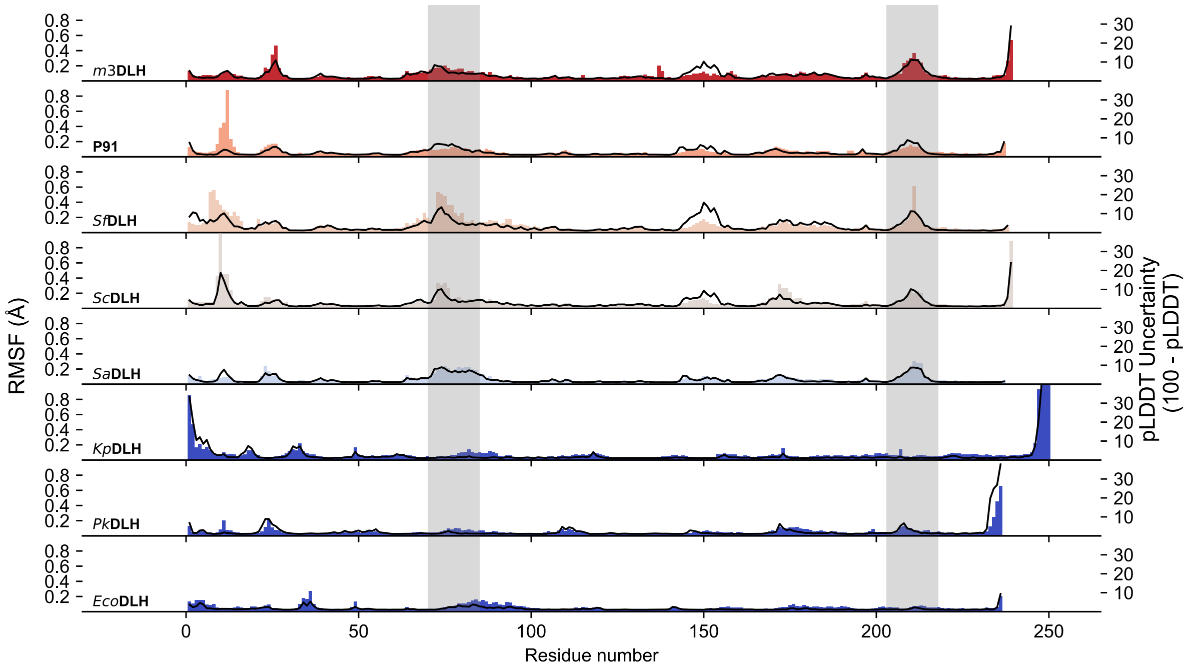

**Figure S19: AlphaFold 3 models predict the increased flexibility of active-site surrounding loops in enzymes with latent phosphotriesterase activity.** This figure presents a comparison of root mean square fluctuations (RMSF) and average inverted pLDDT scores derived from 50 overlayed AlphaFold 3 structural models of all enzymes for which experimental structures are available. The RMSF (coloured bars) reflects the predicted conformational diversity of α-carbon atoms, with higher values representing higher conformational diversity. The inverted pLDDT score (black line) serves as a measure of predicted local confidence, where lower values correspond to higher confidence in the local prediction of α-carbon atoms. The colour of each barplot corresponds to the enzyme's specific promiscuous phosphotriesterase activity. The structural features covering the active site, the C-terminal end of helix B and the adjacent stretch of loop 5, as well as loop 14, are highlighted in grey boxes. Note that the N-terminal 20 residues of *Kp*DLH were truncated for this analysis. Qualitatively, both metrics successfully predict the experimentally observed increased flexibility of the active site-surrounding loops in enzymes exhibiting promiscuous phosphotriesterase activity. However, these predictions do not demonstrate a quantitative correlation between the metrics and promiscuous activity within the closely related, active homologues.

### 3. Supplementary Tables

**Table S1: Overview of the enzymes tested in this study**, their sequence identities, source species, and environments.

| Enzyme | Sequence identity to P91 (%) | Species | Source environment | Reference |
|---|---|---|---|---|
| P91 | 100 | Unknown (metagenomic DNA identified in a functional metagenomic screening) | Ambiguous (pooled metagenomic libraries); either marine sludge or soil or compost (Netherlands) | (13, 15) |
| *Sif*DLH | 56 | *Sinimarinibacterium flocculans* | offshore surface sea water (South China Sea) | (16) |
| *m1*DLH | 56 | unknown (metagenomic DNA) | seawater from aquaculture system (Yantai, China) | MGnify Proteins Database accession number: MGYP00012671522; BioSamples Database accession number: SAMEA104210103 |
| *Sf*DLH | 57 | *Solimonas fluminis* | freshwater river (Han River in Seoul) | (17) |
| *Sc*DLH | 58 | *Solimonas fluminis* K1W22B-7 *(*previously Solimonas cavernae) | water in a karst cave (Guizhou, China) | (18) |
| *Sa*DLH | 57 | *Solimonas aquatica* | freshwater spring (Taiwan) | (19) |
| *Hm*DLH | 60 | *Mangrovimicrobium sediminis* (previously *Haliea* sp. SAOS-164 or *Haliea mangrovi*) | mangrove soil (Goa, India) | (20) |
| *m3*DLH | 65 | unknown (metagenomic DNA, genomic assembly: *Halieaceae* bacterium SZUA-439) | sulphide sediment from hydrothermal vent TVG12 on the South Atlantic Ridge | BioSamples accession number: SAMN09287987 |
| *Mz*DLH | 27 | *Marinobacter zhejiangensis* | sediment from the East China Sea | (21) |
| *Hs*DLH (CMBL) | 12 | *Homo sapiens* | -- | (22) |
| *Kp*DLH | 16 | *Klebsiella pneumoniae* | Ubiquitous (soil, water, plants, animals) | (23) |
| *Pk*DLH | 19 | *Pseudomonas* sp. strain B13 (*Pseudomonas knackmussii*) | sewage treatment plant (Göttingen, Germany) | (24, 25) |
| *Eco*DLH | 16 | *Escherichia coli* | Gut microbiomes of warm-blooded animals | (26) |

**Table S2: Phosphotriesterase kinetics of all enzymes characterised in this study**, measured in 50 mM HEPES-NaOH, 150 mM NaCl, pH 8.0 at 25 °C. All enzymes were purified with an N-terminal StrepII-tag, apart from *Hs*DLH variants, which carry a solubility tag (marked with an asterisk).

| Enzyme | Paraoxon-ethyl | | | Fluorescein di(diethylphosphate) | | | |
|---|---|---|---|---|---|---|---|
| | $k_{cat}$ ($s^{-1}$) | $K_M$ (mM) | $k_{cat}/K_M$ ($M^{-1}$ $s^{-1}$) | $k_{cat}$ ($s^{-1}$) | $K_M$ (μM) | $K_i$ (μM) | $k_{cat}/K_M$ ($M^{-1}$ $s^{-1}$) |
| P91 | 0.12 | 0.58 | 210 | 0.081 | 46 | 290 | 1800 |
| *Sif*DLH | 0.59 | 9.8 | 60 | not determined | | | |
| *m1*DLH | 0.20 | 4.6 | 44 | not determined | | | |
| *Sf*DLH | 0.19 | 5.6 | 35 | not determined | | | |
| *Sc*DLH | 0.11 | 7.6 | 14 | not determined | | | |
| *Sa*DLH | 0.014 | 3.5 | 3.9 | not determined | | | |
| *Hm*DLH | 14 | 1.4 | 10000 | not determined | | | |
| *m3*DLH | 6.5 | 1.1 | 5900 | not determined | | | |
| *Mz*DLH | 0.006 | 5.6 | 1.1 | $9.9 \times 10^{-4}$ | 16 | 750 | 62 |
| *Hs*DLH wild-type | no detectable activity | | | $1.6 \times 10^{-4}$ | 33 | 270 | 4.8 |
| *Hs*DLH wild-type* | no detectable activity | | | $2.2 \times 10^{-5}$ | 59 | -- | 0.37 |
| HsDLH-1E11* | no detectable activity | | | $1.2 \times 10^{-4}$ | 40 | 320 | 2.9 |
| *Kp*DLH | no detectable activity | | | $3.0 \times 10^{-5}$ | 40 | 400 | 0.75 |
| *Pk*DLH | no detectable activity | | | no detectable activity | | | |
| *Eco*DLH | no detectable activity | | | no detectable activity | | | |

* with N-terminal solubility-enhancing tag ($His_6$-TwinStep-SUMO)

**Table S3: Properties of substrates used to measure Michaelis-Menten kinetics.** Extinction coefficients were derived from calibration curves measured in the spectrophotometric setup that was used for reaction monitoring.

| Substrate | Leaving group/detected product | p$K_a$ | Detection wavelength (nm) | Extinction coefficient ($M^{-1}$) |
|---|---|---|---|---|
| 5 | 3-Fluoro-4-nitrophenol | 5.94 | 390 | 10473.2 |
| 1 | Fluorescein | 6.4 | 480 (excitation) / 520 (emission) | $4.17 \times 10^9$ |
| 3 | 3-(2-hydroxyphenyl)propanoic acid | 4.75 | 270 | 844.4 |
| 2 & 4 | 4-Nitrophenol | 7.14 | 405 | 10038.1 |
| 6 | 4-Hydroxybenzaldehyde | 7.66 | 330 | 12483.8 |
| 7 | 4-Cyanophenol | 7.95 | 275 | 6721.7 |
| 8 | 4-Hydroxyacetophenone | 8.05 | 320 | 7242.1 |
| 9 | 3-Cyanophenol | 8.61 | 295 | 1246.2 |
| 10 | 3-Chlorophenol | 9.12 | 276 | 796.0 |

**Table S4: Steady-state catalytic parameters for linear free-energy relationship of P91-WT**, measured in 50 mM HEPES-NaOH, 150 mM NaCl, 1 mM TCEP, pH 8.0.

| Substrate | p$K_a$ of leaving group | Enzyme concentration in the measurement | $k_{cat}$ ($s^{-1}$) | $K_M$ (mM) | $k_{cat}/K_M$ ($M^{-1}\cdot s^{-1}$) |
|---|---|---|---|---|---|
| 5 | 5.94 | 20 nM | 0.57 | 0.55 | 1000 |
| 2 | 7.14 | 0.5 µM | 0.12 | 0.58 | 210 |
| 6 | 7.66 | 1 µM | 0.017 | 1.1 | 15 |
| 7 | 7.95 | 4 µM | 0.0081 | 0.54 | 15 |
| 8 | 8.05 | 3 µM | 0.0038 | 0.9 | 4.2 |
| 9 | 8.61 | 3 µM | 0.0028 | 1.4 | 2.1 |
| 10 | 9.12 | 12.56 µM | no turnover could be detected | | |

**Table S5: Steady-state catalytic parameters for linear free-energy relationship of *Sc*DLH**, measured in 50 mM HEPES-NaOH, 150 mM NaCl, pH 8.0.

| Substrate | p$K_a$ of leaving group | Enzyme concentration in the measurement | $k_{cat}$ ($s^{-1}$) | $K_M$ (mM) | $k_{cat}/K_M$ ($M^{-1}\cdot s^{-1}$) |
|---|---|---|---|---|---|
| 5 | 5.94 | 100 nM | 0.36 | 2.7 | 130 |
| 2 | 7.14 | 5 µM | 0.11 | 7.6 | 14 |
| 6 | 7.66 | 1 µM | 0.0095 | 5.1 | 1.8 |
| 7 | 7.95 | 1 µM | 0.0032 | 3.8 | 0.86 |
| 8 | 8.05 | 2 µM | 0.015 | 33 | 0.44 |
| 9 | 8.61 | 3.91 µM | 0.001 | 7.9 | 0.13 |
| 10 | 9.12 | | no turnover could be detected | | |

**Table S6: Steady-state catalytic parameters for linear free-energy relationship of *m3*DLH**, measured in 50 mM HEPES-NaOH, 150 mM NaCl, pH 8.0.

| Substrate | p$K_a$ of leaving group | Enzyme concentration in the measurement | $k_{cat}$ ($s^{-1}$) | $K_M$ (mM) | $k_{cat}/K_M$ ($M^{-1} \cdot s^{-1}$) |
|---|---|---|---|---|---|
| 5 | 5.94 | 1 nM | 1.8 | 2.0 | 910 |
| 2 | 7.14 | 10 nM | 6.5 | 1.1 | 5900 |
| 6 | 7.66 | 10 nM | 0.45 | 2.3 | 170 |
| 7 | 7.95 | 10 nM | 0.21 | 3.3 | 64 |
| 8 | 8.05 | 100 nM | 0.16 | 3.5 | 46 |
| 9 | 8.61 | 500 nM | 0.022 | 1.9 | 12 |
| 10 | 9.12 | 6.48 μM | 0.00064 | 0.28 | 2.3 |

**Table S7: Overview of residues included into the combinatorial active site library of *Pk*DLH.**

| Position | Residue in *Pk*DLH | Consensus residue in P91/*Sf*DLH/*Sc*DLH | Rationale |
|---|---|---|---|
| 34 | A | F | Second shell |
| 35 | Q | P | Positioning of the tetrad Glu |
| 37 | I | A | Part of assumed oxyanion hole |
| 44 | M | A | Second shell |
| 64 | Y | H | Second shell |
| 88 | W | I | Prominently in entrance of the active site |
| 93 | M | W | Second shell |
| 122 | Y | F | Next to catalytic nucleophile & catalytic pentad |
| 124 | L | F | Next to catalytic nucleophile |
| 143 | G | T | Next to catalytic nucleophile |
| 144 | Y | F | Closeness to catalytic nucleophile |
| 145 | Y | H | (part of catalytic pentad!) |
| 147 | V | G | (positioning of pentad His) |
| 148 | G | L | Second shell |
| 150 | E | P | Second shell, helix breaker? |
| 172 | H | P | Next to triad Asp |
| 173 | F | L | Entrance of active site |
| 201 | G | V/A | Next to triad His |
| 205 | A | T | Interacting with tetrad |
| 206 | R | D/N | Next to/interacting with tetrad Glu |

**Table S8: Summary of fragments combinatorially grafted from P91 and its homologues into *Hs*DLH**.

| Fragment | Sequences | Origin |
|---|---|---|
| F1 | QDI | *Hs*DLH (residues Q49–I51) |
| | PEA | P91-WT/*Hm*DLH/*m3*DLH (residues P36–A38 in P91-WT) |
| | PEL | P91-R2 (residues P36–L38) |
| | PEG | P91-GGRGWV (residues P36–G38) |
| F2 | QLPNT | *Hs*DLH (residues Q55–T59 ) |
| | NDHA | P91-WT/*Hm*DLH/*m3*DLH (residues 42–45 in P91-WT) |
| F3 | YGIV | *Hs*DLH (residues Y155–V158) |
| | HGGL | P91/*Hm*DLH (residues 141–144 in P91-WT) |
| F4 | SGQTHGFVHRKREDCSPADKP | *Hs*DLH (residues S208–P228) |
| | GNALHSFTDPEADTRGMDGLAYDAR | *Hm*DLH (residues G195–R219) |
| | GNAAHSFTDPAADAHGMAGLAYEPL | *m3*DLH (residues G196–L220) |
| | GNAVHSFTDPLAGSHGIPGLAYDAT | P91-WT (residues G195–T219) |
| | GNAVHSFTDPLAGSHGWPGVAYDAT | P91-R2 (residues G195–T219) |

**Table S9: Data collection details and refinement statistics for crystal structures of *m3*DLH, *Sc*DLH, *Sf*DLH, and *Sa*DLH**.

| | ***m3*DLH** | ***Sc*DLH** | ***Sf*DLH** | ***Sa*DLH** |
|---|---|---|---|---|
| **Data Collection** | | | | |
| **Space group** | C 1 2 1 | P 1 $2_1$ 1 | P 1 $2_1$ 1 | P 1 $2_1$ 1 |
| **Cell dimensions** | | | | |
| ***a*, *b*, *c* (Å)** | 78.45, 43.68, 59.65 | 43.55, 123.60, 46.66 | 40.14, 87.84, 71.41 | 53.45, 76.41, 72.46 |
| **α, β, γ (°)** | 90.00, 92.71, 90.00 | 90.00, 100.98, 90.00 | 90.00, 99.17, 90.00 | 90.00, 107.77, 90.00 |
| **Resolution (Å)** | 23.75–1.533 (1.588–1.533) | 40.4–1.85 (1.916–1.85) | 39.63–1.45 (1.502–1.45) | 69.00–1.90 (1.968–1.90) |
| **$R_{sym}$ or $R_{merge}$** | 0.1138 (0.5259) | 0.08356 (0.5364) | 0.0669 (0.2875) | 0.04279 (0.1881) |
| **$CC_{1/2}$** | 0.996 (0.85) | 0.997 (0.909) | 0.998 (0.979) | 0.996 (0.924) |
| **CC*** | 0.999 (0.958) | 0.999 (0.976) | 0.999 (0.995) | 0.999 (0.98) |
| **$I / \sigma I$** | 9.53 (2.13) | 13.14 (3.64) | 12.01 (4.43) | 7.10 (2.75) |
| **Completeness (%)** | 98.36 (95.71) | 97.87 (99.76) | 99.73 (99.14) | 95.49 (95.13) |
| **Redundancy** | 6.7 (5.9) | 6.7 (6.8) | 6.5 (6.3) | 1.9 (1.9) |
| **Refinement** | | | | |
| **Resolution (Å)** | 23.75–1.533 (1.588–1.533) | 40.4–1.85 (1.916–1.85) | 39.63–1.45 (1.502–1.45) | 69.00–1.90 (1.968–1.90) |
| **No. unique reflections** | 29979 (2912) | 40398 (4112) | 86395 (8555) | 41833 (4118) |
| **$R_{work}$ / $R_{free}$** | 0.1638 / 0.2017 | 0.1646 / 0.1943 | 0.1431 / 0.1797 | 0.2425/0.2947 |
| **No. atoms:** | | | | |
| **Protein** | 1938 | 3690 | 3778 | 3626 |
| **Ligand/ion** | - | - | 61 | 2 |
| **Water** | 205 | 504 | 317 | 145 |
| ***B*-factors:** | | | | |
| **Protein** | 22.75 | 28.69 | 24.86 | 42.25 |
| **Ligand/ion** | - | - | 46.29 | 28.74 |
| **Water** | 30.79 | 38.24 | 32.25 | 43.27 |
| **R.m.s. deviations:** | | | | |

| | | | | |
|---|---|---|---|---|
| **Bond lengths (Å)** | 0.008 | 0.003 | 0.008 | 0.008 |
| **Bond angles (°)** | 0.98 | 0.63 | 1.10 | 1.01 |
| **PDB ID** | 7JKA | 7JIZ | 8VB3 | 9N2W |

### 5. Sequences

Sequences of all characterised proteins and all cloned plasmid constructs. Signal peptides, as predicted by SignalP-5.0,(27) are underlined in the protein sequences and were not included into the expression construct (*Mz*DLH, *Eco*DLH). All constructs were initially cloned with an N-terminal StrepII-tag which allows single-step affinity capture at very high purities on Strep-tactin resin. For high-yield purification, as required for transient-state kinetics, the StrepII-tag of *m3*DLH and *Sc*DLH was exchanged for an N-terminal $His_6$-tag for purification by Ni-NTA affinity chromatography. Due to low soluble expression of *Hs*DLH, an N-terminal His-TwinStrep-SUMO-tag was cloned into the construct for higher solubility. *Hs*DLH with this tag was also purified by Ni-NTA affinity chromatography. In the plasmid sequences the coding gene sequences are shown in uppercase, including the respective N-terminal affinity tags. The sequences of N-terminal tags are:

StrepII-tag:

MASWSHPQFEKGA

$His_6$-tag:

MHHHHHHGGS

His-TwinStrep-SUMO-tag:

MGSSHHHHHHSSGLVPRGSHMASWSHPQFEKGGGSGGGSGGSAWSHPQFEKMSDSEVNQEAKPEVKPE
VKPETHINLKVSDGSSEIFFKIKKTTPLRRLMEAFAKRQGKEMDSLRFLYDGIRIQADQTPEDLDMED
NDIIEAHREQIGGS

>*Hm*DLH

METRRITYSDGNTDYLGELYWEAGDAPRPGIVVFPEAFGLNDHARERARRLAAIGYVAFAADLHGGGA
ILDSMEALGPRMQALFADRTLWRAMAGAALDTLAAQAEADAGKLGAIGFCFGGATCLELARSGAALAG
IASFHGGLKEEIDGDAGRIRASVLVLHGAEDPLLGAGTIDAVMTEFRRDQVDWQFTYYGNALHSFTDP
EADTRGMDGLAYDARTEARSWNAMAAFFDELFN

>pASK-IBA5+_StrepII-*Hm*DLH

gattaattcctaattttgttgacactctatcattgatagagttattttaccactccctatcagtgat
agagaaaagtgaaatgaatagttcgacaaaaatctagaaataattttgtttaactttaagaaggagat
atacaaATGGCTAGCTGGAGCCACCCGCAGTTCGAAAAAGGCGCCATGGAAACCCGTCGTATTACCTA
TAGTGATGGCAATACCGATTATCTGGGTGAACTGTATTGGGAAGCCGGTGATGCACCGCGTCCGGGTA
TTGTTGTTTTTCCGGAAGCATTTGGTCTGAATGATCATGCACGTGAACGTGCCCGTCGTCTGGCAGCA
ATTGGTTATGTTGCATTTGCAGCCGATCTGCATGGTGGTGGTGCAATTCTGGATAGCATGGAAGCACT
GGGTCCGCGTATGCAGGCACTGTTTGCAGATCGTACCCTGTGGCGTGCAATGGCAGGCGCAGCACTGG
ATACCCTGGCAGCACAGGCCGAAGCAGATGCAGGTAAACTGGGTGCCATTGGTTTTTGTTTTGGTGGT
GCCACCTGTCTGGAACTGGCACGTAGCGGTGCAGCCCTGGCAGGTATTGCAAGCTTTCATGGTGGTCT
GAAAGAAGAAATTGACGGCGACGCAGGTCGTATTCGTGCAAGCGTTCTGGTTCTGCATGGCGCTGAAG
ATCCGCTGTTAGGTGCAGGCACCATTGATGCAGTTATGACCGAATTTCGTCGTGATCAGGTTGATTGG
CAGTTTACGTATTATGGTAATGCCCTGCATAGCTTTACCGATCCGGAAGCGGATACCCGTGGTATGGA
TGGTCTGGCATATGATGCACGTACCGAAGCACGTAGCTGGAACGCCATGGCAGCATTTTTTGATGAAC
TGTTCAACTAAtgatatctaactaagcttgacctgtgaagtgaaaaatggcgcacattgtgcgacatt
tttttgtctgccgtttaccgctactgcgtcacggatctccacgcgccctgtagcggcgcattaagcg
cggcgggtgtggtggttacgcgcagcgtgaccgctacacttgccagcgccctagcgcccgctcctttc
gctttcttcccttcctttctcgccacgttcgccggctttccccgtcaagctctaaatcgggggctccc
tttagggttccgatttagtgctttacggcacctcgaccccaaaaaacttgattagggtgatggttcac
gtagtgggccatcgccctgatagacggtttttcgccctttgacgttggagtccacgttctttaatagt
ggactcttgttccaaactggaacaacactcaaccctatctcggtctattcttttgatttataagggat
tttgccgatttcggcctattggttaaaaaatgagctgatttaacaaaaatttaacgcgaattttaaca
aaatattaacgcttacaatttcaggtggcacttttcggggaaatgtgcgcggaacccctatttgttta
tttttctaaatacattcaaatatgtatccgctcatgagacaataaccctgataaatgcttcaataata
ttgaaaaaggaagagtatgagtattcaacatttccgtgtcgcccttattcccttttttgcggcatttt
gccttcctgtttttgctcacccagaaacgctggtgaaagtaaaagatgctgaagatcagttgggtgca
cgagtgggttacatcgaactggatctcaacagcggtaagatccttgagagttttcgccccgaagaacg
ttttccaatgatgagcacttttaaagttctgctatgtggcgcggtattatcccgtattgacgccgggc
aagagcaactcggtcgccgcatacactattctcagaatgacttggttgagtactcaccagtcacagaa
aagcatcttacggatggcatgacagtaagagaattatgcagtgctgccataaccatgagtgataacac
tgcggccaacttacttctgacaacgatcggaggaccgaaggagctaaccgcttttttgcacaacatgg

gggatcatgtaactcgccttgatcgttgggaaccggagctgaatgaagccataccaaacgacgagcgt
gacaccacgatgcctgtagcaatggcaacaacgttgcgcaaactattaactggcgaactacttactct
agcttcccggcaacaattgatagactggatggaggcggataaagttgcaggaccacttctgcgctcgg
cccttccggctggctggtttattgctgataaatctggagccggtgagcgtggctctcgcggtatcatt
gcagcactggggccagatggtaagccctcccgtatcgtagttatctacacgacggggagtcaggcaac
tatggatgaacgaaatagacagatcgctgagataggtgcctcactgattaagcattggtaggaattaa
tgatgtctcgtttagataaaagtaaagtgattaacagcgcattagagctgcttaatgaggtcggaatc
gaaggtttaacaacccgtaaactcgcccagaagctaggtgtagagcagcctacattgtattggcatgt
aaaaaataagcgggctttgctcgacgccttagccattgagatgttagataggcaccatactcactttt
gccctttagaaggggaaagctggcaagatttttttacgtaataacgctaaaagttttagatgtgcttta
ctaagtcatcgcgatggagcaaaagtacatttaggtacacggcctacagaaaaacagtatgaaactct
cgaaaatcaattagcctttttatgccaacaaggtttttcactagagaatgcattatatgcactcagcg
cagtggggcattttactttaggttgcgtattggaagatcaagagcatcaagtcgctaaagaagaaagg
gaaacacctactactgatagtatgccgccattattacgacaagctatcgaattatttgatcaccaagg
tgcagagccagccttcttattcggccttgaattgatcatatgcggattagaaaaacaacttaaatgtg
aaagtgggtcttaaaagcagcataacctttttccgtgatggtaacttcactagtttaaaaggatctag
gtgaagatcctttttgataatctcatgaccaaaatcccttaacgtgagttttcgttccactgagcgtc
agaccccgtagaaaagatcaaaggatcttcttgagatcctttttttctgcgcgtaatctgctgcttgc
aaacaaaaaaaccaccgctaccagcggtggtttgtttgccggatcaagagctaccaactctttttccg
aaggtaactggcttcagcagagcgcagataccaaatactgtccttctagtgtagccgtagttaggcca
ccacttcaagaactctgtagcaccgcctacatacctcgctctgctaatcctgttaccagtggctgctg
ccagtggcgataagtcgtgtcttaccgggttggactcaagacgatagttaccggataaggcgcagcgg
tcgggctgaacggggggttcgtgcacacagcccagcttggagcgaacgacctacaccgaactgagata
cctacagcgtgagctatgagaaagcgccacgcttcccgaagggagaaaggcggacaggtatccggtaa
gcggcagggtcggaacaggagagcgcacgagggagcttccagggggaaacgcctggtatctttatagt
cctgtcgggtttcgccacctctgacttgagcgtcgatttttgtgatgctcgtcaggggggcggagcct
atggaaaaacgccagcaacgcggcctttttacggttcctggccttttgctggccttttgctcacatga
cccgacaccatcgaatggccagat

>*m3*DLH

MKHREIRYTDGHTQFVGELHWDEQQGGKCPGVVVFPEAFGLNDHARERARRLAGLGYAALAADLHGDG
RLIDDMEQLRPRMEGLFGDRAAWRALARAALDTLVAQPEVDADRLAAIGFCFGGTTALELARSGASLG
AIVTFHAGLLPELPEDAGRIRGRVLVCHGAEDPLVQKEAIDAVMGEWRRDRVDWQFTFYGNAAHSFTD
PAADAHGMAGLAYEPLTEARSWTAMRNLFDEVFSR

>pASK-IBA5+_StrepII-*m3*DLH

gattaattcctaatttttgttgacactctatcattgatagagttattttaccactccctatcagtgat
agagaaaagtgaaatgaatagttcgacaaaaatctagaaataattttgtttaactttaagaaggagat
atacaaATGGCTAGCTGGAGCCACCCGCAGTTCGAAAAAGGCGCCATGAAACATCGTGAAATTCGTTA
TACCGATGGCCATACACAGTTTGTTGGTGAACTGCATTGGGATGAACAGCAAGGTGGTAAATGTCCGG
GTGTTGTTGTTTTTCCGGAAGCATTTGGTCTGAATGATCATGCACGTGAACGTGCCCGTCGTCTGGCA
GGTCTGGGTTATGCAGCACTGGCAGCCGATCTGCATGGTGATGGTCGTCTGATTGATGATATGGAACA
GCTGCGTCCGCGTATGGAAGGTCTGTTTGGTGATCGTGCAGCATGGCGTGCACTGGCACGTGCAGCCC
TGGATACCCTGGTTGCACAGCCGGAAGTTGATGCCGATCGCCTGGCAGCAATTGGTTTTTGTTTTGGT
GGCACCACCGCACTGGAACTGGCACGCAGCGGTGCAAGCCTGGGTGCAATTGTTACCTTTCATGCAGG
TCTGCTGCCGGAACTGCCTGAAGATGCAGGTCGTATTCGTGGTCGTGTTCTGGTTTGTCATGGTGCAG
AAGATCCGCTGGTTCAGAAAGAAGCAATTGACGCAGTTATGGGTGAATGGCGTCGTGATCGTGTTGAT
TGGCAGTTTACCTTTTATGGTAATGCAGCACATAGCTTTACCGATCCGGCAGCGGATGCACATGGTAT
GGCAGGCCTGGCCTATGAACCGCTGACCGAAGCACGTAGCTGGACCGCAATGCGTAACCTGTTTGATG
AAGTTTTTAGCCGCTAAtgatatctaactaagcttgacctgtgaagtgaaaaatggcgcacattgtgc
gacatttttttgtctgccgtttaccgctactgcgtcacggatctccacgcgccctgtagcggcgcat
taagcgcggcgggtgtggtggttacgcgcagcgtgaccgctacacttgccagcgccctagcgcccgct
cctttcgctttcttcccttcctttctcgccacgttcgccggctttccccgtcaagctctaaatcgggg
gctccctttagggttccgatttagtgctttacggcacctcgaccccaaaaaacttgattagggtgatg
gttcacgtagtgggccatcgccctgatagacggtttttcgccctttgacgttggagtccacgttcttt
aatagtggactcttgttccaaactggaacaacactcaaccctatctcggtctattcttttgatttata
agggattttgccgatttcggcctattggttaaaaaatgagctgatttaacaaaaatttaacgcgaatt
ttaacaaaatattaacgcttacaatttcaggtggcacttttcggggaaatgtgcgcggaacccctatt
tgtttatttttctaaatacattcaaatatgtatccgctcatgagacaataaccctgataaatgcttca
ataatattgaaaaaggaagagtatgagtattcaacatttccgtgtcgcccttattcccttttttgcgg
cattttgccttcctgtttttgctcacccagaaacgctggtgaaagtaaaagatgctgaagatcagttg
ggtgcacgagtgggttacatcgaactggatctcaacagcggtaagatccttgagagttttcgccccga
agaacgttttccaatgatgagcacttttaaagttctgctatgtggcgcggtattatcccgtattgacg
ccgggcaagagcaactcggtcgccgcatacactattctcagaatgacttggttgagtactcaccagtc
acagaaaagcatcttacggatggcatgacagtaagagaattatgcagtgctgccataaccatgagtga
taacactgcggccaacttacttctgacaacgatcggaggaccgaaggagctaaccgcttttttgcaca

acatggggggatcatgtaactcgccttgatcgttgggaaccggagctgaatgaagccataccaaacgac
gagcgtgacaccacgatgcctgtagcaatggcaacaacgttgcgcaaactattaactggcgaactact
tactctagcttcccggcaacaattgatagactggatggaggcggataaagttgcaggaccacttctgc
gctcggcccttccggctggctggtttattgctgataaatctggagccggtgagcgtggctctcgcggt
atcattgcagcactggggccagatggtaagccctcccgtatcgtagttatctacacgacggggagtca
ggcaactatggatgaacgaaatagacagatcgctgagataggtgcctcactgattaagcattggtagg
aattaatgatgtctcgtttagataaaagtaaagtgattaacagcgcattagagctgcttaatgaggtc
ggaatcgaaggtttaacaacccgtaaactcgcccagaagctaggtgtagagcagcctacattgtattg
gcatgtaaaaaataagcgggctttgctcgacgccttagccattgagatgttagataggcaccatactc
acttttgccctttagaaggggaaagctggcaagattttttacgtaataacgctaaaagttttagatgt
gctttactaagtcatcgcgatggagcaaaagtacatttaggtacacggcctacagaaaaacagtatga
aactctcgaaaatcaattagccttttatgccaacaaggtttttcactagagaatgcattatatgcac
tcagcgcagtggggcattttactttaggttgcgtattggaagatcaagagcatcaagtcgctaaagaa
gaaagggaaacacctactactgatagtatgccgccattattacgacaagctatcgaattatttgatca
ccaaggtgcagagccagccttcttattcggccttgaattgatcatatgcggattagaaaaacaactta
aatgtgaaagtgggtcttaaaagcagcataacctttttccgtgatggtaacttcactagtttaaaagg
atctaggtgaagatcctttttgataatctcatgaccaaaatcccttaacgtgagttttcgttccactg
agcgtcagaccccgtagaaaagatcaaaggatcttcttgagatcctttttttctgcgcgtaatctgct
gcttgcaaacaaaaaaaccaccgctaccagcggtggtttgtttgccggatcaagagctaccaactctt
tttccgaaggtaactggcttcagcagagcgcagataccaaatactgtccttctagtgtagccgtagtt
aggccaccacttcaagaactctgtagcaccgcctacatacctcgctctgctaatcctgttaccagtgg
ctgctgccagtggcgataagtcgtgtcttaccgggttggactcaagacgatagttaccggataaggcg
cagcggtcgggctgaacggggggttcgtgcacacagcccagcttggagcgaacgacctacaccgaact
gagatacctacagcgtgagctatgagaaagcgccacgcttcccgaagggagaaaggcggacaggtatc
cggtaagcggcagggtcggaacaggagagcgcacgagggagcttccagggggaaacgcctggtatctt
tatagtcctgtcgggtttcgccacctctgacttgagcgtcgatttttgtgatgctcgtcaggggggcg
gagcctatggaaaaacgccagcaacgcggcctttttacggttcctggccttttgctggccttttgctc
acatgacccgacaccatcgaatggccagat

>pASK-IBA5+_6xHis-*m3*DLH
gattaattcctaatttttgttgacactctatcattgatagagttattttaccactccctatcagtgat
agagaaaagtgaaatgaatagttcgacaaaaatctagaaataattttgtttaactttaagaaggagat
atacaaATGCATCACCATCATCACCACGGTGGAATGAAACATCGTGAAATTCGTTATACCGATGGCCA
TACACAGTTTGTTGGTGAACTGCATTGGGATGAACAGCAAGGTGGTAAATGTCCGGGTGTTGTTGTTT
TTCCGGAAGCATTTGGTCTGAATGATCATGCACGTGAACGTGCCCGTCGTCTGGCAGGTCTGGGTTAT
GCAGCACTGGCAGCCGATCTGCATGGTGATGGTCGTCTGATTGATGATATGGAACAGCTGCGTCCGCG
TATGGAAGGTCTGTTTGGTGATCGTGCAGCATGGCGTGCACTGGCACGTGCAGCCCTGGATACCCTGG

TTGCACAGCCGGAAGTTGATGCCGATCGCCTGGCAGCAATTGGTTTTTGTTTTGGTGGCACCACCGCA
CTGGAACTGGCACGCAGCGGTGCAAGCCTGGGTGCAATTGTTACCTTTCATGCAGGTCTGCTGCCGGA
ACTGCCTGAAGATGCAGGTCGTATTCGTGGTCGTGTTCTGGTTTGTCATGGTGCAGAAGATCCGCTGG
TTCAGAAAGAAGCAATTGACGCAGTTATGGGTGAATGGCGTCGTGATCGTGTTGATTGGCAGTTTACC
TTTTATGGTAATGCAGCACATAGCTTTACCGATCCGGCAGCGGATGCACATGGTATGGCAGGCCTGGC
CTATGAACCGCTGACCGAAGCACGTAGCTGGACCGCAATGCGTAACCTGTTTGATGAAGTTTTTAGCC
GCTAAtgatatctaactaagcttgacctgtgaagtgaaaaatggcgcacattgtgcgacattttttt
gtctgccgtttaccgctactgcgtcacggatctccacgcgccctgtagcggcgcattaagcgcggcgg
gtgtggtggttacgcgcagcgtgaccgctacacttgccagcgccctagcgcccgctcctttcgctttc
ttcccttcctttctcgccacgttcgccggctttccccgtcaagctctaaatcgggggctccctttagg
gttccgatttagtgctttacggcacctcgaccccaaaaaacttgattagggtgatggttcacgtagtg
ggccatcgccctgatagacggtttttcgccctttgacgttggagtccacgttctttaatagtggactc
ttgttccaaactggaacaacactcaaccctatctcggtctattcttttgatttataagggattttgcc
gatttcggcctattggttaaaaaatgagctgatttaacaaaaatttaacgcgaattttaacaaaatat
taacgcttacaatttcaggtggcacttttcggggaaatgtgcgcggaacccctatttgtttatttttc
taaatacattcaaatatgtatccgctcatgagacaataaccctgataaatgcttcaataatattgaaa
aaggaagagtatgagtattcaacatttccgtgtcgcccttattcccttttttgcggcattttgccttc
ctgtttttgctcacccagaaacgctggtgaaagtaaaagatgctgaagatcagttgggtgcacgagtg
ggttacatcgaactggatctcaacagcggtaagatccttgagagttttcgccccgaagaacgttttcc
aatgatgagcacttttaaagttctgctatgtggcgcggtattatcccgtattgacgccgggcaagagc
aactcggtcgccgcatacactattctcagaatgacttggttgagtactcaccagtcacagaaaagcat
cttacggatggcatgacagtaagagaattatgcagtgctgccataaccatgagtgataacactgcggc
caacttacttctgacaacgatcggaggaccgaaggagctaaccgcttttttgcacaacatgggggatc
atgtaactcgccttgatcgttgggaaccggagctgaatgaagccataccaaacgacgagcgtgacacc
acgatgcctgtagcaatggcaacaacgttgcgcaaactattaactggcgaactacttactctagcttc
ccggcaacaattgatagactggatggaggcggataaagttgcaggaccacttctgcgctcggcccttc
cggctggctggtttattgctgataaatctggagccggtgagcgtggctctcgcggtatcattgcagca
ctggggccagatggtaagccctcccgtatcgtagttatctacacgacggggagtcaggcaactatgga
tgaacgaaatagacagatcgctgagataggtgcctcactgattaagcattggtaggaattaatgatgt
ctcgtttagataaaagtaaagtgattaacagcgcattagagctgcttaatgaggtcggaatcgaaggt
ttaacaacccgtaaactcgcccagaagctaggtgtagagcagcctacattgtattggcatgtaaaaaa
taagcgggctttgctcgacgccttagccattgagatgttagataggcaccatactcacttttgccctt
tagaaggggaaagctggcaagattttttacgtaataacgctaaaagttttagatgtgctttactaagt
catcgcgatggagcaaaagtacatttaggtacacggcctacagaaaaacagtatgaaactctcgaaaa
tcaattagcctttttatgccaacaaggtttttcactagagaatgcattatatgcactcagcgcagtgg
ggcattttactttaggttgcgtattggaagatcaagagcatcaagtcgctaaagaagaaaggaaaca
cctactactgatagtatgccgccattattacgacaagctatcgaattatttgatcaccaaggtgcaga

gccagccttcttattcggccttgaattgatcatatgcggattagaaaaacaacttaaatgtgaaagtg
ggtcttaaaagcagcataacctttttccgtgatggtaacttcactagtttaaaggatctaggtgaag
atcctttttgataatctcatgaccaaaatcccttaacgtgagttttcgttccactgagcgtcagaccc
cgtagaaaagatcaaaggatcttcttgagatcctttttttctgcgcgtaatctgctgcttgcaaacaa
aaaaaccaccgctaccagcggtggtttgtttgccggatcaagagctaccaactctttttccgaaggta
actggcttcagcagagcgcagataccaaatactgtccttctagtgtagccgtagttaggccaccactt
caagaactctgtagcaccgcctacatacctcgctctgctaatcctgttaccagtggctgctgccagtg
gcgataagtcgtgtcttaccgggttggactcaagacgatagttaccggataaggcgcagcggtcgggc
tgaacggggggttcgtgcacacagcccagcttggagcgaacgacctacaccgaactgagatacctaca
gcgtgagctatgagaaagcgccacgcttcccgaagggagaaaggcggacaggtatccggtaagcggca
gggtcggaacaggagagcgcacgagggagcttccagggggaaacgcctggtatctttatagtcctgtc
gggtttcgccacctctgacttgagcgtcgatttttgtgatgctcgtcaggggggcggagcctatggaa
aaacgccagcaacgcggcctttttacggttcctggccttttgctggccttttgctcacatgacccgac
accatcgaatggccagat

>P91

MTARKVDYTDGATRCIGEFHWDEGKSGPRPGVVVFPEAFGLNDHAKERARRLADLGFAALAADMHGDA
QVFDAASLSSTIQGYYGDRAHWRRRAQAALDALTAQPEVDGSKVAAIGFCFGGATCLELARTGAPLTA
IVTFHGGLLPEMAGDAGRIQSSVLVCHGADDPLVQDETMKAVMDEFRRDKVDWQVLYLGNAVHSFTDP
LAGSHGIPGLAYDATAEARSWTAMCNLFSELFG

>pASK-IBA5+_StrepII-P91

gattaattcctaatttttgttgacactctatcattgatagagttattttaccactccctatcagtgat
agagaaaagtgaaatgaatagttcgacaaaaatctagaaataattttgtttaactttaagaaggagat
atacaaATGGCTAGCTGGAGCCACCCGCAGTTCGAAAAAGGCGCCATGACAGCAAGAAAAGTCGACTA
CACAGACGGTGCAACCCGCTGTATCGGTGAGTTTCATTGGGATGAAGGCAAGTCGGGCCCGCGTCCCG
GCGTGGTGGTCTTTCCCGAGGCTTTCGGCCTCAACGACCATGCCAAGGAGCGCGCGCGGCGCCTTGCC
GACCTCGGCTTTGCAGCCCTGGCGGCGGATATGCACGGAGACGCCCAGGTTTTCGATGCGGCGAGTCT
CTCATCAACCATACAGGGCTACTACGGCGACCGCGCCCACTGGCGACGTCGTGCGCAGGCAGCGCTCG
ATGCACTGACGGCACAGCCAGAGGTGGACGGCAGCAAGGTGGCGGCCATCGGCTTTTGTTTCGGCGGT
GCCACCTGCCTTGAACTGGCCCGCACAGGTGCGCCGCTGACCGCCATTGTCACCTTCCACGGCGGTTT
GCTGCCGGAGATGGCAGGCGATGCCGGACGGATCCAGTCCAGTGTTCTGGTGTGCCATGGCGCTGATG
ATCCGCTCGTACAGGACGAAACCATGAAGGCCGTCATGGACGAGTTTCGTCGCGACAAGGTGGATTGG
CAGGTGCTCTACCTCGGAAATGCGGTACACAGTTTCACCGATCCACTCGCTGGCAGTCACGGCATACC
CGGGCTGGCCTATGACGCCACTGCCGAAGCCCGGTCGTGGACGGCCATGTGCAATCTGTTCAGTGAAC
TGTTCGGCTGAtgatatctaactaagcttgacctgtgaagtgaaaaatggcgcacattgtgcgacatt
tttttgtctgccgtttaccgctactgcgtcacggatctccacgcgccctgtagcggcgcattaagcg
cggcgggtgtggtggttacgcgcagcgtgaccgctacacttgccagcgccctagcgcccgctcctttc
gctttcttcccttcctttctcgccacgttcgccggctttccccgtcaagctctaaatcgggggctccc
tttagggttccgatttagtgctttacggcacctcgaccccaaaaaacttgattagggtgatggttcac
gtagtgggccatcgccctgatagacggtttttcgccctttgacgttggagtccacgttctttaatagt
ggactcttgttccaaactggaacaacactcaaccctatctcggtctattcttttgatttataagggat
tttgccgatttcggcctattggttaaaaaatgagctgatttaacaaaaatttaacgcgaattttaaca
aaatattaacgcttacaatttcaggtggcacttttcggggaaatgtgcgcggaacccctatttgttta
tttttctaaatacattcaaatatgtatccgctcatgagacaataaccctgataaatgcttcaataata
ttgaaaaaggaagagtatgagtattcaacatttccgtgtcgcccttattcccttttttgcggcatttt
gccttcctgtttttgctcacccagaaacgctggtgaaagtaaaagatgctgaagatcagttgggtgca
cgagtgggttacatcgaactggatctcaacagcggtaagatccttgagagttttcgccccgaagaacg
ttttccaatgatgagcacttttaaagttctgctatgtggcgcggtattatcccgtattgacgccgggc
aagagcaactcggtcgccgcatacactattctcagaatgacttggttgagtactcaccagtcacagaa
aagcatcttacggatggcatgacagtaagagaattatgcagtgctgccataaccatgagtgataacac
tgcggccaacttacttctgacaacgatcggaggaccgaaggagctaaccgcttttttgcacaacatgg

gggatcatgtaactcgccttgatcgttgggaaccggagctgaatgaagccataccaaacgacgagcgt
gacaccacgatgcctgtagcaatggcaacaacgttgcgcaaactattaactggcgaactacttactct
agcttcccggcaacaattgatagactggatggaggcggataaagttgcaggaccacttctgcgctcgg
cccttccggctggctggtttattgctgataaatctggagccggtgagcgtggctctcgcggtatcatt
gcagcactggggccagatggtaagccctcccgtatcgtagttatctacacgacggggagtcaggcaac
tatggatgaacgaaatagacagatcgctgagataggtgcctcactgattaagcattggtaggaattaa
tgatgtctcgtttagataaaagtaaagtgattaacagcgcattagagctgcttaatgaggtcggaatc
gaaggtttaacaacccgtaaactcgcccagaagctaggtgtagagcagcctacattgtattggcatgt
aaaaaataagcgggctttgctcgacgccttagccattgagatgttagataggcaccatactcactttt
gccctttagaaggggaaagctggcaagattttttacgtaataacgctaaaagttttagatgtgcttta
ctaagtcatcgcgatggagcaaaagtacatttaggtacacggcctacagaaaaacagtatgaaactct
cgaaaatcaattagcctttttatgccaacaaggtttttcactagagaatgcattatatgcactcagcg
cagtggggcattttactttaggttgcgtattggaagatcaagagcatcaagtcgctaaagaagaaagg
gaaacacctactactgatagtatgccgccattattacgacaagctatcgaattatttgatcaccaagg
tgcagagccagccttcttattcggccttgaattgatcatatgcggattagaaaaacaacttaaatgtg
aaagtgggtcttaaaagcagcataacctttttccgtgatggtaacttcactagtttaaaaggatctag
gtgaagatcctttttgataatctcatgaccaaaatcccttaacgtgagttttcgttccactgagcgtc
agaccccgtagaaaagatcaaaggatcttcttgagatcctttttttctgcgcgtaatctgctgcttgc
aaacaaaaaaaccaccgctaccagcggtggtttgtttgccggatcaagagctaccaactctttttccg
aaggtaactggcttcagcagagcgcagataccaaatactgtccttctagtgtagccgtagttaggcca
ccacttcaagaactctgtagcaccgcctacatacctcgctctgctaatcctgttaccagtggctgctg
ccagtggcgataagtcgtgtcttaccgggttggactcaagacgatagttaccggataaggcgcagcgg
tcgggctgaacggggggttcgtgcacacagcccagcttggagcgaacgacctacaccgaactgagata
cctacagcgtgagctatgagaaagcgccacgcttcccgaagggagaaaggcggacaggtatccggtaa
gcggcagggtcggaacaggagagcgcacgagggagcttccagggggaaacgcctggtatctttatagt
cctgtcgggtttcgccacctctgacttgagcgtcgatttttgtgatgctcgtcaggggggcggagcct
atggaaaaacgccagcaacgcggcctttttacggttcctggccttttgctggccttttgctcacatga
cccgacaccatcgaatggccagat

>pASK-IBA5+_6xHis-P91
gattaattcctaatttttgttgacactctatcattgatagagttattttaccactccctatcagtgat
agagaaaagtgaaatgaatagttcgacaaaaatctagaaataattttgtttaactttaagaaggagat
atacaaATGCATCACCATCATCACCACGGTGGAATGACAGCAAGAAAAGTCGACTACACAGACGGTGC
AACCCGCTGTATCGGTGAGTTTCATTGGGATGAAGGCAAGTCGGGCCCGCGTCCCGGCGTGGTGGTCT
TTCCCGAGGCTTTCGGCCTCAACGACCATGCCAAGGAGCGCGCGCGGCGCCTTGCCGACCTCGGCTTT
GCAGCCCTGGCGGCGGATATGCACGGAGACGCCCAGGTTTTCGATGCGGCGAGTCTCTCATCAACCAT
ACAGGGCTACTACGGCGACCGCGCCCACTGGCGACGTCGTGCGCAGGCAGCGCTCGATGCACTGACGG

CACAGCCAGAGGTGGACGGCAGCAAGGTGGCGGCCATCGGCTTTTGTTTCGGCGGTGCCACCTGCCTT
GAACTGGCCCGCACAGGTGCGCCGCTGACCGCCATTGTCACCTTCCACGGCGGTTTGCTGCCGGAGAT
GGCAGGCGATGCCGGACGGATCCAGTCCAGTGTTCTGGTGTGCCATGGCGCTGATGATCCGCTCGTAC
AGGACGAAACCATGAAGGCCGTCATGGACGAGTTTCGTCGCGACAAGGTGGATTGGCAGGTGCTCTAC
CTCGGAAATGCGGTACACAGTTTCACCGATCCACTCGCTGGCAGTCACGGCATACCCGGGCTGGCCTA
TGACGCCACTGCCGAAGCCCGGTCGTGGACGGCCATGTGCAATCTGTTCAGTGAACTGTTCGGCTGAt
gatatctaactaagcttgacctgtgaagtgaaaaatggcgcacattgtgcgacatttttttgtctgc
cgtttaccgctactgcgtcacggatctccacgcgccctgtagcggcgcattaagcgcggcgggtgtgg
tggttacgcgcagcgtgaccgctacacttgccagcgccctagcgcccgctcctttcgctttcttccct
tcctttctcgccacgttcgccggctttccccgtcaagctctaaatcgggggctccctttagggttccg
atttagtgctttacggcacctcgaccccaaaaaacttgattagggtgatggttcacgtagtgggccat
cgccctgatagacggtttttcgccctttgacgttggagtccacgttctttaatagtggactcttgttc
caaactggaacaacactcaaccctatctcggtctattcttttgatttataagggattttgccgatttc
ggcctattggttaaaaaatgagctgatttaacaaaaatttaacgcgaattttaacaaaatattaacgc
ttacaatttcaggtggcacttttcggggaaatgtgcgcggaacccctatttgtttatttttctaaata
cattcaaatatgtatccgctcatgagacaataaccctgataaatgcttcaataatattgaaaaaggaa
gagtatgagtattcaacatttccgtgtcgcccttattcccttttttgcggcattttgccttcctgttt
ttgctcacccagaaacgctggtgaaagtaaaagatgctgaagatcagttgggtgcacgagtgggttac
atcgaactggatctcaacagcggtaagatccttgagagttttcgccccgaagaacgttttccaatgat
gagcacttttaaagttctgctatgtggcgcggtattatcccgtattgacgccgggcaagagcaactcg
gtcgccgcatacactattctcagaatgacttggttgagtactcaccagtcacagaaaagcatcttacg
gatggcatgacagtaagagaattatgcagtgctgccataaccatgagtgataacactgcggccaactt
acttctgacaacgatcggaggaccgaaggagctaaccgcttttttgcacaacatgggggatcatgtaa
ctcgccttgatcgttgggaaccggagctgaatgaagccataccaaacgacgagcgtgacaccacgatg
cctgtagcaatggcaacaacgttgcgcaaactattaactggcgaactacttactctagcttcccggca
acaattgatagactggatggaggcggataaagttgcaggaccacttctgcgctcggcccttccggctg
gctggtttattgctgataaatctggagccggtgagcgtggctctcgcggtatcattgcagcactgggg
ccagatggtaagccctcccgtatcgtagttatctacacgacggggagtcaggcaactatggatgaacg
aaatagacagatcgctgagataggtgcctcactgattaagcattggtaggaattaatgatgtctcgtt
tagataaaagtaaagtgattaacagcgcattagagctgcttaatgaggtcggaatcgaaggtttaaca
acccgtaaactcgcccagaagctaggtgtagagcagcctacattgtattggcatgtaaaaaataagcg
ggctttgctcgacgccttagccattgagatgttagataggcaccatactcacttttgccctttagaag
gggaaagctggcaagatttttttacgtaataacgctaaaagttttagatgtgctttactaagtcatcgc
gatggagcaaaagtacatttaggtacacggcctacagaaaaacagtatgaaactctcgaaaatcaatt
agcctttttatgccaacaaggtttttcactagagaatgcattatatgcactcagcgcagtggggcatt
ttactttaggttgcgtattggaagatcaagagcatcaagtcgctaaagaagaaagggaaacacctact
actgatagtatgccgccattattacgacaagctatcgaattatttgatcaccaaggtgcagagccagc

cttcttattcggccttgaattgatcatatgcggattagaaaaacaacttaaatgtgaaagtgggtctt
aaaagcagcataacctttttccgtgatggtaacttcactagtttaaaaggatctaggtgaagatcctt
tttgataatctcatgaccaaaatcccttaacgtgagttttcgttccactgagcgtcagaccccgtaga
aaagatcaaaggatcttcttgagatcctttttttctgcgcgtaatctgctgcttgcaaacaaaaaaac
caccgctaccagcggtggtttgtttgccggatcaagagctaccaactctttttccgaaggtaactggc
ttcagcagagcgcagataccaaatactgtccttctagtgtagccgtagttaggccaccacttcaagaa
ctctgtagcaccgcctacatacctcgctctgctaatcctgttaccagtggctgctgccagtggcgata
agtcgtgtcttaccgggttggactcaagacgatagttaccggataaggcgcagcggtcgggctgaacg
gggggttcgtgcacacagcccagcttggagcgaacgacctacaccgaactgagatacctacagcgtga
gctatgagaaagcgccacgcttcccgaagggagaaaggcggacaggtatccggtaagcggcagggtcg
gaacaggagagcgcacgagggagcttccagggggaaacgcctggtatctttatagtcctgtcgggttt
cgccacctctgacttgagcgtcgatttttgtgatgctcgtcaggggggcggagcctatggaaaaacgc
cagcaacgcggcctttttacggttcctggccttttgctggccttttgctcacatgacccgacaccatc
gaatggccagat

>P91-R2
MTARKVDYTDGATRCIGEFHWDEGKSGPRPGVVVFPELFGLNDHAKERARRLADLGFAALAADMHGDA
QVFDEASVSSTIQGYYGDRAHWRRRAQAALDALTAQPEVDGSKVAAIGFCFGGATCLELARTGAPLTA
IVTFHGGLLPEMAGDAGRIQSSVLVCHGADDPLVQDETMKAVMDEFRRDKVDWQVLYLGNAVHSFTDP
LAGSHGWPGVAYDATAEARSWTAMCNLFSELFG

>pASK-IBA5+_6xHis-P91-R2
gattaattcctaatttttgttgacactctatcattgatagagttattttaccactccctatcagtgat
agagaaaagtgaaatgaatagttcgacaaaaatctagaaataattttgtttaactttaagaaggagat
atacaaATGCATCACCATCATCACCACGGTGGAAGTATGACAGCAAGAAAAGTCGACTACACAGACGG
TGCAACCCGCTGTATCGGTGAGTTTCATTGGGATGAAGGCAAGTCGGGCCCGCGTCCCGGCGTGGTGG
TCTTTCCCGAGCTGTTCGGCCTCAACGACCATGCCAAGGAGCGCGCGCGGCGCCTTGCCGACCTCGGC
TTTGCAGCCCTGGCGGCGGATATGCACGGAGACGCCCAGGTTTTCGATGAGGCGAGTGTGTCATCAAC
CATACAGGGCTACTACGGCGACCGCGCCCACTGGCGACGTCGTGCGCAGGCAGCGCTCGATGCACTGA
CGGCACAGCCAGAGGTGGACGGCAGCAAGGTGGCGGCCATCGGCTTTTGTTTCGGCGGTGCGACCTGC
CTTGAACTGGCCCGCACAGGTGCGCCGCTGACCGCCATTGTCACCTTCCACGGCGGTTTGCTGCCGGA
GATGGCAGGCGATGCCGGACGGATCCAGTCCAGTGTTCTGGTGTGCCATGGCGCTGATGATCCGCTCG
TACAGGACGAAACCATGAAGGCCGTCATGGACGAGTTTCGTCGCGACAAGGTGGATTGGCAGGTGCTC
TACCTCGGAAATGCGGTACACAGTTTCACCGATCCACTCGCTGGCAGTCACGGCTGGCCCGGGGTTGC
CTATGACGCCACTGCCGAAGCCCGGTCGTGGACGGCCATGTGCAATCTGTTCAGTGAACTGTTCGGCT
GAtgatatctaactaagcttgacctgtgaagtgaaaaatggcgcacattgtgcgacatttttttgtc
tgccgtttaccgctactgcgtcacggatctccacgcgccctgtagcggcgcattaagcgcggcgggtg

tggtggttacgcgcagcgtgaccgctacacttgccagcgccctagcgcccgctcctttcgctttcttc
ccttcctttctcgccacgttcgccggctttccccgtcaagctctaaatcgggggctccctttagggtt
ccgatttagtgctttacggcacctcgaccccaaaaaacttgattagggtgatggttcacgtagtgggc
catcgccctgatagacggtttttcgccctttgacgttggagtccacgttctttaatagtggactcttg
ttccaaactggaacaacactcaaccctatctcggtctattcttttgatttataagggattttgccgat
ttcggcctattggttaaaaaatgagctgatttaacaaaaatttaacgcgaattttaacaaaatattaa
cgcttacaatttcaggtggcacttttcggggaaatgtgcgcggaacccctatttgtttatttttctaa
atacattcaaatatgtatccgctcatgagacaataaccctgataaatgcttcaataatattgaaaaag
gaagagtatgagtattcaacatttccgtgtcgcccttattcccttttttgcggcattttgccttcctg
tttttgctcacccagaaacgctggtgaaagtaaaagatgctgaagatcagttgggtgcacgagtgggt
tacatcgaactggatctcaacagcggtaagatccttgagagttttcgccccgaagaacgttttccaat
gatgagcacttttaaagttctgctatgtggcgcggtattatcccgtattgacgccgggcaagagcaac
tcggtcgccgcatacactattctcagaatgacttggttgagtactcaccagtcacagaaaagcatctt
acggatggcatgacagtaagagaattatgcagtgctgccataaccatgagtgataacactgcggccaa
cttacttctgacaacgatcggaggaccgaaggagctaaccgcttttttgcacaacatgggggatcatg
taactcgccttgatcgttgggaaccggagctgaatgaagccataccaaacgacgagcgtgacaccacg
atgcctgtagcaatggcaacaacgttgcgcaaactattaactggcgaactacttactctagcttcccg
gcaacaattgatagactggatggaggcggataaagttgcaggaccacttctgcgctcggcccttccgg
ctggctggtttattgctgataaatctggagccggtgagcgtggctctcgcggtatcattgcagcactg
gggccagatggtaagccctcccgtatcgtagttatctacacgacggggagtcaggcaactatggatga
acgaaatagacagatcgctgagataggtgcctcactgattaagcattggtaggaattaatgatgtctc
gtttagataaaagtaaagtgattaacagcgcattagagctgcttaatgaggtcggaatcgaaggttta
acaacccgtaaactcgcccagaagctaggtgtagagcagcctacattgtattggcatgtaaaaaataa
gcgggctttgctcgacgccttagccattgagatgttagataggcaccatactcacttttgccctttag
aaggggaaagctggcaagattttttacgtaataacgctaaaagttttagatgtgctttactaagtcat
cgcgatggagcaaaagtacatttaggtacacggcctacagaaaaacagtatgaaactctcgaaaatca
attagcctttttatgccaacaaggtttttcactagagaatgcattatatgcactcagcgcagtggggc
attttactttaggttgcgtattggaagatcaagagcatcaagtcgctaaagaagaaagggaaacacct
actactgatagtatgccgccattattacgacaagctatcgaattatttgatcaccaaggtgcagagcc
agccttcttattcggccttgaattgatcatatgcggattagaaaaacaacttaaatgtgaaagtgggt
cttaaaagcagcataacctttttccgtgatggtaacttcactagtttaaaaggatctaggtgaagatc
ctttttgataatctcatgaccaaaatcccttaacgtgagttttcgttccactgagcgtcagaccccgt
agaaaagatcaaaggatcttcttgagatcctttttttctgcgcgtaatctgctgcttgcaaacaaaaa
aaccaccgctaccagcggtggtttgtttgccggatcaagagctaccaactctttttccgaaggtaact
ggcttcagcagagcgcagataccaaatactgtccttctagtgtagccgtagttaggccaccacttcaa
gaactctgtagcaccgcctacatacctcgctctgctaatcctgttaccagtggctgctgccagtggcg
ataagtcgtgtcttaccgggttggactcaagacgatagttaccggataaggcgcagcggtcgggctga

acgggggttcgtgcacacagcccagcttggagcgaacgacctacaccgaactgagatacctacagcg
tgagctatgagaaagcgccacgcttcccgaagggagaaaggcggacaggtatccggtaagcggcaggg
tcggaacaggagagcgcacgagggagcttccagggggaaacgcctggtatctttatagtcctgtcggg
tttcgccacctctgacttgagcgtcgatttttgtgatgctcgtcaggggggcggagcctatggaaaaa
cgccagcaacgcggcctttttacggttcctggccttttgctggccttttgctcacatgacccgacacc
atcgaatggccagat

>P91 insertion variant
MTARKVDYTDGATRCIGEFHWDEGKSGPRPGVVVFPEAFGLNDHAKERARRLADLGFAALAADMHGDA
QVFTAASLSSTNMHGDAQVFQAASLSSTIQGYYGDRAHWRRRAQAALDALTAQPEVDGSKVAAIGFCF
GGATCLELARTGAPLTAIVTFHGGLLPEMAGDAGRIQSSVLVCHGADDPLVQDETMKAVMDEFRRDKV
DWQVLYLGNAVHSFTDPLAGSHGIPGLAYDATAEARSWTAMCNLFSELFG

>pASK-IBA5+_StrepII-P91-insertion-variant
gattaattcctaatttttgttgacactctatcattgatagagttattttaccactccctatcagtgat
agagaaaagtgaaatgaatagttcgacaaaaatctagaaataattttgtttaactttaagaaggagat
atacaaATGGCTAGCTGGAGCCACCCGCAGTTCGAAAAAGGCGCCATGACAGCAAGAAAAGTCGACTA
CACAGACGGTGCAACCCGCTGTATCGGTGAGTTTCATTGGGATGAAGGCAAGTCGGGCCCGCGTCCCG
GCGTGGTGGTCTTTCCCGAGGCTTTCGGCCTCAACGACCATGCCAAGGAGCGCGCGCGGCGCCTTGCC
GACCTCGGCTTTGCAGCCCTGGCGGCGGATATGCACGGAGACGCCCAGGTTTTCACGGCGGCGAGTCT
CTCATCAACCAATATGCACGGAGACGCCCAGGTTTTCCAGGCGGCGAGTCTCTCATCAACCATACAGG
GCTACTACGGCGACCGCGCCCACTGGCGACGTCGTGCGCAGGCAGCGCTCGATGCACTGACGGCACAG
CCAGAGGTGGACGGCAGCAAGGTGGCGGCCATCGGCTTTTGTTTCGGCGGTGCCACCTGCCTTGAACT
GGCCCGCACAGGTGCGCCGCTGACCGCCATTGTCACCTTCCACGGCGGTTTGCTGCCGGAGATGGCAG
GCGATGCCGGACGGATCCAGTCCAGTGTTCTGGTGTGCCATGGCGCTGATGATCCGCTCGTACAGGAC
GAAACCATGAAGGCCGTCATGGACGAGTTTCGTCGCGACAAGGTGGATTGGCAGGTGCTCTACCTCGG
AAATGCGGTACACAGTTTCACCGATCCACTCGCTGGCAGTCACGGCATACCCGGGCTGGCCTATGACG
CCACTGCCGAAGCCCGGTCGTGGACGGCCATGTGCAATCTGTTCAGTGAACTGTTCGGCTGAtgatat
ctaactaagcttgacctgtgaagtgaaaaatggcgcacattgtgcgacatttttttgtctgccgttt
accgctactgcgtcacggatctccacgcgccctgtagcggcgcattaagcgcggcgggtgtggtggtt
acgcgcagcgtgaccgctacacttgccagcgccctagcgcccgctcctttcgctttcttcccttcctt
tctcgccacgttcgccggctttccccgtcaagctctaaatcgggggctccctttagggttccgattta
gtgctttacggcacctcgaccccaaaaaacttgattagggtgatggttcacgtagtgggccatcgccc
tgatagacggtttttcgccctttgacgttggagtccacgttctttaatagtggactcttgttccaaac
tggaacaacactcaaccctatctcggtctattcttttgatttataagggattttgccgatttcggcct
attggttaaaaaatgagctgatttaacaaaaatttaacgcgaattttaacaaaatattaacgcttaca
atttcaggtggcacttttcggggaaatgtgcgcggaacccctatttgtttatttttctaaatacattc

aaatatgtatccgctcatgagacaataaccctgataaatgcttcaataatattgaaaaaggaagagta
tgagtattcaacatttccgtgtcgcccttattcccttttttgcggcattttgccttcctgtttttgct
cacccagaaacgctggtgaaagtaaaagatgctgaagatcagttgggtgcacgagtgggttacatcga
actggatctcaacagcggtaagatccttgagagttttcgccccgaagaacgttttccaatgatgagca
cttttaaagttctgctatgtggcgcggtattatcccgtattgacgccgggcaagagcaactcggtcgc
cgcatacactattctcagaatgacttggttgagtactcaccagtcacagaaaagcatcttacggatgg
catgacagtaagagaattatgcagtgctgccataaccatgagtgataacactgcggccaacttacttc
tgacaacgatcggaggaccgaaggagctaaccgcttttttgcacaacatgggggatcatgtaactcgc
cttgatcgttgggaaccggagctgaatgaagccataccaaacgacgagcgtgacaccacgatgcctgt
agcaatggcaacaacgttgcgcaaactattaactggcgaactacttactctagcttcccggcaacaat
tgatagactggatggaggcggataaagttgcaggaccacttctgcgctcggcccttccggctggctgg
tttattgctgataaatctggagccggtgagcgtggctctcgcggtatcattgcagcactggggccaga
tggtaagccctcccgtatcgtagttatctacacgacggggagtcaggcaactatggatgaacgaaata
gacagatcgctgagataggtgcctcactgattaagcattggtaggaattaatgatgtctcgtttagat
aaaagtaaagtgattaacagcgcattagagctgcttaatgaggtcggaatcgaaggtttaacaacccg
taaactcgcccagaagctaggtgtagagcagcctacattgtattggcatgtaaaaaataagcgggctt
tgctcgacgccttagccattgagatgttagataggcaccatactcacttttgccctttagaaggggaa
agctggcaagattttttacgtaataacgctaaaagttttagatgtgctttactaagtcatcgcgatgg
agcaaaagtacatttaggtacacggcctacagaaaaacagtatgaaactctcgaaaatcaattagcct
ttttatgccaacaaggtttttcactagagaatgcattatatgcactcagcgcagtggggcattttact
ttaggttgcgtattggaagatcaagagcatcaagtcgctaaagaagaaagggaaacacctactactga
tagtatgccgccattattacgacaagctatcgaattatttgatcaccaaggtgcagagccagccttct
tattcggccttgaattgatcatatgcggattagaaaaacaacttaaatgtgaaagtgggtcttaaaag
cagcataacctttttccgtgatggtaacttcactagtttaaaaggatctaggtgaagatcctttttga
taatctcatgaccaaaatcccttaacgtgagttttcgttccactgagcgtcagaccccgtagaaaaga
tcaaaggatcttcttgagatcctttttttctgcgcgtaatctgctgcttgcaaacaaaaaaaccaccg
ctaccagcggtggtttgtttgccggatcaagagctaccaactctttttccgaaggtaactggcttcag
cagagcgcagataccaaatactgtccttctagtgtagccgtagttaggccaccacttcaagaactctg
tagcaccgcctacatacctcgctctgctaatcctgttaccagtggctgctgccagtggcgataagtcg
tgtcttaccgggttggactcaagacgatagttaccggataaggcgcagcggtcgggctgaacgggggg
ttcgtgcacacagcccagcttggagcgaacgacctacaccgaactgagatacctacagcgtgagctat
gagaaagcgccacgcttcccgaagggagaaaggcggacaggtatccggtaagcggcagggtcggaaca
ggagagcgcacgagggagcttccagggggaaacgcctggtatctttatagtcctgtcgggtttcgcca
cctctgacttgagcgtcgatttttgtgatgctcgtcaggggggcggagcctatggaaaaacgccagca
acgcggcctttttacggttcctggccttttgctggccttttgctcacatgacccgacaccatcgaatg
gccagat

>*Sif*DLH

MKTQSIEYASGPTRLVGHLAWDADIGARRPGVIVFPEAFGLNAHARERAERLARLGYVALAADLLGDG
RVFDNLPSVVPSIKALYADRTAWRARARAALEVLLARPEVDRERLGAIGFCFGGSTALELARSGAPLS
AVATFHAGLLPRLPEDAGRIGSRVLICHGDDDPVVNQDALATVVDELRRDRVDWQLARYGNTVHSFTD
PQADARNNPGFAYNALADRRSWAAMRQLFDEAFALP

>pASK-IBA5+_StrepII-*Sif*DLH

gattaattcctaattttgttgacactctatcattgatagagttattttaccactccctatcagtgat
agagaaaagtgaaatgaatagttcgacaaaaatctagaaataattttgtttaactttaagaaggagat
atacaaATGGCTAGCTGGAGCCACCCGCAGTTCGAAAAAGGCGCCAAAACCCAGAGCATTGAATATGC
AAGCGGTCCGACACGTCTGGTTGGTCATCTGGCATGGGATGCAGATATTGGTGCACGTCGTCCGGGTG
TTATTGTTTTTCCGGAAGCATTTGGTCTGAATGCACATGCACGTGAACGTGCAGAACGTCTGGCACGT
CTGGGTTATGTTGCACTGGCAGCCGATCTGCTTGGTGATGGTCGTGTTTTTGATAATCTGCCGAGCGT
TGTTCCGAGCATTAAAGCACTGTATGCAGATCGTACCGCATGGCGTGCACGTGCCCGTGCAGCACTGG
AAGTTCTGCTGGCTCGTCCGGAAGTTGATCGTGAACGCCTGGGTGCAATTGGTTTTTGTTTTGGTGGT
AGCACCGCACTGGAACTGGCACGTAGTGGTGCACCGCTGAGCGCAGTTGCAACCTTTCATGCAGGTCT
GCTGCCACGTCTGCCGGAAGATGCAGGTCGTATTGGTAGCCGTGTTCTGATTTGTCATGGTGATGATG
ATCCGGTTGTTAATCAGGATGCACTGGCCACCGTTGTTGATGAACTGCGTCGTGATCGTGTTGATTGG
CAGCTGGCACGTTATGGTAATACCGTTCATAGCTTTACCGATCCGCAGGCAGATGCACGTAATAATCC
GGGTTTTGCATATAATGCCCTGGCAGATCGTCGTAGCTGGGCAGCAATGCGTCAGCTGTTTGATGAAG
CCTTTGCACTGCCGTAAtgatatctaactaagcttgacctgtgaagtgaaaaatggcgcacattgtgc
gacatttttttgtctgccgtttaccgctactgcgtcacggatctccacgcgccctgtagcggcgcat
taagcgcggcgggtgtggtggttacgcgcagcgtgaccgctacacttgccagcgccctagcgcccgct
cctttcgctttcttcccttcctttctcgccacgttcgccggctttccccgtcaagctctaaatcgggg
gctccctttagggttccgatttagtgctttacggcacctcgaccccaaaaaacttgattagggtgatg
gttcacgtagtgggccatcgccctgatagacggtttttcgccctttgacgttggagtccacgttcttt
aatagtggactcttgttccaaactggaacaacactcaaccctatctcggtctattcttttgatttata
agggattttgccgatttcggcctattggttaaaaaatgagctgatttaacaaaaatttaacgcgaatt
ttaacaaaatattaacgcttacaatttcaggtggcacttttcggggaaatgtgcgcggaacccctatt
tgtttatttttctaaatacattcaaatatgtatccgctcatgagacaataaccctgataaatgcttca
ataatattgaaaaaggaagagtatgagtattcaacatttccgtgtcgcccttattcccttttttgcgg
cattttgccttcctgtttttgctcacccagaaacgctggtgaaagtaaaagatgctgaagatcagttg
ggtgcacgagtgggttacatcgaactggatctcaacagcggtaagatccttgagagttttcgccccga
agaacgttttccaatgatgagcacttttaaagttctgctatgtggcgcggtattatcccgtattgacg
ccgggcaagagcaactcggtcgccgcatacactattctcagaatgacttggttgagtactcaccagtc
acagaaaagcatcttacggatggcatgacagtaagagaattatgcagtgctgccataaccatgagtga
taacactgcggccaacttacttctgacaacgatcggaggaccgaaggagctaaccgcttttttgcaca

acatggggatcatgtaactcgccttgatcgttgggaaccggagctgaatgaagccataccaaacgac
gagcgtgacaccacgatgcctgtagcaatggcaacaacgttgcgcaaactattaactggcgaactact
tactctagcttcccggcaacaattgatagactggatggaggcggataaagttgcaggaccacttctgc
gctcggcccttccggctggctggtttattgctgataaatctggagccggtgagcgtggctctcgcggt
atcattgcagcactggggccagatggtaagccctcccgtatcgtagttatctacacgacggggagtca
ggcaactatggatgaacgaaatagacagatcgctgagataggtgcctcactgattaagcattggtagg
aattaatgatgtctcgtttagataaaagtaaagtgattaacagcgcattagagctgcttaatgaggtc
ggaatcgaaggtttaacaacccgtaaactcgcccagaagctaggtgtagagcagcctacattgtattg
gcatgtaaaaaataagcgggctttgctcgacgccttagccattgagatgttagataggcaccatactc
acttttgccctttagaaggggaaagctggcaagatttttacgtaataacgctaaaagttttagatgt
gctttactaagtcatcgcgatggagcaaaagtacatttaggtacacggcctacagaaaaacagtatga
aactctcgaaaatcaattagcctttttatgccaacaaggtttttcactagagaatgcattatatgcac
tcagcgcagtggggcattttactttaggttgcgtattggaagatcaagagcatcaagtcgctaaagaa
gaaagggaaacacctactactgatagtatgccgccattattacgacaagctatcgaattatttgatca
ccaaggtgcagagccagccttcttattcggccttgaattgatcatatgcggattagaaaaacaactta
aatgtgaaagtgggtcttaaaagcagcataacctttttccgtgatggtaacttcactagtttaaagg
atctaggtgaagatcctttttgataatctcatgaccaaaatcccttaacgtgagttttcgttccactg
agcgtcagaccccgtagaaaagatcaaaggatcttcttgagatcctttttttctgcgcgtaatctgct
gcttgcaaacaaaaaaaccaccgctaccagcggtggtttgtttgccggatcaagagctaccaactctt
tttccgaaggtaactggcttcagcagagcgcagataccaaatactgtccttctagtgtagccgtagtt
aggccaccacttcaagaactctgtagcaccgcctacatacctcgctctgctaatcctgttaccagtgg
ctgctgccagtggcgataagtcgtgtcttaccgggttggactcaagacgatagttaccggataaggcg
cagcggtcgggctgaacggggggttcgtgcacacagcccagcttggagcgaacgacctacaccgaact
gagatacctacagcgtgagctatgagaaagcgccacgcttcccgaagggagaaaggcggacaggtatc
cggtaagcggcagggtcggaacaggagagcgcacgagggagcttccagggggaaacgcctggtatctt
tatagtcctgtcgggtttcgccacctctgacttgagcgtcgatttttgtgatgctcgtcaggggggcg
gagcctatggaaaaacgccagcaacgcggcctttttacggttcctggccttttgctggccttttgctc
acatgacccgacaccatcgaatggccagat

>*m1*DLH

MKTETIEYSDGGTTCIGHLAWDDTQTGPRPGIVVFSEAYGLNDHARIRAERLAALGFVALAADLHGNG
LVYGDMASLGPAIQALYADRSAWRARALAAFNTLVALPQVDTNQTAAIGFCFGGATCFELARTGAPLG
GITVFHAGVIPELPEDKGRISGQVLICQGADDPVVKKEAVDAVTAELSRDKVDWQYIVYANTGHSFTD
PDADARNMPGFAYNALAEERSWMAMRLQYHEIFTVT

>pASK-IBA5+_StrepII-*m1*DLH

gattaattcctaattttgttgacactctatcattgatagagttattttaccactccctatcagtgat
agagaaaagtgaaatgaatagttcgacaaaaatctagaaataattttgtttaactttaagaaggagat
atacaaATGGCTAGCTGGAGCCACCCGCAGTTCGAAAAAGGCGCCAAAACCGAAACCATTGAATATAG
TGATGGTGGCACCACCTGTATTGGTCATCTGGCATGGGATGATACCCAGACCGGTCCGCGTCCGGGTA
TTGTTGTTTTTAGCGAAGCATATGGTCTGAATGATCATGCACGTATTCGTGCAGAACGTCTGGCAGCA
CTGGGTTTTGTTGCACTGGCAGCCGATCTGCATGGTAATGGTCTGGTTTATGGTGATATGGCAAGCCT
GGGTCCTGCAATTCAGGCACTGTATGCCGATCGTAGCGCATGGCGTGCACGTGCCCTGGCAGCATTTA
ATACCCTGGTTGCACTGCCGCAGGTTGATACCAATCAGACCGCAGCAATTGGTTTTTGTTTTGGTGGT
GCAACCTGTTTTGAACTGGCACGTACAGGTGCACCGTTAGGTGGTATTACCGTTTTTCATGCCGGTGT
TATTCCGGAACTGCCGGAAGATAAAGGTCGTATTAGCGGTCAGGTTCTGATTTGTCAGGGTGCAGATG
ATCCGGTTGTTAAAAAAGAAGCAGTTGACGCAGTTACCGCAGAACTGAGCCGTGATAAAGTTGATTGG
CAGTATATTGTGTATGCCAATACCGGTCATAGCTTTACCGATCCGGATGCAGATGCACGTAATATGCC
TGGTTTTGCATATAATGCACTGGCCGAAGAACGTAGCTGGATGGCAATGCGTCTGCAGTATCATGAAA
TCTTTACCGTGACCTAAtgatatctaactaagcttgacctgtgaagtgaaaaatggcgcacattgtgc
gacatttttttgtctgccgtttaccgctactgcgtcacggatctccacgcgccctgtagcggcgcat
taagcgcggcgggtgtggtggttacgcgcagcgtgaccgctacacttgccagcgccctagcgcccgct
cctttcgctttcttcccttcctttctcgccacgttcgccggctttccccgtcaagctctaaatcgggg
gctccctttagggttccgatttagtgctttacggcacctcgaccccaaaaaacttgattagggtgatg
gttcacgtagtgggccatcgccctgatagacggtttttcgccctttgacgttggagtccacgttcttt
aatagtggactcttgttccaaactggaacaacactcaaccctatctcggtctattcttttgatttata
agggattttgccgatttcggcctattggttaaaaaatgagctgatttaacaaaaatttaacgcgaatt
ttaacaaaatattaacgcttacaatttcaggtggcacttttcggggaaatgtgcgcggaacccctatt
tgtttatttttctaaatacattcaaatatgtatccgctcatgagacaataaccctgataaatgcttca
ataatattgaaaaaggaagagtatgagtattcaacatttccgtgtcgcccttattcccttttttgcgg
cattttgccttcctgtttttgctcacccagaaacgctggtgaaagtaaaagatgctgaagatcagttg
ggtgcacgagtgggttacatcgaactggatctcaacagcggtaagatccttgagagttttcgccccga
agaacgttttccaatgatgagcacttttaaagttctgctatgtggcgcggtattatcccgtattgacg
ccgggcaagagcaactcggtcgccgcatacactattctcagaatgacttggttgagtactcaccagtc
acagaaaagcatcttacggatggcatgacagtaagagaattatgcagtgctgccataaccatgagtga
taacactgcggccaacttacttctgacaacgatcggaggaccgaaggagctaaccgcttttttgcaca

acatgggggatcatgtaactcgccttgatcgttgggaaccggagctgaatgaagccataccaaacgac
gagcgtgacaccacgatgcctgtagcaatggcaacaacgttgcgcaaactattaactggcgaactact
tactctagcttcccggcaacaattgatagactggatggaggcggataaagttgcaggaccacttctgc
gctcggcccttccggctggctggtttattgctgataaatctggagccggtgagcgtggctctcgcggt
atcattgcagcactggggccagatggtaagccctcccgtatcgtagttatctacacgacggggagtca
ggcaactatggatgaacgaaatagacagatcgctgagataggtgcctcactgattaagcattggtagg
aattaatgatgtctcgtttagataaaagtaaagtgattaacagcgcattagagctgcttaatgaggtc
ggaatcgaaggtttaacaacccgtaaactcgcccagaagctaggtgtagagcagcctacattgtattg
gcatgtaaaaaataagcgggctttgctcgacgccttagccattgagatgttagataggcaccatactc
acttttgccctttagaaggggaaagctggcaagattttttacgtaataacgctaaaagttttagatgt
gctttactaagtcatcgcgatggagcaaaagtacatttaggtacacggcctacagaaaaacagtatga
aactctcgaaaatcaattagccttttatgccaacaaggtttttcactagagaatgcattatatgcac
tcagcgcagtggggcattttactttaggttgcgtattggaagatcaagagcatcaagtcgctaaagaa
gaaagggaaacacctactactgatagtatgccgccattattacgacaagctatcgaattatttgatca
ccaaggtgcagagccagccttcttattcggccttgaattgatcatatgcggattagaaaaacaactta
aatgtgaaagtgggtcttaaaagcagcataacctttttccgtgatggtaacttcactagtttaaagg
atctaggtgaagatcctttttgataatctcatgaccaaaatcccttaacgtgagttttcgttccactg
agcgtcagaccccgtagaaaagatcaaaggatcttcttgagatcctttttttctgcgcgtaatctgct
gcttgcaaacaaaaaaaccaccgctaccagcggtggtttgtttgccggatcaagagctaccaactctt
tttccgaaggtaactggcttcagcagagcgcagataccaaatactgtccttctagtgtagccgtagtt
aggccaccacttcaagaactctgtagcaccgcctacatacctcgctctgctaatcctgttaccagtgg
ctgctgccagtggcgataagtcgtgtcttaccgggttggactcaagacgatagttaccggataaggcg
cagcggtcgggctgaacggggggttcgtgcacacagcccagcttggagcgaacgacctacaccgaact
gagatacctacagcgtgagctatgagaaagcgccacgcttcccgaagggagaaaggcggacaggtatc
cggtaagcggcagggtcggaacaggagagcgcacgagggagcttccagggggaaacgcctggtatctt
tatagtcctgtcgggtttcgccacctctgacttgagcgtcgatttttgtgatgctcgtcaggggggcg
gagcctatggaaaaacgccagcaacgcggcctttttacggttcctggccttttgctggccttttgctc
acatgacccgacaccatcgaatggccagat

>*Sf*DLH

MHQQPIETTENGQRHIHQFFLDETLQGPRPGVLVFPEAFGLGDHALQRARRLAELGYAALAVDIHGEG
REFQDLAQVRPAILALFGDRAAWRARLQAAHELLRAQPQVDAARTAAIGFCFGGACSLELARSGAPLS
AIVTFHAGLQPPLEADAGKIKAKVLVCHGAEDPLMKPEPLAAILAELTRDKVDWQLLSHGNVVHSFTN
PDADARGAPGFAYNAGADRRSWAAMQGLFAEVFA

>pASK-IBA5+_StrepII-*Sf*DLH

gattaattcctaatttttgttgacactctatcattgatagagttattttaccactccctatcagtgat
agagaaaagtgaaatgaatagttcgacaaaaatctagaaataattttgtttaactttaagaaggagat
atacaaATGGCTAGCTGGAGCCACCCGCAGTTCGAAAAAGGCGCCATGCATCAGCAGCCGATTGAAAC
CACCGAAAATGGTCAGCGTCATATCCATCAGTTTTTCTGGATGAAACACTGCAGGGTCCGCGTCCGG
GTGTTCTGGTTTTTCCGGAAGCATTTGGTCTGGGTGATCATGCACTGCAGCGTGCACGTCGTCTGGCA
GAACTGGGTTATGCAGCACTGGCAGTTGATATTCATGGTGAAGGTCGTGAATTTCAGGATCTGGCACA
GGTTCGTCCGGCAATTCTGGCACTGTTTGGTGATCGTGCAGCATGGCGTGCCCGTCTGCAGGCAGCAC
ATGAACTGCTGCGTGCCCAGCCGCAGGTTGATGCAGCCCGTACCGCAGCAATTGGTTTTTGTTTTGGT
GGTGCATGTAGCCTGGAACTGGCACGTAGTGGTGCACCGCTGAGCGCAATTGTTACCTTTCATGCAGG
TCTGCAGCCTCCGCTGGAAGCAGATGCAGGTAAAATCAAAGCAAAAGTTCTGGTGTGTCATGGTGCCG
AAGATCCGCTGATGAAACCGGAACCGCTGGCAGCCATTCTGGCCGAACTGACCCGTGATAAAGTTGAT
TGGCAGCTGCTGAGCCATGGTAATGTTGTTCATAGCTTTACCAATCCGGATGCAGATGCACGTGGCGC
ACCGGGTTTTGCATATAATGCCGGTGCAGATCGTCGTAGCTGGGCAGCAATGCAGGGTCTGTTTGCCG
AAGTTTTTGCATAAtgatatctaactaagcttgacctgtgaagtgaaaaatggcgcacattgtgcgac
atttttttgtctgccgtttaccgctactgcgtcacggatctccacgcgccctgtagcggcgcattaa
gcgcggcgggtgtggtggttacgcgcagcgtgaccgctacacttgccagcgccctagcgcccgctcct
ttcgctttcttcccttcctttctcgccacgttcgccggctttccccgtcaagctctaaatcgggggct
ccctttagggttccgatttagtgctttacggcacctcgaccccaaaaaacttgattagggtgatggtt
cacgtagtgggccatcgccctgatagacggtttttcgccctttgacgttggagtccacgttctttaat
agtggactcttgttccaaactggaacaacactcaaccctatctcggtctattcttttgatttataagg
gattttgccgatttcggcctattggttaaaaaatgagctgatttaacaaaaatttaacgcgaatttta
acaaaatattaacgcttacaatttcaggtggcacttttcggggaaatgtgcgcggaacccctatttgt
ttatttttctaaatacattcaaatatgtatccgctcatgagacaataaccctgataaatgcttcaata
atattgaaaaaggaagagtatgagtattcaacatttccgtgtcgcccttattcccttttttgcggcat
tttgccttcctgtttttgctcacccagaaacgctggtgaaagtaaaagatgctgaagatcagttgggt
gcacgagtgggttacatcgaactggatctcaacagcggtaagatccttgagagttttcgccccgaaga
acgttttccaatgatgagcacttttaaagttctgctatgtggcgcggtattatcccgtattgacgccg
ggcaagagcaactcggtcgccgcatacactattctcagaatgacttggttgagtactcaccagtcaca
gaaaagcatcttacggatggcatgacagtaagagaattatgcagtgctgccataaccatgagtgataa
cactgcggccaacttacttctgacaacgatcggaggaccgaaggagctaaccgctttttttgcacaaca

tggggggatcatgtaactcgccttgatcgttgggaaccggagctgaatgaagccataccaaacgacgag
cgtgacaccacgatgcctgtagcaatggcaacaacgttgcgcaaactattaactggcgaactacttac
tctagcttcccggcaacaattgatagactggatggaggcggataaagttgcaggaccacttctgcgct
cggcccttccggctggctggtttattgctgataaatctggagccggtgagcgtggctctcgcggtatc
attgcagcactggggccagatggtaagccctcccgtatcgtagttatctacacgacggggagtcaggc
aactatggatgaacgaaatagacagatcgctgagataggtgcctcactgattaagcattggtaggaat
taatgatgtctcgtttagataaaagtaaagtgattaacagcgcattagagctgcttaatgaggtcgga
atcgaaggtttaacaacccgtaaactcgcccagaagctaggtgtagagcagcctacattgtattggca
tgtaaaaaataagcgggctttgctcgacgccttagccattgagatgttagataggcaccatactcact
tttgccctttagaaggggaaagctggcaagattttttacgtaataacgctaaaagttttagatgtgct
ttactaagtcatcgcgatggagcaaaagtacatttaggtacacggcctacagaaaaacagtatgaaac
tctcgaaaatcaattagccttttatgccaacaaggtttttcactagagaatgcattatatgcactca
gcgcagtggggcattttactttaggttgcgtattggaagatcaagagcatcaagtcgctaaagaagaa
agggaaacacctactactgatagtatgccgccattattacgacaagctatcgaattatttgatcacca
aggtgcagagccagccttcttattcggccttgaattgatcatatgcggattagaaaaacaacttaaat
gtgaaagtgggtcttaaaagcagcataacctttttccgtgatggtaacttcactagtttaaaggatc
taggtgaagatcctttttgataatctcatgaccaaaatcccttaacgtgagttttcgttccactgagc
gtcagaccccgtagaaaagatcaaaggatcttcttgagatcctttttttctgcgcgtaatctgctgct
tgcaaacaaaaaaaccaccgctaccagcggtggtttgtttgccggatcaagagctaccaactcttttt
ccgaaggtaactggcttcagcagagcgcagataccaaatactgtccttctagtgtagccgtagttagg
ccaccacttcaagaactctgtagcaccgcctacatacctcgctctgctaatcctgttaccagtggctg
ctgccagtggcgataagtcgtgtcttaccgggttggactcaagacgatagttaccggataaggcgcag
cggtcgggctgaacggggggttcgtgcacacagcccagcttggagcgaacgacctacaccgaactgag
atacctacagcgtgagctatgagaaagcgccacgcttcccgaagggagaaaggcggacaggtatccgg
taagcggcagggtcggaacaggagagcgcacgagggagcttccagggggaaacgcctggtatcttat
agtcctgtcgggtttcgccacctctgacttgagcgtcgatttttgtgatgctcgtcaggggggcggag
cctatggaaaaacgccagcaacgcggcctttttacggttcctggccttttgctggccttttgctcaca
tgacccgacaccatcgaatggccagat

>*Sc*DLH

MRKQKIEYGNGPTQFHGWLIRDDSLDGVRPGVLVFPEAYGLNEHAIERAERLAQLGYVALAADMHGGG
VVYSDTATLGPAIRSLFGDRAEWRARAQAALDALLAQPQVDRDRVAAIGFCFGGATCLELARSGAPLS
ALVTFHAGLQPPLEADAGRITGKVLICHGAEDPLMKPEALNAVLAELSRDRVDWQLLSFGGVAHSFTN
PDADARGAPGFAYNANADRRSWAAMQGLFAEVFAN

>pASK-IBA5+_StrepII-*Sc*DLH

gattaattcctaattttgttgacactctatcattgatagagttattttaccactccctatcagtgat
agagaaaagtgaaatgaatagttcgacaaaaatctagaaataattttgtttaactttaagaaggagat
atacaaATGGCTAGCTGGAGCCACCCGCAGTTCGAAAAAGGCGCCATGCGTAAACAGAAAATTGAATA
TGGTAATGGTCCGACGCAGTTTCATGGTTGGCTGATTCGTGATGATAGCCTGGATGGTGTTCGTCCGG
GTGTTCTGGTTTTTCCGGAAGCATATGGTCTGAATGAACATGCAATTGAACGTGCAGAACGTCTGGCA
CAGCTGGGTTATGTTGCACTGGCAGCAGATATGCATGGTGGTGGTGTTGTTTATAGCGATACCGCAAC
ACTGGGTCCTGCAATTCGTAGCCTGTTTGGTGATCGTGCAGAATGGCGTGCACGTGCCCAGGCAGCAC
TGGATGCACTGCTGGCCCAGCCGCAGGTTGATCGTGATCGTGTTGCAGCAATTGGTTTTTGTTTTGGT
GGTGCAACCTGTCTGGAACTGGCACGTAGTGGTGCACCGCTGAGCGCACTGGTTACCTTTCATGCAGG
TCTGCAGCCTCCGCTGGAAGCAGATGCAGGTCGTATTACCGGCAAAGTTCTGATTTGTCATGGTGCCG
AAGATCCGCTGATGAAACCGGAAGCACTGAATGCAGTTCTGGCGGAACTGAGCCGTGATCGCGTTGAT
TGGCAGCTGCTGAGCTTTGGTGGCGTTGCACATAGCTTTACCAATCCGGATGCAGATGCACGTGGCGC
ACCGGGTTTTGCATATAATGCCAATGCAGATCGTCGTAGCTGGGCAGCAATGCAGGGTCTGTTTGCCG
AAGTTTTTGCAAACTAAtgatatctaactaagcttgacctgtgaagtgaaaaatggcgcacattgtgc
gacatttttttgtctgccgtttaccgctactgcgtcacggatctccacgcgccctgtagcggcgcat
taagcgcggcgggtgtggtggttacgcgcagcgtgaccgctacacttgccagcgccctagcgcccgct
cctttcgctttcttcccttcctttctcgccacgttcgccggctttccccgtcaagctctaaatcgggg
gctccctttagggttccgatttagtgctttacggcacctcgaccccaaaaaacttgattagggtgatg
gttcacgtagtgggccatcgccctgatagacggtttttcgccctttgacgttggagtccacgttcttt
aatagtggactcttgttccaaactggaacaacactcaaccctatctcggtctattcttttgatttata
agggattttgccgatttcggcctattggttaaaaaatgagctgatttaacaaaaatttaacgcgaatt
ttaacaaaatattaacgcttacaatttcaggtggcacttttcggggaaatgtgcgcggaacccctatt
tgtttatttttctaaatacattcaaatatgtatccgctcatgagacaataaccctgataaatgcttca
ataatattgaaaaaggaagagtatgagtattcaacatttccgtgtcgcccttattcccttttttgcgg
cattttgccttcctgtttttgctcacccagaaacgctggtgaaagtaaaagatgctgaagatcagttg
ggtgcacgagtgggttacatcgaactggatctcaacagcggtaagatccttgagagttttcgccccga
agaacgttttccaatgatgagcacttttaaagttctgctatgtggcgcggtattatcccgtattgacg
ccgggcaagagcaactcggtcgccgcatacactattctcagaatgacttggttgagtactcaccagtc
acagaaaagcatcttacggatggcatgacagtaagagaattatgcagtgctgccataaccatgagtga
taacactgcggccaacttacttctgacaacgatcggaggaccgaaggagctaaccgcttttttgcaca

acatggggatcatgtaactcgccttgatcgttgggaaccggagctgaatgaagccataccaaacgac
gagcgtgacaccacgatgcctgtagcaatggcaacaacgttgcgcaaactattaactggcgaactact
tactctagcttcccggcaacaattgatagactggatggaggcggataaagttgcaggaccacttctgc
gctcggcccttccggctggctggtttattgctgataaatctggagccggtgagcgtggctctcgcggt
atcattgcagcactggggccagatggtaagccctcccgtatcgtagttatctacacgacggggagtca
ggcaactatggatgaacgaaatagacagatcgctgagataggtgcctcactgattaagcattggtagg
aattaatgatgtctcgtttagataaaagtaaagtgattaacagcgcattagagctgcttaatgaggtc
ggaatcgaaggtttaacaacccgtaaactcgcccagaagctaggtgtagagcagcctacattgtattg
gcatgtaaaaaataagcgggctttgctcgacgccttagccattgagatgttagataggcaccatactc
acttttgccctttagaaggggaaagctggcaagattttttacgtaataacgctaaaagttttagatgt
gctttactaagtcatcgcgatggagcaaaagtacatttaggtacacggcctacagaaaaacagtatga
aactctcgaaaatcaattagcctttttatgccaacaaggtttttcactagagaatgcattatatgcac
tcagcgcagtggggcattttactttaggttgcgtattggaagatcaagagcatcaagtcgctaaagaa
gaaagggaaacacctactactgatagtatgccgccattattacgacaagctatcgaattatttgatca
ccaaggtgcagagccagccttcttattcggccttgaattgatcatatgcggattagaaaaacaactta
aatgtgaaagtgggtcttaaaagcagcataacctttttccgtgatggtaacttcactagtttaaagg
atctaggtgaagatcctttttgataatctcatgaccaaaatcccttaacgtgagttttcgttccactg
agcgtcagaccccgtagaaaagatcaaaggatcttcttgagatcctttttttctgcgcgtaatctgct
gcttgcaaacaaaaaaaccaccgctaccagcggtggtttgtttgccggatcaagagctaccaactctt
tttccgaaggtaactggcttcagcagagcgcagataccaaatactgtccttctagtgtagccgtagtt
aggccaccacttcaagaactctgtagcaccgcctacatacctcgctctgctaatcctgttaccagtgg
ctgctgccagtggcgataagtcgtgtcttaccgggttggactcaagacgatagttaccggataaggcg
cagcggtcgggctgaacggggggttcgtgcacacagcccagcttggagcgaacgacctacaccgaact
gagatacctacagcgtgagctatgagaaagcgccacgcttcccgaagggagaaaggcggacaggtatc
cggtaagcggcagggtcggaacaggagagcgcacgagggagcttccagggggaaacgcctggtatctt
tatagtcctgtcgggtttcgccacctctgacttgagcgtcgatttttgtgatgctcgtcaggggggcg
gagcctatggaaaaacgccagcaacgcggcctttttacggttcctggccttttgctggccttttgctc
acatgacccgacaccatcgaatggccagat

>pASK-IBA5+_6xHis-*Sc*DLH

gattaattcctaatttttgttgacactctatcattgatagagttattttaccactccctatcagtgat
agagaaaagtgaaatgaatagttcgacaaaaatctagaaataattttgtttaactttaagaaggagat
atacaaATGCATCACCATCATCACCACGGTGGAATGCGTAAACAGAAAATTGAATATGGTAATGGTCC
GACGCAGTTTCATGGTTGGCTGATTCGTGATGATAGCCTGGATGGTGTTCGTCCGGGTGTTCTGGTTT
TTCCGGAAGCATATGGTCTGAATGAACATGCAATTGAACGTGCAGAACGTCTGGCACAGCTGGGTTAT
GTTGCACTGGCAGCAGATATGCATGGTGGTGGTGTTGTTTATAGCGATACCGCAACACTGGGTCCTGC
AATTCGTAGCCTGTTTGGTGATCGTGCAGAATGGCGTGCACGTGCCCAGGCAGCACTGGATGCACTGC

TGGCCCAGCCGCAGGTTGATCGTGATCGTGTTGCAGCAATTGGTTTTTGTTTTGGTGGTGCAACCTGT
CTGGAACTGGCACGTAGTGGTGCACCGCTGAGCGCACTGGTTACCTTTCATGCAGGTCTGCAGCCTCC
GCTGGAAGCAGATGCAGGTCGTATTACCGGCAAAGTTCTGATTTGTCATGGTGCCGAAGATCCGCTGA
TGAAACCGGAAGCACTGAATGCAGTTCTGGCGGAACTGAGCCGTGATCGCGTTGATTGGCAGCTGCTG
AGCTTTGGTGGCGTTGCACATAGCTTTACCAATCCGGATGCAGATGCACGTGGCGCACCGGGTTTTGC
ATATAATGCCAATGCAGATCGTCGTAGCTGGGCAGCAATGCAGGGTCTGTTTGCCGAAGTTTTTGCAA
ACTAAtgatatctaactaagcttgacctgtgaagtgaaaaatggcgcacattgtgcgacattttttt
gtctgccgtttaccgctactgcgtcacggatctccacgcgccctgtagcggcgcattaagcgcggcgg
gtgtggtggttacgcgcagcgtgaccgctacacttgccagcgccctagcgcccgctcctttcgctttc
ttcccttcctttctcgccacgttcgccggctttccccgtcaagctctaaatcgggggctccctttagg
gttccgatttagtgctttacggcacctcgaccccaaaaaacttgattagggtgatggttcacgtagtg
ggccatcgccctgatagacggtttttcgccctttgacgttggagtccacgttctttaatagtggactc
ttgttccaaactggaacaacactcaaccctatctcggtctattcttttgatttataagggattttgcc
gatttcggcctattggttaaaaaatgagctgatttaacaaaaatttaacgcgaattttaacaaaatat
taacgcttacaatttcaggtggcacttttcggggaaatgtgcgcggaacccctatttgtttatttttc
taaatacattcaaatatgtatccgctcatgagacaataaccctgataaatgcttcaataatattgaaa
aaggaagagtatgagtattcaacatttccgtgtcgcccttattcccttttttgcggcattttgccttc
ctgtttttgctcacccagaaacgctggtgaaagtaaaagatgctgaagatcagttgggtgcacgagtg
ggttacatcgaactggatctcaacagcggtaagatccttgagagttttcgccccgaagaacgttttcc
aatgatgagcacttttaaagttctgctatgtggcgcggtattatcccgtattgacgccgggcaagagc
aactcggtcgccgcatacactattctcagaatgacttggttgagtactcaccagtcacagaaaagcat
cttacggatggcatgacagtaagagaattatgcagtgctgccataaccatgagtgataacactgcggc
caacttacttctgacaacgatcggaggaccgaaggagctaaccgcttttttgcacaacatgggggatc
atgtaactcgccttgatcgttgggaaccggagctgaatgaagccataccaaacgacgagcgtgacacc
acgatgcctgtagcaatggcaacaacgttgcgcaaactattaactggcgaactacttactctagcttc
ccggcaacaattgatagactggatggaggcggataaagttgcaggaccacttctgcgctcggcccttc
cggctggctggtttattgctgataaatctggagccggtgagcgtggctctcgcggtatcattgcagca
ctggggccagatggtaagccctcccgtatcgtagttatctacacgacggggagtcaggcaactatgga
tgaacgaaatagacagatcgctgagataggtgcctcactgattaagcattggtaggaattaatgatgt
ctcgtttagataaaagtaaagtgattaacagcgcattagagctgcttaatgaggtcggaatcgaaggt
ttaacaacccgtaaactcgcccagaagctaggtgtagagcagcctacattgtattggcatgtaaaaaa
taagcgggctttgctcgacgccttagccattgagatgttagataggcaccatactcacttttgccctt
tagaaggggaaagctggcaagattttttacgtaataacgctaaaagttttagatgtgctttactaagt
catcgcgatggagcaaaagtacatttaggtacacggcctacagaaaaacagtatgaaactctcgaaaa
tcaattagcctttttatgccaacaaggtttttcactagagaatgcattatatgcactcagcgcagtgg
ggcattttactttaggttgcgtattggaagatcaagagcatcaagtcgctaaagaagaaagggaaaca
cctactactgatagtatgccgccattattacgacaagctatcgaattatttgatcaccaaggtgcaga

gccagccttcttattcggccttgaattgatcatatgcggattagaaaaacaacttaaatgtgaaagtg
ggtcttaaaagcagcataacctttttccgtgatggtaacttcactagtttaaaaggatctaggtgaag
atcctttttgataatctcatgaccaaaatcccttaacgtgagttttcgttccactgagcgtcagaccc
cgtagaaaagatcaaaggatcttcttgagatcctttttttctgcgcgtaatctgctgcttgcaaacaa
aaaaaccaccgctaccagcggtggtttgtttgccggatcaagagctaccaactctttttccgaaggta
actggcttcagcagagcgcagataccaaatactgtccttctagtgtagccgtagttaggccaccactt
caagaactctgtagcaccgcctacatacctcgctctgctaatcctgttaccagtggctgctgccagtg
gcgataagtcgtgtcttaccgggttggactcaagacgatagttaccggataaggcgcagcggtcgggc
tgaacggggggttcgtgcacacagcccagcttggagcgaacgacctacaccgaactgagatacctaca
gcgtgagctatgagaaagcgccacgcttcccgaagggagaaaggcggacaggtatccggtaagcggca
gggtcggaacaggagagcgcacgagggagcttccagggggaaacgcctggtatctttatagtcctgtc
gggtttcgccacctctgacttgagcgtcgatttttgtgatgctcgtcaggggggcggagcctatggaa
aaacgccagcaacgcggcctttttacggttcctggccttttgctggccttttgctcacatgacccgac
accatcgaatggccagat

>*Sa*DLH

MKSQQVVYRGAGRRFLGELYWDEAATQPAPGVLVFPDAFGLADHARERAQRLAQLGYVALAADLHGEG
AVYEDVASMRVHLQPLFENRADWRARAQAALAALQAQTPVDAQRLAAIGFCLGGATCLELARCGAPLK
AIVGFHAGVLAPLPGDEQKIQAKVLLCQGADDPLIKKENMAAVEAELRRDRVDWQLIVYGNAVHSFTN
RDAATRQSPAMAYDAAADRRSWAAMQGLLAEVF

>pASK-IBA5+_StrepII-*Sa*DLH

gattaattcctaattttgttgacactctatcattgatagagttattttaccactccctatcagtgat
agagaaaagtgaaatgaatagttcgacaaaaatctagaaataattttgtttaactttaagaaggagat
atacaaATGGCTAGCTGGAGCCACCCGCAGTTCGAAAAAGGCGCCAAAAGCCAGCAGGTTGTTTATCG
TGGTGCAGGTCGTCGTTTTCTGGGTGAACTGTATTGGGATGAAGCAGCAACCCAGCCTGCACCGGGTG
TTCTGGTTTTTCCGGATGCATTTGGTCTGGCAGATCATGCACGTGAACGTGCACAGCGTCTGGCACAG
CTGGGTTATGTTGCACTGGCAGCCGATCTGCATGGTGAAGGTGCAGTTTATGAAGATGTTGCAAGCAT
GCGTGTTCATCTGCAGCCGCTGTTTGAAAATCGTGCAGATTGGCGTGCACGTGCCCAGGCAGCCCTGG
CAGCACTGCAGGCACAGACACCGGTTGATGCCCAACGTCTGGCAGCAATTGGTTTTTGTTTAGGTGGT
GCAACCTGTCTGGAACTGGCACGTTGTGGTGCACCGCTGAAAGCAATTGTTGGTTTTCATGCCGGTGT
GCTGGCACCGCTGCCTGGTGATGAACAGAAAATTCAGGCAAAAGTTCTGCTGTGTCAGGGTGCAGATG
ATCCGCTGATCAAAAAAGAAAATATGGCAGCAGTTGAAGCAGAACTGCGTCGTGATCGTGTTGATTGG
CAGCTGATTGTTTATGGTAATGCCGTTCATAGCTTTACCAATCGTGATGCAGCGACCCGTCAGAGTCC
GGCAATGGCATATGATGCAGCAGCAGATCGTCGTAGCTGGGCAGCCATGCAGGGTCTGCTGGCAGAAG
TTTTCTAAtgatatctaactaagcttgacctgtgaagtgaaaaatggcgcacattgtgcgacattttt
tttgtctgccgtttaccgctactgcgtcacggatctccacgcgccctgtagcggcgcattaagcgcgg
cgggtgtggtggttacgcgcagcgtgaccgctacacttgccagcgccctagcgcccgctcctttcgct
ttcttcccttcctttctcgccacgttcgccggctttccccgtcaagctctaaatcgggggctcccttt
agggttccgatttagtgctttacggcacctcgaccccaaaaaacttgattagggtgatggttcacgta
gtgggccatcgccctgatagacggtttttcgccctttgacgttggagtccacgttctttaatagtgga
ctcttgttccaaactggaacaacactcaaccctatctcggtctattcttttgatttataagggatttt
gccgatttcggcctattggttaaaaaatgagctgatttaacaaaaatttaacgcgaattttaacaaaa
tattaacgcttacaatttcaggtggcacttttcggggaaatgtgcgcggaacccctatttgtttattt
ttctaaatacattcaaatatgtatccgctcatgagacaataaccctgataaatgcttcaataatattg
aaaaaggaagagtatgagtattcaacatttccgtgtcgcccttattcccttttttgcggcattttgcc
ttcctgtttttgctcacccagaaacgctggtgaaagtaaaagatgctgaagatcagttgggtgcacga
gtgggttacatcgaactggatctcaacagcggtaagatccttgagagttttcgccccgaagaacgttt
tccaatgatgagcacttttaaagttctgctatgtggcgcggtattatcccgtattgacgccgggcaag
agcaactcggtcgccgcatacactattctcagaatgacttggttgagtactcaccagtcacagaaaag
catcttacggatggcatgacagtaagagaattatgcagtgctgccataaccatgagtgataacactgc
ggccaacttacttctgacaacgatcggaggaccgaaggagctaaccgcttttttgcacaacatgggg

atcatgtaactcgccttgatcgttgggaaccggagctgaatgaagccataccaaacgacgagcgtgac
accacgatgcctgtagcaatggcaacaacgttgcgcaaactattaactggcgaactacttactctagc
ttcccggcaacaattgatagactggatggaggcggataaagttgcaggaccacttctgcgctcggccc
ttccggctggctggtttattgctgataaatctggagccggtgagcgtggctctcgcggtatcattgca
gcactggggccagatggtaagccctcccgtatcgtagttatctacacgacggggagtcaggcaactat
ggatgaacgaaatagacagatcgctgagataggtgcctcactgattaagcattggtaggaattaatga
tgtctcgtttagataaaagtaaagtgattaacagcgcattagagctgcttaatgaggtcggaatcgaa
ggtttaacaacccgtaaactcgcccagaagctaggtgtagagcagcctacattgtattggcatgtaaa
aaataagcgggctttgctcgacgccttagccattgagatgttagataggcaccatactcacttttgcc
ctttagaaggggaaagctggcaagatttttacgtaataacgctaaaagttttagatgtgctttacta
agtcatcgcgatggagcaaaagtacatttaggtacacggcctacagaaaaacagtatgaaactctcga
aaatcaattagcctttttatgccaacaaggtttttcactagagaatgcattatatgcactcagcgcag
tggggcattttactttaggttgcgtattggaagatcaagagcatcaagtcgctaaagaagaaaggaa
acacctactactgatagtatgccgccattattacgacaagctatcgaattatttgatcaccaaggtgc
agagccagccttcttattcggccttgaattgatcatatgcggattagaaaaacaacttaaatgtgaaa
gtgggtcttaaaagcagcataacctttttccgtgatggtaacttcactagtttaaaaggatctaggtg
aagatcctttttgataatctcatgaccaaaatcccttaacgtgagttttcgttccactgagcgtcaga
ccccgtagaaaagatcaaaggatcttcttgagatcctttttttctgcgcgtaatctgctgcttgcaaa
caaaaaaaccaccgctaccagcggtggtttgtttgccggatcaagagctaccaactctttttccgaag
gtaactggcttcagcagagcgcagataccaaatactgtccttctagtgtagccgtagttaggccacca
cttcaagaactctgtagcaccgcctacatacctcgctctgctaatcctgttaccagtggctgctgcca
gtggcgataagtcgtgtcttaccgggttggactcaagacgatagttaccggataaggcgcagcggtcg
ggctgaacggggggttcgtgcacacagcccagcttggagcgaacgacctacaccgaactgagatacct
acagcgtgagctatgagaaagcgccacgcttcccgaagggagaaaggcggacaggtatccggtaagcg
gcagggtcggaacaggagagcgcacgagggagcttccagggggaaacgcctggtatctttatagtcct
gtcgggtttcgccacctctgacttgagcgtcgatttttgtgatgctcgtcaggggggcggagcctatg
gaaaaacgccagcaacgcggcctttttacggttcctggccttttgctggccttttgctcacatgaccc
gacaccatcgaatggccagat

>*Mz*DLH

MKPHALLHPLIVLAIAPALVQAELHTEEINYRVGDDQFTGYLAYDDQISGQRPGILIVHEWWGHNEFA
RQQAERLATEGFTAFALDMYGSGKVADHPDNARQFMQAATENSDVIRERFEAAMRLLQDQPTVDASKI
AAQGYCFGGAVVLNMARMGMDLAGVVSIHGSLASPIQAEPGRVKARVQVYTGGADQMVPADQVAALVH
EMQSAGVDLTLTSYPGVKHSFSNPDADVVAERFGMPVAYDEQAAERTWRGTLAFYQELFGR

>pASK-IBA5+_StrepII-*Mz*DLH

gattaattcctaattttgttgacactctatcattgatagagttattttaccactccctatcagtgat
agagaaaagtgaaatgaatagttcgacaaaaatctagaaataattttgtttaactttaagaaggagat
atacaaATGGCTAGCTGGAGCCACCCGCAGTTCGAAAAAGGCGCCATGAAACCGCATGCACTGCTGCA
TCCGCTGATTGTTCTGGCAATTGCACCGGCACTGGTTCAGGCAGAACTGCATACCGAAGAAATTAACT
ATCGTGTTGGTGATGATCAGTTTACCGGTTATCTGGCATATGATGATCAAATTAGCGGTCAGCGTCCG
GGTATTCTGATTGTGCATGAATGGTGGGGTCATAATGAATTTGCACGTCAGCAGGCCGAACGTCTGGC
AACCGAAGGTTTTACCGCATTTGCACTGGATATGTATGGTAGCGGTAAAGTTGCAGATCATCCGGATA
ATGCCCGTCAGTTTATGCAGGCAGCAACCGAAAATAGTGATGTTATTCGTGAACGTTTTGAAGCAGCA
ATGCGTCTGCTGCAGGATCAGCCGACCGTTGATGCAAGCAAAATTGCAGCACAGGGTTATTGTTTTGG
TGGTGCAGTTGTTCTGAATATGGCACGTATGGGTATGGATCTGGCAGGCGTTGTTAGCATTCATGGTA
GCCTGGCAAGCCCGATTCAGGCGGAACCGGGTCGTGTTAAAGCACGTGTTCAGGTTTATACCGGTGGT
GCCGATCAGATGGTTCCGGCAGATCAGGTTGCAGCCCTGGTTCATGAAATGCAGTCAGCCGGTGTTGA
TCTGACCCTGACCAGCTATCCGGGTGTTAAACATAGCTTTAGCAATCCGGATGCAGATGTTGTTGCAG
AACGCTTTGGTATGCCGGTTGCCTATGATGAACAGGCAGCAGAACGTACCTGGCGTGGCACCCTGGCA
TTCTATCAAGAACTGTTTGGTCGTTAAtgatatctaactaagcttgacctgtgaagtgaaaaatggcg
cacattgtgcgacatttttttgtctgccgtttaccgctactgcgtcacggatctccacgcgccctgt
agcggcgcattaagcgcggcgggtgtggtggttacgcgcagcgtgaccgctacacttgccagcgccct
agcgcccgctcctttcgctttcttcccttcctttctcgccacgttcgccggctttccccgtcaagctc
taaatcgggggctccctttagggttccgatttagtgctttacggcacctcgaccccaaaaaacttgat
tagggtgatggttcacgtagtgggccatcgccctgatagacggtttttcgccctttgacgttggagtc
cacgttctttaatagtggactcttgttccaaactggaacaacactcaaccctatctcggtctattctt
ttgatttataagggattttgccgatttcggcctattggttaaaaaatgagctgatttaacaaaaattt
aacgcgaattttaacaaaatattaacgcttacaatttcaggtggcacttttcggggaaatgtgcgcgg
aacccctatttgtttatttttctaaatacattcaaatatgtatccgctcatgagacaataaccctgat
aaatgcttcaataatattgaaaaaggaagagtatgagtattcaacatttccgtgtcgcccttattccc
ttttttgcggcattttgccttcctgtttttgctcacccagaaacgctggtgaaagtaaaagatgctga
agatcagttgggtgcacgagtgggttacatcgaactggatctcaacagcggtaagatccttgagagtt
ttcgccccgaagaacgttttccaatgatgagcacttttaaagttctgctatgtggcgcggtattatcc
cgtattgacgccgggcaagagcaactcggtcgccgcatacactattctcagaatgacttggttgagta
ctcaccagtcacagaaaagcatcttacggatggcatgacagtaagagaattatgcagtgctgccataa

ccatgagtgataacactgcggccaacttacttctgacaacgatcggaggaccgaaggagctaaccgct
tttttgcacaacatgggggatcatgtaactcgccttgatcgttgggaaccggagctgaatgaagccat
accaaacgacgagcgtgacaccacgatgcctgtagcaatggcaacaacgttgcgcaaactattaactg
gcgaactacttactctagcttcccggcaacaattgatagactggatggaggcggataaagttgcagga
ccacttctgcgctcggcccttccggctggctggtttattgctgataaatctggagccggtgagcgtgg
ctctcgcggtatcattgcagcactggggccagatggtaagccctcccgtatcgtagttatctacacga
cggggagtcaggcaactatggatgaacgaaatagacagatcgctgagataggtgcctcactgattaag
cattggtaggaattaatgatgtctcgtttagataaaagtaaagtgattaacagcgcattagagctgct
taatgaggtcggaatcgaaggtttaacaacccgtaaactcgcccagaagctaggtgtagagcagccta
cattgtattggcatgtaaaaaataagcgggctttgctcgacgccttagccattgagatgttagatagg
caccatactcacttttgccctttagaaggggaaagctggcaagattttttacgtaataacgctaaaag
ttttagatgtgctttactaagtcatcgcgatggagcaaaagtacatttaggtacacggcctacagaaa
aacagtatgaaactctcgaaaatcaattagcctttttatgccaacaaggtttttcactagagaatgca
ttatatgcactcagcgcagtggggcattttactttaggttgcgtattggaagatcaagagcatcaagt
cgctaaagaagaaagggaaacacctactactgatagtatgccgccattattacgacaagctatcgaat
tatttgatcaccaaggtgcagagccagccttcttattcggccttgaattgatcatatgcggattagaa
aaacaacttaaatgtgaaagtgggtcttaaaagcagcataacctttttccgtgatggtaacttcacta
gtttaaaaggatctaggtgaagatcctttttgataatctcatgaccaaaatcccttaacgtgagtttt
cgttccactgagcgtcagaccccgtagaaaagatcaaaggatcttcttgagatcctttttttctgcgc
gtaatctgctgcttgcaaacaaaaaaaccaccgctaccagcggtggtttgtttgccggatcaagagct
accaactctttttccgaaggtaactggcttcagcagagcgcagataccaaatactgtccttctagtgt
agccgtagttaggccaccacttcaagaactctgtagcaccgcctacatacctcgctctgctaatcctg
ttaccagtggctgctgccagtggcgataagtcgtgtcttaccgggttggactcaagacgatagttacc
ggataaggcgcagcggtcgggctgaacggggggttcgtgcacacagcccagcttggagcgaacgacct
acaccgaactgagatacctacagcgtgagctatgagaaagcgccacgcttcccgaagggagaaaggcg
gacaggtatccggtaagcggcagggtcggaacaggagagcgcacgagggagcttccagggggaaacgc
ctggtatctttatagtcctgtcgggtttcgccacctctgacttgagcgtcgatttttgtgatgctcgt
caggggggcggagcctatggaaaaacgccagcaacgcggcctttttacggttcctggccttttgctgg
ccttttgctcacatgacccgacaccatcgaatggccagat

>*Hs*DLH (CMBL)

MANEAYPCPCDIGHRLEYGGLGREVQVEHIKAYVTKSPVDAGKAVIVIQDIFGWQLPNTRYIADMISG
NGYTTIVPDFFVGQEPWDPSGDWSIFPEWLKTRNAQKIDREISAILKYLKQQCHAQKIGIVGFCWGGT
AVHHLMMKYSEFRAGVSVYGIVKDSEDIYNLKNPTLFIFAENDVVIPLKDVSLLTQKLKEHCKVEYQI
KTFSGQTHGFVHRKREDCSPADKPYIDEARRNLIEWLNKYM

>pASK-IBA5+_StrepII-*Hs*DLH

gattaattcctaattttgttgacactctatcattgatagagttattttaccactccctatcagtgat
agagaaaagtgaaatgaatagttcgacaaaaatctagaaataattttgtttaactttaagaaggagat
atacaaATGGCTAGCTGGAGCCACCCGCAGTTCGAAAAAGGCGCCATGGCAAATGAAGCATATCCGTG
TCCGTGTGATATTGGTCATCGTCTGGAATATGGTGGTCTGGGTCGTGAAGTTCAGGTTGAACATATTA
AAGCCTACGTTACCAAAAGTCCGGTTGATGCAGGTAAAGCCGTTATTGTTATTCAGGATATTTTTGGT
TGGCAACTGCCGAATACACGTTATATTGCAGATATGATTAGCGGCAATGGCTATACCACCATTGTTCC
GGATTTTTTTGTTGGTCAAGAACCGTGGGATCCGAGCGGTGATTGGAGCATTTTTCCGGAATGGCTGA
AAACCCGTAATGCCCAGAAAATTGATCGTGAAATTAGCGCCATTCTGAAGTATCTGAAACAGCAGTGT
CATGCACAGAAAATCGGTATTGTTGGTTTTTGCTGGGGTGGCACCGCAGTTCATCACCTGATGATGAA
ATATTCAGAATTTCGTGCCGGTGTTAGCGTGTATGGTATTGTTAAAGATAGCGAGGATATCTATAACC
TGAAAAATCCGACGCTGTTTATCTTTGCCGAAAACGATGTTGTTATCCCGCTGAAAGATGTTAGCCTG
CTGACCCAGAAACTGAAAGAACATTGCAAAGTGGAATACCAGATCAAAACCTTTAGCGGTCAGACCCA
TGGTTTTGTTCATCGTAAACGTGAAGATTGTAGTCCGGCAGATAAACCGTATATTGATGAAGCACGTC
GCAATCTGATTGAGTGGCTGAACAAATATATGTAAtgatatctaactaagcttgacctgtgaagtgaa
aaatggcgcacattgtgcgacatttttttgtctgccgtttaccgctactgcgtcacggatctccacg
cgccctgtagcggcgcattaagcgcggcgggtgtggtggttacgcgcagcgtgaccgctacacttgcc
agcgccctagcgcccgctcctttcgctttcttcccttcctttctcgccacgttcgccggctttccccg
tcaagctctaaatcgggggctccctttagggttccgatttagtgctttacggcacctcgaccccaaaa
aacttgattagggtgatggttcacgtagtgggccatcgccctgatagacggtttttcgccctttgacg
ttggagtccacgttctttaatagtggactcttgttccaaactggaacaacactcaaccctatctcggt
ctattcttttgatttataagggattttgccgatttcggcctattggttaaaaaatgagctgatttaac
aaaaatttaacgcgaattttaacaaaatattaacgcttacaatttcaggtggcacttttcggggaaat
gtgcgcggaacccctatttgtttatttttctaaatacattcaaatatgtatccgctcatgagacaata
accctgataaatgcttcaataatattgaaaaaggaagagtatgagtattcaacatttccgtgtcgccc
ttattcccttttttgcggcattttgccttcctgtttttgctcacccagaaacgctggtgaaagtaaaa
gatgctgaagatcagttgggtgcacgagtgggttacatcgaactggatctcaacagcggtaagatcct
tgagagttttcgccccgaagaacgttttccaatgatgagcacttttaaagttctgctatgtggcgcgg
tattatcccgtattgacgccgggcaagagcaactcggtcgccgcatacactattctcagaatgacttg
gttgagtactcaccagtcacagaaaagcatcttacggatggcatgacagtaagagaattatgcagtgc
tgccataaccatgagtgataacactgcggccaacttacttctgacaacgatcggaggaccgaaggagc

taaccgctttttttgcacaacatgggggatcatgtaactcgccttgatcgttgggaaccggagctgaat
gaagccataccaaacgacgagcgtgacaccacgatgcctgtagcaatggcaacaacgttgcgcaaact
attaactggcgaactacttactctagcttcccggcaacaattgatagactggatggaggcggataaag
ttgcaggaccacttctgcgctcggcccttccggctggctggtttattgctgataaatctggagccggt
gagcgtggctctcgcggtatcattgcagcactggggccagatggtaagccctcccgtatcgtagttat
ctacacgacggggagtcaggcaactatggatgaacgaaatagacagatcgctgagataggtgcctcac
tgattaagcattggtaggaattaatgatgtctcgtttagataaaagtaaagtgattaacagcgcatta
gagctgcttaatgaggtcggaatcgaaggtttaacaacccgtaaactcgcccagaagctaggtgtaga
gcagcctacattgtattggcatgtaaaaaataagcgggctttgctcgacgccttagccattgagatgt
tagataggcaccatactcacttttgccctttagaaggggaaagctggcaagattttttacgtaataac
gctaaaagttttagatgtgctttactaagtcatcgcgatggagcaaaagtacatttaggtacacggcc
tacagaaaaacagtatgaaactctcgaaaatcaattagcctttttatgccaacaaggtttttcactag
agaatgcattatatgcactcagcgcagtggggcattttactttaggttgcgtattggaagatcaagag
catcaagtcgctaaagaagaaagggaaacacctactactgatagtatgccgccattattacgacaagc
tatcgaattatttgatcaccaaggtgcagagccagccttcttattcggccttgaattgatcatatgcg
gattagaaaaacaacttaaatgtgaaagtgggtcttaaaagcagcataacctttttccgtgatggtaa
cttcactagtttaaaaggatctaggtgaagatcctttttgataatctcatgaccaaaatcccttaacg
tgagttttcgttccactgagcgtcagaccccgtagaaaagatcaaaggatcttcttgagatcctttt
ttctgcgcgtaatctgctgcttgcaaacaaaaaaaccaccgctaccagcggtggtttgtttgccggat
caagagctaccaactctttttccgaaggtaactggcttcagcagagcgcagataccaaatactgtcct
tctagtgtagccgtagttaggccaccacttcaagaactctgtagcaccgcctacatacctcgctctgc
taatcctgttaccagtggctgctgccagtggcgataagtcgtgtcttaccgggttggactcaagacga
tagttaccggataaggcgcagcggtcgggctgaacggggggttcgtgcacacagcccagcttggagcg
aacgacctacaccgaactgagatacctacagcgtgagctatgagaaagcgccacgcttcccgaaggga
gaaaggcggacaggtatccggtaagcggcagggtcggaacaggagagcgcacgagggagcttccaggg
ggaaacgcctggtatctttatagtcctgtcgggtttcgccacctctgacttgagcgtcgatttttgtg
atgctcgtcaggggggcggagcctatggaaaaacgccagcaacgcggcctttttacggttcctggcct
tttgctggccttttgctcacatgacccgacaccatcgaatggccagat

>pASK-IBA5+_6xHis-TwinStrep-SUMO-*Hs*DLH
gattaattcctaatttttgttgacactctatcattgatagagttattttaccactccctatcagtgat
agagaaaagtgaaatgaatagttcgacaaaaatctagaaataattttgtttaactttaagaaggagat
atacaaATGGGCAGCAGCCATCATCATCATCATCACAGCAGCGGCCTGGTGCCGCGCGGCAGCCATAT
GGCTAGCTGGAGCCATCCGCAGTTTGAAAAAGGTGGTGGTAGCGGTGGTGGTTCAGGTGGTAGTGCAT
GGTCACACCCTCAGTTTGAGAAAATGTCGGACTCAGAAGTCAATCAAGAAGCTAAGCCAGAGGTCAAG
CCAGAAGTCAAGCCTGAGACTCACATCAATTTAAAGGTGTCCGATGGATCTTCAGAGATCTTCTTCaA
GATCAAAAAGAccACTCCTTTaAGAaggCTGATggAAGCGTTCGCTAAAAGACAGGGTAAGGAAATGG

ACTCCTTAAGATTCTTGTACGACGGTATTAGAATCCAAGCTGATCAGACCCCTGAAGATTTGGACATG
GAGGATAACGATATTATTGAGGCTCACAGAGAACAGATTGGTGGATCCATGGCAAATGAAGCATATCC
GTGTCCGTGTGATATTGGTCATCGTCTGGAATATGGTGGTCTGGGTCGTGAAGTTCAGGTTGAACATA
TTAAAGCCTACGTTACCAAAAGTCCGGTTGATGCAGGTAAAGCCGTTATTGTTATTCAGGATATTTTT
GGTTGGCAACTGCCGAATACACGTTATATTGCAGATATGATTAGCGGCAATGGCTATACCACCATTGT
TCCGGATTTTTTTGTTGGTCAAGAACCGTGGGATCCGAGCGGTGATTGGAGCATTTTTCCGGAATGGC
TGAAAACCCGTAATGCCCAGAAAATTGATCGTGAAATTAGCGCCATTCTGAAGTATCTGAAACAGCAG
TGTCATGCACAGAAAATCGGTATTGTTGGTTTTTGCTGGGGTGGCACCGCAGTTCATCACCTGATGAT
GAAATATTCAGAATTTCGTGCCGGTGTTAGCGTGTATGGTATTGTTAAAGATAGCGAGGATATCTATA
ACCTGAAAAATCCGACGCTGTTTATCTTTGCCGAAAACGATGTTGTTATCCCGCTGAAAGATGTTAGC
CTGCTGACCCAGAAACTGAAAGAACATTGCAAAGTGGAATACCAGATCAAAACCTTTAGCGGTCAGAC
CCATGGTTTTGTTCATCGTAAACGTGAAGATTGTAGTCCGGCAGATAAACCGTATATTGATGAAGCAC
GTCGCAATCTGATTGAGTGGCTGAACAAATATATGTAAtgatatctaactaagcttgacctgtgaagt
gaaaaatggcgcacattgtgcgacatttttttgtctgccgtttaccgctactgcgtcacggatctcc
acgcgccctgtagcggcgcattaagcgcggcgggtgtggtggttacgcgcagcgtgaccgctacactt
gccagcgccctagcgcccgctcctttcgctttcttcccttcctttctcgccacgttcgccggctttcc
ccgtcaagctctaaatcgggggctccctttagggttccgatttagtgctttacggcacctcgacccca
aaaaacttgattagggtgatggttcacgtagtgggccatcgccctgatagacggtttttcgccctttg
acgttggagtccacgttctttaatagtggactcttgttccaaactggaacaacactcaaccctatctc
ggtctattcttttgatttataagggattttgccgatttcggcctattggttaaaaaatgagctgattt
aacaaaaatttaacgcgaattttaacaaaatattaacgcttacaatttcaggtggcacttttcgggga
aatgtgcgcggaacccctatttgtttatttttctaaatacattcaaatatgtatccgctcatgagaca
ataaccctgataaatgcttcaataatattgaaaaaggaagagtatgagtattcaacatttccgtgtcg
cccttattcccttttttgcggcattttgccttcctgtttttgctcacccagaaacgctggtgaaagta
aaagatgctgaagatcagttgggtgcacgagtgggttacatcgaactggatctcaacagcggtaagat
ccttgagagttttcgccccgaagaacgttttccaatgatgagcacttttaaagttctgctatgtggcg
cggtattatcccgtattgacgccgggcaagagcaactcggtcgccgcatacactattctcagaatgac
ttggttgagtactcaccagtcacagaaaagcatcttacggatggcatgacagtaagagaattatgcag
tgctgccataaccatgagtgataacactgcggccaacttacttctgacaacgatcggaggaccgaagg
agctaaccgcttttttgcacaacatgggggatcatgtaactcgccttgatcgttgggaaccggagctg
aatgaagccataccaaacgacgagcgtgacaccacgatgcctgtagcaatggcaacaacgttgcgcaa
actattaactggcgaactacttactctagcttcccggcaacaattgatagactggatggaggcggata
aagttgcaggaccacttctgcgctcggcccttccggctggctggtttattgctgataaatctggagcc
ggtgagcgtggctctcgcggtatcattgcagcactggggccagatggtaagccctcccgtatcgtagt
tatctacacgacggggagtcaggcaactatggatgaacgaaatagacagatcgctgagataggtgcct
cactgattaagcattggtaggaattaatgatgtctcgtttagataaaagtaaagtgattaacagcgca
ttagagctgcttaatgaggtcggaatcgaaggtttaacaacccgtaaactcgcccagaagctaggtgt

agagcagcctacattgtattggcatgtaaaaaataagcgggctttgctcgacgccttagccattgaga
tgttagataggcaccatactcacttttgccctttagaaggggaaagctggcaagattttttacgtaat
aacgctaaaagttttagatgtgctttactaagtcatcgcgatggagcaaaagtacatttaggtacacg
gcctacagaaaaacagtatgaaactctcgaaaatcaattagcctttttatgccaacaaggtttttcac
tagagaatgcattatatgcactcagcgcagtggggcattttactttaggttgcgtattggaagatcaa
gagcatcaagtcgctaaagaagaaagggaaacacctactactgatagtatgccgccattattacgaca
agctatcgaattatttgatcaccaaggtgcagagccagccttcttattcggccttgaattgatcatat
gcggattagaaaaacaacttaaatgtgaaagtgggtcttaaaagcagcataacctttttccgtgatgg
taacttcactagtttaaaaggatctaggtgaagatcctttttgataatctcatgaccaaaatccctta
acgtgagttttcgttccactgagcgtcagaccccgtagaaaagatcaaaggatcttcttgagatcctt
tttttctgcgcgtaatctgctgcttgcaaacaaaaaaaccaccgctaccagcggtggtttgtttgccg
gatcaagagctaccaactctttttccgaaggtaactggcttcagcagagcgcagataccaaatactgt
ccttctagtgtagccgtagttaggccaccacttcaagaactctgtagcaccgcctacatacctcgctc
tgctaatcctgttaccagtggctgctgccagtggcgataagtcgtgtcttaccgggttggactcaaga
cgatagttaccggataaggcgcagcggtcgggctgaacggggggttcgtgcacacagcccagcttgga
gcgaacgacctacaccgaactgagatacctacagcgtgagctatgagaaagcgccacgcttcccgaag
ggagaaaggcggacaggtatccggtaagcggcagggtcggaacaggagagcgcacgagggagcttcca
gggggaaacgcctggtatctttatagtcctgtcgggtttcgccacctctgacttgagcgtcgattttt
gtgatgctcgtcaggggggcggagcctatggaaaaacgccagcaacgcggcctttttacggttcctgg
ccttttgctggccttttgctcacatgacccgacaccatcgaatggccagat

>*Hs*DLH-1E11

MANEAYPCPCDIGHRLEYGGLGREVQVEHIKAYVTKSPVDAGKAVIVIPEAFGWQLPNTRYIADMISG
NGYTTIVPDFFVGQEPWDPSGDWSIFPEWLKTRNAQKIDREISAILKYLKQQCHAQKIGIVGFCWGGT
AVHHLMMKYSEFRAGVSVYGIVKDSEDIYNLKNPTLFIFAENDVVIPLKDVSLLTQKLKEHCKVEYQI
KTFGNAVHSFTDPLAGSHGWPGVAYDATYIDEARRNLIEWLNKYM

>pASK-IBA5+_6xHis-TwinStrep-SUMO-*Hs*DLH-1E11

gattaattcctaatttttgttgacactctatcattgatagagttattttaccactccctatcagtgat
agagaaaagtgaaatgaatagttcgacaaaaatctagaaataattttgtttaactttaagaaggagat
atacaaATGGGCAGCAGCCATCATCATCATCATCACAGCAGCGGCCTGGTGCCGCGCGGCAGCCATAT
GGCTAGCTGGAGCCATCCGCAGTTTGAAAAAGGTGGTGGTAGCGGTGGTGGTTCAGGTGGTAGTGCAT
GGTCACACCCTCAGTTTGAGAAAATGTCGGACTCAGAAGTCAATCAAGAAGCTAAGCCAGAGGTCAAG
CCAGAAGTCAAGCCTGAGACTCACATCAATTTAAAGGTGTCCGATGGATCTTCAGAGATCTTCTTCaA
GATCAAAAAGAccACTCCTTTaAGAaggCTGATggAAGCGTTCGCTAAAAGACAGGGTAAGGAAATGG
ACTCCTTAAGATTCTTGTACGACGGTATTAGAATCCAAGCTGATCAGACCCCTGAAGATTTGGACATG
GAGGATAACGATATTATTGAGGCTCACAGAGAACAGATTGGTGGATCCATGGCAAATGAAGCATATCC

GTGTCCGTGTGATATTGGTCATCGTCTGGAATATGGTGGTCTGGGTCGTGAAGTTCAGGTTGAACATA
TTAAAGCCTACGTTACCAAAAGTCCGGTTGATGCAGGTAAAGCCGTTATTGTTATTCCGGAAGCGTTT
GGCTGGCAGCTTCCAAACACCCGTTATATTGCAGATATGATTAGCGGCAATGGCTATACCACCATTGT
TCCGGATTTTTTTGTTGGTCAAGAACCGTGGGATCCGAGCGGTGATTGGAGCATTTTTCCGGAATGGC
TGAAAACCCGTAATGCCCAGAAAATTGATCGTGAAATTAGCGCCATTCTGAAGTATCTGAAACAGCAG
TGTCATGCACAGAAAATCGGTATTGTTGGTTTTTGCTGGGGTGGCACCGCAGTTCATCACCTGATGAT
GAAATATTCAGAATTTCGTGCCGGTGTTAGCGTGTACGGCATCGTGAAAGATAGCGAGGATATCTATA
ACCTGAAAAATCCGACGCTGTTTATCTTTGCCGAAAACGATGTTGTTATCCCGCTGAAAGATGTTAGC
CTGCTGACCCAGAAACTGAAAGAACATTGCAAAGTGGAATACCAGATCAAAACCTTTGGAAATGCGGT
ACACAGTTTCACCGATCCACTCGCTGGCAGTCACGGCTGGCCCGGGGTTGCCTATGACGCCACTTACA
TTGATGAAGCACGTCGCAATCTGATTGAGTGGCTGAACAAATATATGTAAtgatatctaactaagctt
gacctgtgaagtgaaaaatggcgcacattgtgcgacatttttttgtctgccgtttaccgctactgcg
tcacggatctccacgcgccctgtagcggcgcattaagcgcggcgggtgtggtggttacgcgcagcgtg
accgctacacttgccagcgccctagcgcccgctcctttcgctttcttcccttcctttctcgccacgtt
cgccggctttccccgtcaagctctaaatcgggggctccctttagggttccgatttagtgctttacggc
acctcgaccccaaaaaacttgattagggtgatggttcacgtagtgggccatcgccctgatagacggtt
tttcgccctttgacgttggagtccacgttctttaatagtggactcttgttccaaactggaacaacact
caaccctatctcggtctattcttttgatttataagggattttgccgatttcggcctattggttaaaaa
atgagctgatttaacaaaaatttaacgcgaattttaacaaaatattaacgcttacaatttcaggtggc
acttttcggggaaatgtgcgcggaacccctatttgtttatttttctaaatacattcaaatatgtatcc
gctcatgagacaataaccctgataaatgcttcaataatattgaaaaaggaagagtatgagtattcaac
atttccgtgtcgcccttattcccttttttgcggcattttgccttcctgtttttgctcacccagaaacg
ctggtgaaagtaaaagatgctgaagatcagttgggtgcacgagtgggttacatcgaactggatctcaa
cagcggtaagatccttgagagttttcgccccgaagaacgttttccaatgatgagcacttttaaagttc
tgctatgtggcgcggtattatcccgtattgacgccgggcaagagcaactcggtcgccgcatacactat
tctcagaatgacttggttgagtactcaccagtcacagaaaagcatcttacggatggcatgacagtaag
agaattatgcagtgctgccataaccatgagtgataacactgcggccaacttacttctgacaacgatcg
gaggaccgaaggagctaaccgcttttttgcacaacatgggggatcatgtaactcgccttgatcgttgg
gaaccggagctgaatgaagccataccaaacgacgagcgtgacaccacgatgcctgtagcaatggcaac
aacgttgcgcaaactattaactggcgaactacttactctagcttcccggcaacaattgatagactgga
tggaggcggataaagttgcaggaccacttctgcgctcggcccttccggctggctggtttattgctgat
aaatctggagccggtgagcgtggctctcgcggtatcattgcagcactggggccagatggtaagccctc
ccgtatcgtagttatctacacgacggggagtcaggcaactatggatgaacgaaatagacagatcgctg
agataggtgcctcactgattaagcattggtaggaattaatgatgtctcgtttagataaaagtaaagtg
attaacagcgcattagagctgcttaatgaggtcggaatcgaaggtttaacaacccgtaaactcgccca
gaagctaggtgtagagcagcctacattgtattggcatgtaaaaaataagcgggctttgctcgacgcct
tagccattgagatgttagataggcaccatactcacttttgccctttagaaggggaaagctggcaagat

tttttacgtaataacgctaaaagttttagatgtgctttactaagtcatcgcgatggagcaaaagtaca
tttaggtacacggcctacagaaaaacagtatgaaactctcgaaaatcaattagccttttatgccaac
aaggtttttcactagagaatgcattatatgcactcagcgcagtggggcattttactttaggttgcgta
ttggaagatcaagagcatcaagtcgctaaagaagaaagggaaacacctactactgatagtatgccgcc
attattacgacaagctatcgaattatttgatcaccaaggtgcagagccagccttcttattcggccttg
aattgatcatatgcggattagaaaaacaacttaaatgtgaaagtgggtcttaaaagcagcataacctt
tttccgtgatggtaacttcactagtttaaaaggatctaggtgaagatcctttttgataatctcatgac
caaaatcccttaacgtgagttttcgttccactgagcgtcagaccccgtagaaaagatcaaaggatctt
cttgagatcctttttttctgcgcgtaatctgctgcttgcaaacaaaaaaaccaccgctaccagcggtg
gtttgtttgccggatcaagagctaccaactctttttccgaaggtaactggcttcagcagagcgcagat
accaaatactgtccttctagtgtagccgtagttaggccaccacttcaagaactctgtagcaccgccta
catacctcgctctgctaatcctgttaccagtggctgctgccagtggcgataagtcgtgtcttaccggg
ttggactcaagacgatagttaccggataaggcgcagcggtcgggctgaacggggggttcgtgcacaca
gcccagcttggagcgaacgacctacaccgaactgagatacctacagcgtgagctatgagaaagcgcca
cgcttcccgaagggagaaaggcggacaggtatccggtaagcggcagggtcggaacaggagagcgcacg
agggagcttccagggggaaacgcctggtatctttatagtcctgtcgggtttcgccacctctgacttga
gcgtcgatttttgtgatgctcgtcaggggggcggagcctatggaaaaacgccagcaacgcggcctttt
tacggttcctggccttttgctggccttttgctcacatgacccgacaccatcgaatggccagat

>*Kp*DLH

MTTTKQPGFAPAASPHAATAVHTPEEHIIAGETSIPSQGENMPAYHARPKNADGPLPIVIVVQEIFGV
HEHIRDLCRRLAQEGYLAIAPELYFRQGDPNEYHDIPTLFKELVSKVPDAQVLADLDHVASWAARHGG
DAHRLLITGFCWGGRITWLYAAHNPQLKAAVAWYGKLVGEKSLNSPKHPVDIAVDLNAPVLGLYGAKD
ASIPQDTVETMRQALRAANATAEIVVYPEADHAFNADYRASYHEESAKDGWQRMLAWFAQYGGKKG

>pASK-IBA5+_StrepII-*Kp*DLH

gattaattcctaattttttgttgacactctatcattgatagagttattttaccactccctatcagtgat
agagaaaagtgaaatgaatagttcgacaaaaatctagaaataattttgtttaactttaagaaggagat
atacaaATGGCTAGCTGGAGCCACCCGCAGTTCGAAAAAGGCGCCATGACCACCACCAAACAGCCTGG
TTTTGCACCGGCAGCAAGTCCGCATGCAGCAACCGCAGTTCATACACCGGAAGAACATATTATTGCCG
GTGAAACCAGCATTCCGAGCCAGGGTGAAAATATGCCTGCATATCATGCACGTCCGAAAAATGCAGAT
GGTCCGCTGCCGATTGTTATTGTTGTTCAAGAAATTTTTGGCGTGCACGAACATATTCGTGATCTGTG
TCGTCGTCTGGCACAAGAAGGTTATCTGGCAATTGCACCGGAACTGTATTTTCGTCAGGGTGATCCGA
ATGAATATCACGATATTCCGACGCTGTTTAAAGAACTGGTTAGCAAAGTTCCGGATGCACAGGTTCTG
GCAGATCTGGATCATGTTGCAAGCTGGGCAGCACGTCATGGTGGTGATGCACATCGTCTGCTGATTAC
CGGTTTTTGTTGGGGTGGTCGTATTACCTGGCTGTATGCAGCACATAATCCGCAGCTGAAAGCAGCAG
TTGCATGGTATGGTAAACTGGTTGGTGAAAAAAGCCTGAATAGCCCGAAACATCCGGTTGATATTGCA
GTTGATCTGAACGCACCGGTTCTGGGTCTGTATGGTGCAAAAGATGCAAGCATTCCGCAGGATACCGT
TGAAACCATGCGTCAGGCACTGCGTGCAGCAAATGCCACCGCAGAAATTGTTGTTTATCCGGAAGCAG
ATCATGCCTTTAATGCAGATTATCGTGCAAGCTATCATGAAGAAAGCGCCAAAGATGGTTGGCAGCGT
ATGCTGGCATGGTTTGCACAGTATGGTGGTAAAAAAGGCTAAtgatatctaactaagcttgacctgtg
aagtgaaaaatggcgcacattgtgcgacattttttttgtctgccgtttaccgctactgcgtcacggat
ctccacgcgccctgtagcggcgcattaagcgcggcgggtgtggtggttacgcgcagcgtgaccgctac
acttgccagcgccctagcgcccgctcctttcgctttcttcccttcctttctcgccacgttcgccggct
ttccccgtcaagctctaaatcgggggctccctttagggttccgatttagtgctttacggcacctcgac
cccaaaaaacttgattagggtgatggttcacgtagtgggccatcgccctgatagacggtttttcgccc
tttgacgttggagtccacgttctttaatagtggactcttgttccaaactggaacaacactcaacccta
tctcggtctattcttttgatttataagggattttgccgatttcggcctattggttaaaaaatgagctg
atttaacaaaaatttaacgcgaattttaacaaaatattaacgcttacaatttcaggtggcacttttcg
gggaaatgtgcgcggaacccctatttgtttatttttctaaatacattcaaatatgtatccgctcatga
gacaataaccctgataaatgcttcaataatattgaaaaaggaagagtatgagtattcaacatttccgt
gtcgcccttattcccttttttgcggcattttgccttcctgtttttgctcacccagaaacgctggtgaa
agtaaaagatgctgaagatcagttgggtgcacgagtgggttacatcgaactggatctcaacagcggta
agatccttgagagttttcgccccgaagaacgttttccaatgatgagcacttttaaagttctgctatgt
ggcgcggtattatcccgtattgacgccgggcaagagcaactcggtcgccgcatacactattctcagaa
tgacttggttgagtactcaccagtcacagaaaagcatcttacggatggcatgacagtaagagaattat

gcagtgctgccataaccatgagtgataacactgcggccaacttacttctgacaacgatcggaggaccg
aaggagctaaccgcttttttgcacaacatgggggatcatgtaactcgccttgatcgttgggaaccgga
gctgaatgaagccataccaaacgacgagcgtgacaccacgatgcctgtagcaatggcaacaacgttgc
gcaaactattaactggcgaactacttactctagcttcccggcaacaattgatagactggatggaggcg
gataaagttgcaggaccacttctgcgctcggcccttccggctggctggtttattgctgataaatctgg
agccggtgagcgtggctctcgcggtatcattgcagcactggggccagatggtaagccctcccgtatcg
tagttatctacacgacggggagtcaggcaactatggatgaacgaaatagacagatcgctgagataggt
gcctcactgattaagcattggtaggaattaatgatgtctcgtttagataaaagtaaagtgattaacag
cgcattagagctgcttaatgaggtcggaatcgaaggtttaacaacccgtaaactcgcccagaagctag
gtgtagagcagcctacattgtattggcatgtaaaaaataagcgggctttgctcgacgccttagccatt
gagatgttagataggcaccatactcacttttgccctttagaaggggaaagctggcaagattttttacg
taataacgctaaaagttttagatgtgctttactaagtcatcgcgatggagcaaaagtacatttaggta
cacggcctacagaaaaacagtatgaaactctcgaaaatcaattagccttttatgccaacaaggtttt
tcactagagaatgcattatatgcactcagcgcagtggggcattttactttaggttgcgtattggaaga
tcaagagcatcaagtcgctaaagaagaaagggaaacacctactactgatagtatgccgccattattac
gacaagctatcgaattatttgatcaccaaggtgcagagccagccttcttattcggccttgaattgatc
atatgcggattagaaaaacaacttaaatgtgaaagtgggtcttaaaagcagcataacctttttccgtg
atggtaacttcactagtttaaaaggatctaggtgaagatcctttttgataatctcatgaccaaaatcc
cttaacgtgagttttcgttccactgagcgtcagaccccgtagaaaagatcaaaggatcttcttgagat
cctttttttctgcgcgtaatctgctgcttgcaaacaaaaaaaccaccgctaccagcggtggtttgttt
gccggatcaagagctaccaactctttttccgaaggtaactggcttcagcagagcgcagataccaaata
ctgtccttctagtgtagccgtagttaggccaccacttcaagaactctgtagcaccgcctacatacctc
gctctgctaatcctgttaccagtggctgctgccagtggcgataagtcgtgtcttaccgggttggactc
aagacgatagttaccggataaggcgcagcggtcgggctgaacggggggttcgtgcacacagcccagct
tggagcgaacgacctacaccgaactgagatacctacagcgtgagctatgagaaagcgccacgcttccc
gaagggagaaaggcggacaggtatccggtaagcggcagggtcggaacaggagagcgcacgagggagct
tccagggggaaacgcctggtatctttatagtcctgtcgggtttcgccacctctgacttgagcgtcgat
ttttgtgatgctcgtcaggggggcggagcctatggaaaaacgccagcaacgcggcctttttacggttc
ctggccttttgctggccttttgctcacatgacccgacaccatcgaatggccagat

>*Eco*DLH

MLCLKKHQLRSATMPRLTAKDFPQELLDYYDYYAHGKISKREFLNLAAKYAVGGMTALALFDLLKPNY
ALATQVEFTDPEIFAEYITYPSPNGHGEVRGYLVKPAKMSGKTPAVVVVHENRGLNPYIEDVARRVAK
AGYIALAPDGLNSVGGYPGNDDKGRELQQQVDPTKLMNDFFAAIEFMQRYPQATGKVGITGFCYGGGV
SNAAAVAYPELACAVPFYGRQAPTADVAKIEAPLLLHFAELDTRINEGWPAYEAALKANNKVYEAYIY
PGVNHGFHNDSTPRYDKSAADLAWQRTLKWFDKYLS

>pASK-IBA5+_StrepII-*Eco*DLH

gattaattcctaatttttgttgacactctatcattgatagagttattttaccactccctatcagtgat
agagaaaagtgaaatgaatagttcgacaaaaatctagaaataattttgtttaactttaagaaggagat
atacaaATGGCTAGCTGGAGCCACCCGCAGTTCGAAAAAGGCGCCATGCTGTGCCTGAAAAAACATCA
GCTGCGTAGCGCAACCATGCCTCGTCTGACCGCAAAAGATTTTCCGCAAGAACTGCTGGATTATTATG
ATTATTACGCCCATGGCAAAATCAGCAAACGCGAATTTCTGAATCTGGCAGCAAAATATGCAGTTGGT
GGTATGACCGCACTGGCACTGTTTGATCTGCTGAAACCGAATTATGCACTGGCAACCCAGGTTGAATT
TACCGATCCGGAAATTTTTGCCGAGTATATCACCTATCCGAGTCCGAATGGTCATGGTGAAGTTCGTG
GTTATCTGGTTAAACCTGCAAAAATGAGCGGTAAAACACCGGCAGTTGTTGTTGTTCATGAAAATCGT
GGTCTGAACCCGTATATTGAAGATGTTGCACGTCGTGTTGCAAAAGCAGGTTATATTGCCCTGGCACC
GGATGGTCTGAATAGCGTTGGTGGTTATCCGGGTAATGATGATAAAGGTCGTGAACTGCAGCAGCAGG
TTGATCCGACCAAACTGATGAATGATTTTTTTGCAGCCATCGAATTCATGCAGCGTTATCCGCAGGCA
ACCGGTAAAGTTGGTATTACCGGTTTTTGTTATGGTGGTGGTGTTAGCAATGCAGCAGCCGTTGCATA
TCCGGAACTGGCATGTGCAGTTCCGTTTTATGGTCGTCAGGCACCGACCGCAGATGTTGCCAAAATTG
AAGCACCGCTGCTGTTACATTTTGCAGAACTGGATACCCGTATTAATGAAGGTTGGCCTGCATATGAA
GCAGCACTGAAAGCAAACAACAAAGTGTATGAGGCATATATCTATCCGGGTGTGAATCATGGCTTTCA
TAATGATAGCACACCGCGTTATGATAAAAGCGCAGCAGATCTGGCATGGCAGCGTACCCTGAAATGGT
TTGATAAATATCTGAGCTAAtgatatctaactaagcttgacctgtgaagtgaaaaatggcgcacattg
tgcgacatttttttgtctgccgtttaccgctactgcgtcacggatctccacgcgccctgtagcggcg
cattaagcgcggcgggtgtggtggttacgcgcagcgtgaccgctacacttgccagcgccctagcgccc
gctcctttcgctttcttcccttcctttctcgccacgttcgccggctttccccgtcaagctctaaatcg
ggggctccctttagggttccgatttagtgctttacggcacctcgaccccaaaaaacttgattagggtg
atggttcacgtagtgggccatcgccctgatagacggtttttcgccctttgacgttggagtccacgttc
tttaatagtggactcttgttccaaactggaacaacactcaaccctatctcggtctattcttttgattt
ataagggattttgccgatttcggcctattggttaaaaaatgagctgatttaacaaaaatttaacgcga
attttaacaaaatattaacgcttacaatttcaggtggcacttttcggggaaatgtgcgcggaacccct
atttgtttatttttctaaatacattcaaatatgtatccgctcatgagacaataaccctgataaatgct
tcaataatattgaaaaaggaagagtatgagtattcaacatttccgtgtcgcccttattcccttttttg
cggcattttgccttcctgtttttgctcacccagaaacgctggtgaaagtaaaagatgctgaagatcag
ttgggtgcacgagtgggttacatcgaactggatctcaacagcggtaagatccttgagagttttcgccc

cgaagaacgttttccaatgatgagcacttttaaagttctgctatgtggcgcggtattatcccgtattg
acgccgggcaagagcaactcggtcgccgcatacactattctcagaatgacttggttgagtactcacca
gtcacagaaaagcatcttacggatggcatgacagtaagagaattatgcagtgctgccataaccatgag
tgataacactgcggccaacttacttctgacaacgatcggaggaccgaaggagctaaccgcttttttgc
acaacatgggggatcatgtaactcgccttgatcgttgggaaccggagctgaatgaagccataccaaac
gacgagcgtgacaccacgatgcctgtagcaatggcaacaacgttgcgcaaactattaactggcgaact
acttactctagcttcccggcaacaattgatagactggatggaggcggataaagttgcaggaccacttc
tgcgctcggcccttccggctggctggtttattgctgataaatctggagccggtgagcgtggctctcgc
ggtatcattgcagcactggggccagatggtaagccctcccgtatcgtagttatctacacgacggggag
tcaggcaactatggatgaacgaaatagacagatcgctgagataggtgcctcactgattaagcattggt
aggaattaatgatgtctcgtttagataaaagtaaagtgattaacagcgcattagagctgcttaatgag
gtcggaatcgaaggtttaacaacccgtaaactcgcccagaagctaggtgtagagcagcctacattgta
ttggcatgtaaaaaataagcgggctttgctcgacgccttagccattgagatgttagataggcaccata
ctcacttttgccctttagaaggggaaagctggcaagattttttacgtaataacgctaaaagttttaga
tgtgctttactaagtcatcgcgatggagcaaaagtacatttaggtacacggcctacagaaaaacagta
tgaaactctcgaaaatcaattagcctttttatgccaacaaggtttttcactagagaatgcattatatg
cactcagcgcagtggggcattttactttaggttgcgtattggaagatcaagagcatcaagtcgctaaa
gaagaaagggaaacacctactactgatagtatgccgccattattacgacaagctatcgaattatttga
tcaccaaggtgcagagccagccttcttattcggccttgaattgatcatatgcggattagaaaaacaac
ttaaatgtgaaagtgggtcttaaaagcagcataacctttttccgtgatggtaacttcactagtttaaa
aggatctaggtgaagatcctttttgataatctcatgaccaaaatcccttaacgtgagttttcgttcca
ctgagcgtcagaccccgtagaaaagatcaaaggatcttcttgagatcctttttttctgcgcgtaatct
gctgcttgcaaacaaaaaaaccaccgctaccagcggtggtttgtttgccggatcaagagctaccaact
ctttttccgaaggtaactggcttcagcagagcgcagataccaaatactgtccttctagtgtagccgta
gttaggccaccacttcaagaactctgtagcaccgcctacatacctcgctctgctaatcctgttaccag
tggctgctgccagtggcgataagtcgtgtcttaccgggttggactcaagacgatagttaccggataag
gcgcagcggtcgggctgaacggggggttcgtgcacacagcccagcttggagcgaacgacctacaccga
actgagatacctacagcgtgagctatgagaaagcgccacgcttcccgaagggagaaaggcggacaggt
atccggtaagcggcagggtcggaacaggagagcgcacgagggagcttccagggggaaacgcctggtat
ctttatagtcctgtcgggtttcgccacctctgacttgagcgtcgatttttgtgatgctcgtcagggg
gcggagcctatggaaaaacgccagcaacgcggcctttttacggttcctggccttttgctggcctttt
ctcacatgacccgacaccatcgaatggccagat

>*Pk*DLH

MLTEGISIQSYDGHTFGALVGSPAKAPAPVIVIAQEIFGVNAFMRETVSWLVDQGYAAVCPDLYARQA
PGTALDPQDERQREQAYKLWQAFDMEAGVGDLEAAIRYARHQPYSNGKVGLVGYCLGGALAFLVAAKG
YVDRAVGYYGVGLEKQLNKVPEVKHPALFHMGGQDHFVPAPSRQLITEGFGANPLLQVHWYEEAGHSF
ARTSSSGYVASAAALANERTLDFLAPLQSKKP

>pASK-IBA5+_StrepII-*Pk*DLH

gattaattcctaatttttgttgacactctatcattgatagagttattttaccactccctatcagtgat
agagaaaagtgaaatgaatagttcgacaaaaatctagaaataattttgtttaactttaagaaggagat
atacaaATGGCTAGCTGGAGCCACCCGCAGTTCGAAAAAGGCGCCATGCTGACCGAAGGTATTAGCAT
TCAGAGCTATGATGGTCATACCTTTGGTGCACTGGTTGGTAGTCCGGCAAAAGCACCGGCACCGGTTA
TTGTTATTGCACAAGAAATTTTTGGCGTGAACGCCTTTATGCGTGAAACCGTTAGCTGGCTGGTTGAT
CAGGGTTATGCAGCAGTTTGTCCGGATCTGTATGCACGTCAGGCACCGGGTACAGCACTGGATCCGCA
GGATGAACGTCAGCGTGAACAGGCATATAAACTGTGGCAGGCATTTGATATGGAAGCCGGTGTTGGTG
ATCTGGAAGCAGCAATTCGTTATGCCCGTCATCAGCCGTATAGCAATGGTAAAGTTGGTCTGGTTGGT
TATTGCTTAGGTGGTGCCCTGGCATTTCTGGTTGCAGCAAAAGGTTATGTTGATCGTGCAGTGGGTTA
TTATGGTGTTGGCCTGGAAAAACAGCTGAATAAAGTTCCGGAAGTTAAACATCCGGCACTGTTTCATA
TGGGTGGTCAGGATCATTTTGTTCCGGCACCGAGCCGTCAGCTGATTACCGAAGGTTTTGGTGCAAAT
CCGCTGCTGCAGGTTCATTGGTATGAAGAAGCAGGTCATAGCTTTGCACGTACCAGCAGCAGTGGTTA
TGTGGCAAGCGCAGCAGCACTGGCAAATGAACGTACCCTGGATTTTCTGGCACCGCTGCAGAGCAAAA
AACCGTAAtgatatctaactaagcttgacctgtgaagtgaaaaatggcgcacattgtgcgacattttt
tttgtctgccgtttaccgctactgcgtcacggatctccacgcgccctgtagcggcgcattaagcgcgg
cgggtgtggtggttacgcgcagcgtgaccgctacacttgccagcgccctagcgcccgctcctttcgct
ttcttcccttcctttctcgccacgttcgccggctttccccgtcaagctctaaatcgggggctcccttt
agggttccgatttagtgctttacggcacctcgaccccaaaaaacttgattagggtgatggttcacgta
gtgggccatcgccctgatagacggtttttcgccctttgacgttggagtccacgttctttaatagtgga
ctcttgttccaaactggaacaacactcaaccctatctcggtctattcttttgatttataagggatttt
gccgatttcggcctattggttaaaaaatgagctgatttaacaaaaatttaacgcgaattttaacaaaa
tattaacgcttacaatttcaggtggcacttttcggggaaatgtgcgcggaacccctatttgtttattt
ttctaaatacattcaaatatgtatccgctcatgagacaataaccctgataaatgcttcaataatattg
aaaaaggaagagtatgagtattcaacatttccgtgtcgcccttattcccttttttgcggcattttgcc
ttcctgtttttgctcacccagaaacgctggtgaaagtaaaagatgctgaagatcagttgggtgcacga
gtgggttacatcgaactggatctcaacagcggtaagatccttgagagttttcgccccgaagaacgttt
tccaatgatgagcacttttaaagttctgctatgtggcgcggtattatcccgtattgacgccgggcaag
agcaactcggtcgccgcatacactattctcagaatgacttggttgagtactcaccagtcacagaaaag
catcttacggatggcatgacagtaagagaattatgcagtgctgccataaccatgagtgataacactgc
ggccaacttacttctgacaacgatcggaggaccgaaggagctaaccgcttttttgcacaacatgggg

atcatgtaactcgccttgatcgttgggaaccggagctgaatgaagccataccaaacgacgagcgtgac
accacgatgcctgtagcaatggcaacaacgttgcgcaaactattaactggcgaactacttactctagc
ttcccggcaacaattgatagactggatggaggcggataaagttgcaggaccacttctgcgctcggccc
ttccggctggctggtttattgctgataaatctggagccggtgagcgtggctctcgcggtatcattgca
gcactggggccagatggtaagccctcccgtatcgtagttatctacacgacggggagtcaggcaactat
ggatgaacgaaatagacagatcgctgagataggtgcctcactgattaagcattggtaggaattaatga
tgtctcgtttagataaaagtaaagtgattaacagcgcattagagctgcttaatgaggtcggaatcgaa
ggtttaacaacccgtaaactcgcccagaagctaggtgtagagcagcctacattgtattggcatgtaaa
aaataagcgggctttgctcgacgccttagccattgagatgttagataggcaccatactcacttttgcc
ctttagaaggggaaagctggcaagatttttacgtaataacgctaaaagttttagatgtgctttacta
agtcatcgcgatggagcaaaagtacatttaggtacacggcctacagaaaaacagtatgaaactctcga
aaatcaattagcctttttatgccaacaaggtttttcactagagaatgcattatatgcactcagcgcag
tggggcattttactttaggttgcgtattggaagatcaagagcatcaagtcgctaaagaagaaaggaa
acacctactactgatagtatgccgccattattacgacaagctatcgaattatttgatcaccaaggtgc
agagccagccttcttattcggccttgaattgatcatatgcggattagaaaaacaacttaaatgtgaaa
gtgggtcttaaaagcagcataacctttttccgtgatggtaacttcactagtttaaaaggatctaggtg
aagatcctttttgataatctcatgaccaaaatcccttaacgtgagttttcgttccactgagcgtcaga
ccccgtagaaaagatcaaaggatcttcttgagatcctttttttctgcgcgtaatctgctgcttgcaaa
caaaaaaaccaccgctaccagcggtggtttgtttgccggatcaagagctaccaactctttttccgaag
gtaactggcttcagcagagcgcagataccaaatactgtccttctagtgtagccgtagttaggccacca
cttcaagaactctgtagcaccgcctacatacctcgctctgctaatcctgttaccagtggctgctgcca
gtggcgataagtcgtgtcttaccgggttggactcaagacgatagttaccggataaggcgcagcggtcg
ggctgaacggggggttcgtgcacacagcccagcttggagcgaacgacctacaccgaactgagatacct
acagcgtgagctatgagaaagcgccacgcttcccgaagggagaaaggcggacaggtatccggtaagcg
gcagggtcggaacaggagagcgcacgagggagcttccagggggaaacgcctggtatctttatagtcct
gtcgggtttcgccacctctgacttgagcgtcgatttttgtgatgctcgtcaggggggcggagcctatg
gaaaaacgccagcaacgcggcctttttacggttcctggccttttgctggccttttgctcacatgaccc
gacaccatcgaatggccagat

>pASK-IBA5+_6xHis-*Pk*DLH
gattaattcctaatttttgttgacactctatcattgatagagttattttaccactccctatcagtgat
agagaaaagtgaaatgaatagttcgacaaaaatctagaaataattttgtttaactttaagaaggagat
atacaaATGCATCACCATCATCACCACGGTGGAAGTCTGACCGAAGGTATTAGCATTCAGAGCTATGA
TGGTCATACCTTTGGTGCACTGGTTGGTAGTCCGGCAAAAGCACCGGCACCGGTTATTGTTATTGCAC
AAGAAATTTTTGGCGTGAACGCCTTTATGCGTGAAACCGTTAGCTGGCTGGTTGATCAGGGTTATGCA
GCAGTTTGTCCGGATCTGTATGCACGTCAGGCACCGGGTACAGCACTGGATCCGCAGGATGAACGTCA
GCGTGAACAGGCATATAAACTGTGGCAGGCATTTGATATGGAAGCCGGTGTTGGTGATCTGGAAGCAG

CAATTCGTTATGCCCGTCATCAGCCGTATAGCAATGGTAAAGTTGGTCTGGTTGGTTATTGCTTAGGT
GGTGCCCTGGCATTTCTGGTTGCAGCAAAAGGTTATGTTGATCGTGCAGTGGGTTATTATGGTGTTGG
CCTGGAAAAACAGCTGAATAAAGTTCCGGAAGTTAAACATCCGGCACTGTTTCATATGGGTGGTCAGG
ATCATTTTGTTCCGGCACCGAGCCGTCAGCTGATTACCGAAGGTTTTGGTGCAAATCCGCTGCTGCAG
GTTCATTGGTATGAAGAAGCAGGTCATAGCTTTGCACGTACCAGCAGCAGTGGTTATGTGGCAAGCGC
AGCAGCACTGGCAAATGAACGTACCCTGGATTTTCTGGCACCGCTGCAGAGCAAAAAACCGTAAtgat
atctaactaagcttgacctgtgaagtgaaaaatggcgcacattgtgcgacatttttttgtctgccgt
ttaccgctactgcgtcacggatctccacgcgccctgtagcggcgcattaagcgcggcgggtgtggtgg
ttacgcgcagcgtgaccgctacacttgccagcgccctagcgcccgctcctttcgctttcttcccttcc
tttctcgccacgttcgccggctttccccgtcaagctctaaatcgggggctccctttagggttccgatt
tagtgctttacggcacctcgaccccaaaaaacttgattagggtgatggttcacgtagtgggccatcgc
cctgatagacggtttttcgccctttgacgttggagtccacgttctttaatagtggactcttgttccaa
actggaacaacactcaaccctatctcggtctattcttttgatttataagggattttgccgatttcggc
ctattggttaaaaaatgagctgatttaacaaaaatttaacgcgaattttaacaaaatattaacgctta
caatttcaggtggcacttttcggggaaatgtgcgcggaacccctatttgtttatttttctaaatacat
tcaaatatgtatccgctcatgagacaataaccctgataaatgcttcaataatattgaaaaaggaagag
tatgagtattcaacatttccgtgtcgcccttattcccttttttgcggcattttgccttcctgtttttg
ctcacccagaaacgctggtgaaagtaaaagatgctgaagatcagttgggtgcacgagtgggttacatc
gaactggatctcaacagcggtaagatccttgagagttttcgccccgaagaacgttttccaatgatgag
cacttttaaagttctgctatgtggcgcggtattatcccgtattgacgccgggcaagagcaactcggtc
gccgcatacactattctcagaatgacttggttgagtactcaccagtcacagaaaagcatcttacggat
ggcatgacagtaagagaattatgcagtgctgccataaccatgagtgataacactgcggccaacttact
tctgacaacgatcggaggaccgaaggagctaaccgcttttttgcacaacatgggggatcatgtaactc
gccttgatcgttgggaaccggagctgaatgaagccataccaaacgacgagcgtgacaccacgatgcct
gtagcaatggcaacaacgttgcgcaaactattaactggcgaactacttactctagcttcccggcaaca
attgatagactggatggaggcggataaagttgcaggaccacttctgcgctcggcccttccggctggct
ggtttattgctgataaatctggagccggtgagcgtggctctcgcggtatcattgcagcactggggcca
gatggtaagccctcccgtatcgtagttatctacacgacggggagtcaggcaactatggatgaacgaaa
tagacagatcgctgagataggtgcctcactgattaagcattggtaggaattaatgatgtctcgtttag
ataaaagtaaagtgattaacagcgcattagagctgcttaatgaggtcggaatcgaaggtttaacaacc
cgtaaactcgcccagaagctaggtgtagagcagcctacattgtattggcatgtaaaaaataagcgggc
tttgctcgacgccttagccattgagatgttagataggcaccatactcacttttgccctttagaagggg
aaagctggcaagatttttttacgtaataacgctaaaagttttagatgtgctttactaagtcatcgcgat
ggagcaaaagtacatttaggtacacggcctacagaaaaacagtatgaaactctcgaaaatcaattagc
cttttatgccaacaaggtttttcactagagaatgcattatatgcactcagcgcagtggggcatttta
ctttaggttgcgtattggaagatcaagagcatcaagtcgctaaagaagaaagggaaacacctactact
gatagtatgccgccattattacgacaagctatcgaattatttgatcaccaaggtgcagagccagcctt

cttattcggccttgaattgatcatatgcggattagaaaaacaacttaaatgtgaaagtgggtcttaaa
agcagcataacctttttccgtgatggtaacttcactagtttaaaaggatctaggtgaagatcctttttt
gataatctcatgaccaaaatcccttaacgtgagttttcgttccactgagcgtcagaccccgtagaaaa
gatcaaaggatcttcttgagatcctttttttctgcgcgtaatctgctgcttgcaaacaaaaaaaccac
cgctaccagcggtggtttgtttgccggatcaagagctaccaactctttttccgaaggtaactggcttc
agcagagcgcagataccaaatactgtccttctagtgtagccgtagttaggccaccacttcaagaactc
tgtagcaccgcctacatacctcgctctgctaatcctgttaccagtggctgctgccagtggcgataagt
cgtgtcttaccgggttggactcaagacgatagttaccggataaggcgcagcggtcgggctgaacgggg
ggttcgtgcacacagcccagcttggagcgaacgacctacaccgaactgagatacctacagcgtgagct
atgagaaagcgccacgcttcccgaagggagaaaggcggacaggtatccggtaagcggcagggtcggaa
caggagagcgcacgagggagcttccagggggaaacgcctggtatctttatagtcctgtcgggtttcgc
cacctctgacttgagcgtcgatttttgtgatgctcgtcaggggggcggagcctatggaaaaacgccag
caacgcggcctttttacggttcctggccttttgctggccttttgctcacatgacccgacaccatcgaa
tggccagat
